# Somatic epimutations cap genetic determinism in the human diploid chromatin epigenome

**DOI:** 10.1101/2024.06.14.599122

**Authors:** Mitchell R. Vollger, Elliott G. Swanson, Shane J. Neph, Benjamin J. Mallory, Danilo Dubocanin, Ching-Huang Ho, Y. H. Hank Cheng, Jane Ranchalis, Katherine M. Munson, Adriana E. Sedeño-Cortés, Devaraja G. Mudeppa, William E. Fondrie, Stephanie C. Bohaczuk, Maxwell A. Dippel, Yizi Mao, Nancy L. Parmalee, William T. Harvey, Youngjun Kwon, Gage H. Garcia, Kendra Hoekzema, Jeffrey G. Meyer, Mine Cicek, Evan E. Eichler, William S. Noble, Daniela M. Witten, James T. Bennett, John P. Ray, Andrew B. Stergachis

## Abstract

Diploid human cells contain two non-identical genomes, and differences in their regulation underlie human development and disease. We present Fiber-seq Inferred Regulatory Elements (FIRE) and show that FIRE provides a more comprehensive and quantitative snapshot of the accessible chromatin landscape across the 6 Gbp diploid human genome, overcoming previously unrecognized biases in existing regulatory element catalogs. FIRE enables comprehensive detection of haplotype-selective chromatin accessibility (HSCA), exposing novel imprinted elements lacking underlying parent-of-origin CpG methylation differences, and gene regulatory modules that permit genes to escape X chromosome inactivation. We uncover that the human leukocyte antigen (HLA) locus harbors the most HSCA in immune cells, where we resolve specific transcription factor (TF) binding events disrupted by disease-associated variants. Finally, we demonstrate that the regulatory landscape of a cell is littered with autosomal somatic chromatin epimutations that are propagated by clonal expansions to create mitotically stable and non-genetically deterministic chromatin alterations.

## Introduction

Advances in long-read sequencing (*1*, *2*) have enabled the routine *de novo* assembly of 6 Gbp diploid human genomes at reference quality (*3*, *4*), resulting in the completion of the first human genome (*5–7*) and pangenome (*8–10*). These advances have improved our understanding of the structure of the human genome and human genetic variation, enabling researchers to investigate genetic variants within their native haplotype context along the 6 Gbp diploid human genome. Now, our next challenge is studying the function of this 6 Gbp genome, as this can shed light on how non-coding genetic variants contribute to human disease risk, uncover somatic epimutations that selectively arise on only a single haplotype, and illuminate gene regulatory patterns in genetically complex regions of the genome that show the greatest sequence diversity between humans.

Despite the potential for studying the function of genetic variation along the diploid genome, most chromatin assays are reliant on collapsed 3 Gbp representations of a human genome. Specifically, given the inconsistent density of genetic variation within the human genome, short-read-based chromatin methods are poorly suited for uniquely phasing chromatin across the diploid genome, and even when an element happens to overlap a heterozygous single-nucleotide variant (SNV), read-mapping artifacts are known to confound measures of allele-specific chromatin accessibility.

Long-read single-molecule chromatin assays (*11–14*) have overcome these challenges by leveraging the ability of long reads to correctly map to the diploid genome (*12*, *15*, *16*). For example, Fiber-seq uses a non-specific *N^6^*-adenine methyltransferase (m6A-MTase) to stencil protein occupancy footprints along DNA molecules in the form of m6A-modified bases (**Fig. 1A**) (*11*). These m6A-modified DNA molecules are sequenced using a single-molecule platform (*17*), enabling the synchronous readout of the genetic sequence, CpG methylation status, and chromatin architecture of each multi-kilobase sequencing read. Prior work has used this data to accurately identify the presence of transcription factors (*18*); however, despite the promise of single-molecule chromatin assays, current approaches for analyzing this new data type are sensitive to potential experimental batch effects, and are unable to identify putative regulatory elements *de novo*, resulting in a dearth of new knowledge about how gene regulation is structured across the *entire* 6 Gbp genome, and how this structure changes across different tissues and disease states. To realize the full potential of long-read single-molecule chromatin assays, we developed a robust machine learning classifier that enables the precise delineation of chromatin *architectures* across the diploid genome with single-molecule and single-haplotype precision at near-nucleotide resolution. Furthermore, we leverage this approach across multiple cohorts to uncover the basic principles guiding diploid human gene regulation, including the relative contributions of rare and common genetic variants, imprinting, somatic epimutations, and X chromosome inactivation.

**Figure 1.**
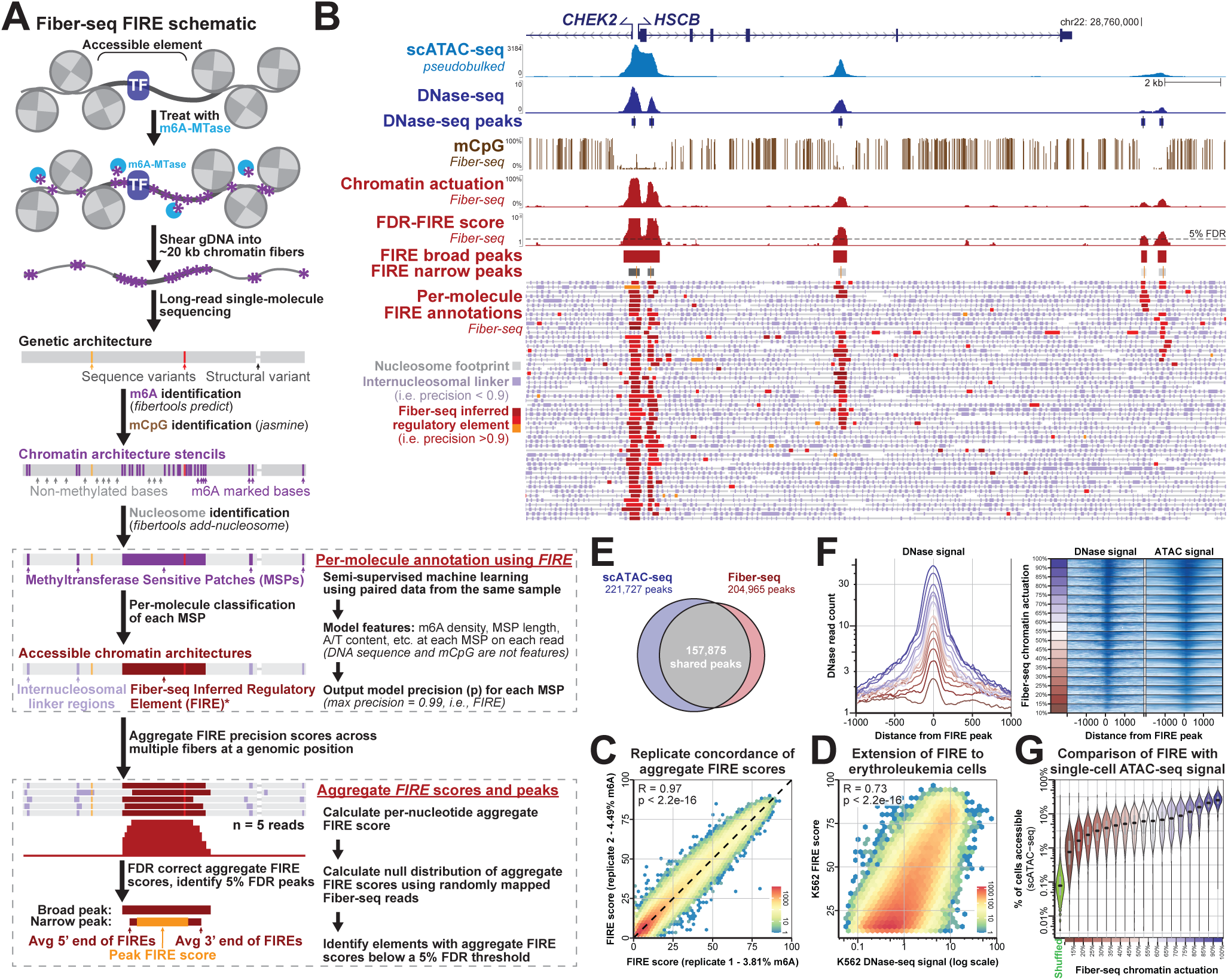
Fiber-seq Inferred Regulatory Elements and benchmarking against existing chromatin accessibility measures. **A)** A schematic of Fiber-seq experimental and computational processing, including the identification of Fiber-seq Inferred Regulatory Elements (FIREs). **B)** Genomic locus comparing the relationship between scATAC–seq, DNase–seq, mCpG, FIRE percent chromatin actuation, and FIRE peaks in GM12878. Below are individual Fiber-seq reads with MTase Sensitive Patches (MSPs) marked in purple, nucleosomes marked in gray, and FIRE elements marked in red. White regions separate individual reads. **C)** Correlation of FIRE score within the peaks of two technical replicates of COLO829BL (two-sided *t*-test). **D)** Correlation of FIRE score with bulk DNase-seq in K562 accessible peaks (two-sided *t*-test). **E)** Venn diagram showing the overlap of FIRE and scATAC peaks in GM12878. **F)** (Left) Average count of DNase I reads over FIRE peaks binned by their percent actuation (red-blue color scale). (Right) Percentile normalized scATAC and DNase I signal for 100 random FIRE peaks across each percent actuation bin. **G)** Comparison of percent actuation quantified by Fiber-seq and scATAC-seq. scATAC-seq accessibility values represent the fraction of single cells with at least one sequenced fragment overlapping the respective peak. FIRE peaks are binned by Fiber-seq percent actuation (left) and scATAC-seq percent actuation (right).

## Results

### Single-molecule Fiber-seq inferred regulatory elements

To generate a haplotype-resolved map of human gene regulation, we first sought to develop a machine learning classifier for Fiber-seq that could accurately classify m6A-modified regions (*i.e.,* MTase Sensitive Patches [MSPs]) as actuated single-molecule regulatory elements (**Fig. 1A**). Importantly, accurately measuring chromatin actuation *de novo* at the level of single fibers permits the quantitative measurement of chromatin accessibility at a regulatory element as the percent of molecules with chromatin actuation, a potentially more intuitive and quantitative metric of chromatin than is currently possible using traditional chromatin assays. Although existing single-molecule stenciling methods have used MSP size as a simple single-molecule classifier for this process (*11*, *12*), this simple classifier has relatively poor performance for the *de novo* identification of putative regulatory elements genome-wide (**Fig. S1D,E**), likely owing to its inability to disentangle stochastic, unstable nucleosome occupancy events from actuated regulatory elements. Furthermore, we have found that MSP size is highly sensitive to technical sample-to-sample variation in the total rate of adenine methylation (**Fig. S1A,B**), which can result in systematic biases in identifying accessible chromatin using single-molecule stenciling methods (*19*) (**Fig. S1C**). To resolve this challenge limiting the broader adoption of single-molecule chromatin stenciling methods, we develop a tool for the accurate *de novo* classification of putative regulatory elements using only Fiber-seq data that does not need any prior knowledge of the underlying reference genome, or the exact primary sequence of the underlying fiber. Importantly, this structure enables this tool to generalize across different cell types that employ unique sets of binding elements to regulate their accessible chromatin (*15*, *16*, *20*, *21*), complex genomic regions not present within the reference GRCh38 (*22*, *23*), as well as organisms that have completely distinct genomic architectures (*24*).

To create our training data, we generated multiple Fiber-seq datasets from the reference cell line GM12878 (totaling 136-fold coverage) that differed by over 2.3-fold in their global m6A methylation rate, enabling the final model to extend to analytical fluctuations in the Fiber-seq method (**Fig. S1A-C**). Although our final tool works irrespective of a reference genome, labels for the training data relied on GRCh38, as this enabled us to use the extensive short-read epigenetic data from GM12878 to assign positive and negative labels to individual MSPs. Specifically, MSPs that overlapped GM12878 DNase-seq DHSs or GM12878 CTCF ChIP-seq peaks were assigned a positive label. In contrast, MSPs from short-read mappable regions of the genome without these short-read epigenetic peak annotations were assigned a negative label (**Fig. S2, Supplemental Table 1**). Notably, regulatory elements identified using bulk methods can exhibit marked heterogeneity in their single-molecule actuation pattern (*i.e.,* can be an actuated element, or merely an internucleosomal linker region at the single-molecule level) (*11*, *12*, *14*) (**Fig. S3**). As such, MSPs ascribed a positive label based on their genomic location relative to bulk chromatin measurements likely represent a mixture of MSPs arising from actuated regulatory elements and others arising from internucleosomal linker regions. Consequently, our training data is best described as mixed-positive labels and clean-negative labels, a training paradigm best approached using a semi-supervised machine learning framework (*25–27*). To implement a semi-supervised approach, we trained an XGBoost model with five-fold cross-validation (*28*), iteratively retraining the model using the learned prediction from the previous iteration’s model to create positive labels at 95% estimated precision until the number of positive identifications at 95% estimated precision (*Methods,* **Supplemental Note**, **Fig. S2A-C**) in the validation set ceased to increase (15 iterations). We applied this final model to our held-out test data to create an estimated precision based on the model’s prediction score (*Methods*). We then used the estimated precision to classify 1,959,888,668 MSPs across 24,771,424 chromatin Fiber-seq reads as either **<u>F</u>**iber-seq **I**nferred **<u>R</u>**egulatory **<u>E</u>**lements (FIREs; 95% estimated precision or greater; n=32,006,894) or internucleosomal linker regions (less than 95% estimated precision). Consistent with the model training, MSPs classified as FIREs were markedly enriched in DHSs compared to using MSP length alone as a classifier (**Fig. S1D**).

### Aggregating across Fiber-seq inferred regulatory elements

As no other methods exist for classifying per-molecule chromatin architectures, to appropriately benchmark our FIRE classifications, we next needed to develop a method for performing the *de novo* identification of putative regulatory elements genome-wide using these FIREs (**Fig. 1B**). To accomplish this, we first created a coverage-normalized aggregate FIRE score (*Methods*) that combines the FIRE classification across multiple molecules for every base in the genome. We then calibrated these aggregate FIRE scores using a null aggregate FIRE score distribution, enabling us to calculate a false discovery rate (FDR, *Methods*, **Fig. S4A,B**) and identify genomic positions that met a 5% FDR threshold. *De novo* Fiber-seq accessible chromatin peaks were directly identified based on local maximums that met the 5% FDR threshold, and the bounds of each peak were defined based on the median start and end positions of the underlying single-molecule FIRE elements (**Fig. 1A,B**). Finally, at each peak, we calculated the percentage of all Fiber-seq reads with a FIRE element that overlapped the local maximum of the peak, hereafter referred to as the “percent actuation.” In total, we identified 204,965 FIRE peaks in GM12878, which were robust to underlying changes in our model (**Fig. S5**), and overlapped 89% of the previously annotated GM12878 DNaseI hypersensitive sites (**Supplemental Table 2**). Notably, this approach provides an intuitive, biologically meaningful metric of chromatin accessibility and peak width, with the boundaries of each FIRE peak corresponding to the nucleotide-precise median start/stop positions of the underlying single-molecule data, and the intensity reflecting the exact fraction of chromatin fibers with accessibility.

### Benchmarking FIRE single-molecule measures of chromatin actuation

To test the generalizability of our approach, we generated two biological replicates of Fiber-seq from a different human cell line (COLO829BL) (totaling 319-fold coverage), which demonstrated that we can readily extend our model to other samples and that FIRE scores are highly concordant between replicates (*R*=0.97) (**Fig. 1C**) even in the setting of experimental differences in the overall methylation rate between replicates (3.81% vs. 4.49% in replicate 1 versus replicate 2, respectively). Furthermore, we showed that FIRE scores were stable after down-sampling our GM12878 data from 135- to 30-fold coverage (**Fig. S4C,D**). In addition, we generated Fiber-seq data from different cell types (K562, A549, Jurkat, and THP-1 cells) and demonstrated that overall FIRE measures of chromatin actuation are strongly correlated with those obtained using DNase-seq (**Figs. 1D, S6A,B, S7A-C**). We also benchmarked our FIRE peaks and chromatin actuation measures against single-cell ATAC-seq (scATAC-seq) data, as this data type was not used in our training. Specifically, we pseudo-bulked an entire 26,910-cell ENCODE GM12878 scATAC-seq dataset and then calculated peaks using MACS2. Overall, we found that peaks of accessibility identified using Fiber-seq and scATAC-seq showed substantial concordance, with 77% of all FIRE peaks also called by scATAC-seq (**Fig. 1e; Fig. S6A,B**). Furthermore, we found scATAC-seq signal support for even the least actuated Fiber-seq peaks (10-15%) (**Fig. 1F**) and that scATAC signal increases at peaks with higher Fiber-seq percent actuation (**Fig. 1F,G**), indicating that overall Fiber-seq percent actuation scores are in agreement with short-read bulk measures of chromatin accessibility.

In total, 22.2% of the FIRE peaks were unique to FIRE and not called using either the scATAC-seq or DNase-seq data (*i.e.,* FIRE-specific peaks). These FIRE-specific peaks were on average narrower in terms of their chromatin accessibility (**Fig. 2A,B**), did not show major changes in GC content (**Figs. S7D and S8**), and were significantly enriched in REST ChIP-seq peaks that demonstrated single-molecule TF occupancy patterns at these REST sites (**Fig. S7E-F**). To validate the biological relevance of these FIRE-specific peaks, we overlapped all FIRE peaks with genetic variants in GM12878 cells that show genome-wide significant associations (GWAS) with different human traits and diseases (*29*). We observed that FIRE peaks were significantly enriched for overlapping disease-associated GWAS variants, as expected (p-value = 8.40e-71, two-sided Fisher’s exact test) (*30*). However, unlike scATAC-seq-specific or DNase-specific peaks, FIRE-specific peaks also showed a comparable enrichment for disease-associated GWAS variants (**Fig. 2C**, p-value = 1.32e-17, two-sided Fisher’s exact test), showing that these Fiber-seq unique elements are of comparably high quality as Fiber-seq peaks that have ATAC-seq and DNase-seq support. Overall, these findings demonstrate that FIRE enables the accurate and robust *de novo* identification of accessible chromatin elements solely using long-read epigenomic data.

**Figure 2.**
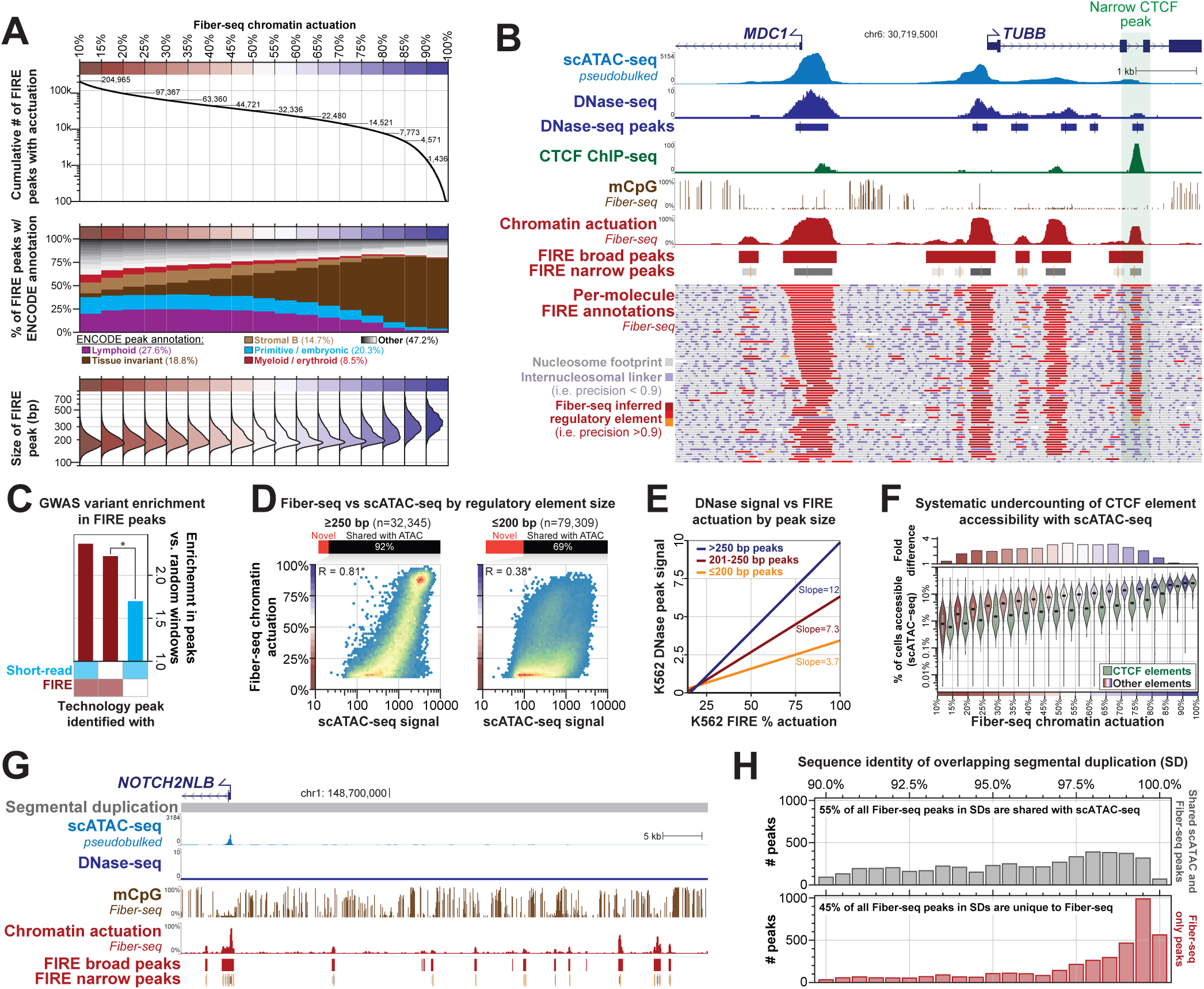
Chromatin features within FIRE-specific peaks. **A)** Features of FIRE peaks binned by percent actuation. **B)** Genomic locus comparing the relationship between scATAC-seq, DNase–seq, mCpG, CTCF ChIP-seq, FIRE percent chromatin actuation, and FIRE peaks in GM12878. A representative CTCF site with greater accessibility in Fiber-seq data than scATAC-seq and DNase–seq is highlighted in green (right). **C)** Per-base enrichment of GWAS variants within shared peaks between FIRE and DNase/scATAC, FIRE only peaks, and peaks unique to DNase or scATAC as compared to shuffled random windows of the same size (p-value = 1.32e-17, two-sided Fisher’s exact test). **D)** Correlation of FIRE percent actuation and scATAC-seq signal within FIRE peaks faceted by FIRE peak size (Pearson’s correlation; p < 2.2e−16 two-sided t-test). **E)** Prediction of percent FIRE actuation from DNase peak as signal using a linear model for different bins of FIRE peak size. **F)** Comparison of scATAC-seq signal to FIRE in peaks with (green) and without (red-blue) CTCF ChIP-seq peaks. **G)** Genomic locus of *NOTCH2NLB* comparing the ability to map into repetitive regions between scATAC-seq, DNase–seq, and Fiber-seq. **H)** FIRE peaks within segmental duplications stratified by the sequence identity of the underlying segmental duplication. FIRE peaks with a shared scATAC-seq peak are colored in gray, and peaks unique to FIRE are colored in red.

### FIRE provides more accurate measures of chromatin accessibility

We next investigated differences between FIRE measures of chromatin accessibility and those of existing short-read chromatin assays. First, we observed that controlling for GC content improved the correlation between short-read chromatin assays and Fiber-seq, likely reflecting GC biases that are induced during the PCR amplification steps of these short-read assays (**Fig. S4E**). Second, we observed that 78.4% of the GM12878 peaks identified only in Fiber-seq were <200 bp in length, raising the possibility that short-read chromatin assays may be selectively biased against detecting accessibility at shorter regulatory elements (**Fig. 2d, S7D**). Consistent with this, we observed that although short-read measures of chromatin accessibility are strongly associated with Fiber-seq percent actuation for elements >250 bp in length, this association markedly deteriorates for elements <200 bp in length (**Fig. 2D,E**), which are enriched in CTCF ChIP-seq peaks (26.5% of all peaks ≤200 bp). This potential size-dependent bias also appeared to extend to scATAC-seq measures of chromatin accessibility. Specifically, elements detected in 10-15% of cells as measured by scATAC-seq can have Fiber-seq actuation ranging from <10% to >90% of fibers (**Fig. 1F**), with scATAC-seq signal being 3.47-fold lower at CTCF ChIP-seq peaks compared to other scATAC-peaks with a similar Fiber-seq percent actuation (**Fig. 2F**). These findings suggest that existing short-read maps of chromatin accessibility are reporting inaccurate chromatin accessibility measurements at CTCF binding elements, and consistent with this, we observed that FIRE actuation at a CTCF ChIP-seq peak was markedly better at predicting CTCF ChIP-seq signal at that site than ATAC-seq signal (*R^2^* of 0.41 vs. 0.17, respectively) (**Fig. S9A**). Overall, this size-dependent bias resulted in the width of a peak being more predictive of ATAC-seq Tn5 insertions at a site than the actual number of chromatin fibers on which that peak is actuated (**Fig. 2F, S7C**). Although the exact cause of this phenomenon is unknown, it is likely driven by a combination of artifacts inherent to short-read chromatin methods, including: (1) available template region for Tn5 transposition/DNaseI nicking; (2) fragment size selection; (3) PCR amplification preferences; and (4) short-read sequencing biases.

### FIRE provides a more complete genome-wide map of chromatin accessibility

We found that 8.66% (n=3,933) of the 45,420 GM12878 peaks identified only in Fiber-seq mapped to segmentally duplicated (SD) regions of the human genome (**Fig. 2G,H**). SDs comprise ∼200 Mbp of genomic sequence (*31*, *32*) that are known to contribute to a variety of human diseases (*33*, *34*), but these regions have been challenging to study owing to mapping issues of short-read chromatin assays to these highly similar duplicated sequences (*35*). Overall, 45% (3,933/8,700) of all Fiber-seq FIRE peaks in SDs are unique to FIRE, and the SDs with the highest sequence identity contained the highest fraction of FIRE unique peaks (**Fig. 2H**). Together, this demonstrates that not only can FIRE better resolve chromatin architectures within the traditionally mappable portion of the genome, but it is also able to uniquely resolve chromatin patterns across complex genomic regions that are largely impenetrable to short-read chromatin assays.

### A haplotype-resolved view of human gene regulation

We next assessed whether haplotype-phasing our Fiber-seq reads could enable us to map chromatin across the 6 Gbp diploid human genome. For GM12878, each ∼20kb read on average spans at least one heterozygous variant, and parental short-read sequencing data can bin these reads based on their maternal or paternal origin, enabling the accurate haplotype phasing of 87.9% of reads across GRCh38 (*Methods*). Using these haplotype-phased reads, we pseudo-bulked our per-read chromatin architectures by haplotype, generating haplotype-specific chromatin actuation, CpG methylation, nucleosome positioning, and TF occupancy patterns genome-wide (**Fig. 3A**). Notably, we observed that elements showed more variability in actuation between haplotypes than between Fiber-seq replicates (**Fig. S9B**), highlighting the reproducibility of FIRE results between replicates and the central role of individual haplotypes in guiding chromatin accessibility patterns. We identified elements with haplotype-selective chromatin accessibility (HSCA) by comparing the percent actuation between the two haplotypes using a Fisher’s exact test with a genome-wide FDR correction (Benjamini-Hochberg). This resulted in the identification of 9,773 elements with nominally significant (p-value <0.05) HSCAs in GM12878 cells, with 1,231 of these elements meeting genome-wide significance along the autosomes (FDR 5%) (**Fig. 3A,B**). As expected, haplotype-selective peaks were enriched at known imprinted sites, such as the *GNAS* locus, enabling the precise demarcation of actuated elements and TF binding events that are impacted by imprinting at these sites (**Fig. 3A**). However, known imprinting sites comprised only 5% of all haplotype-selective peaks (**Fig. 3C**), indicating that other features are the major drivers of haplotype-selective chromatin genome-wide.

**Figure 3.**
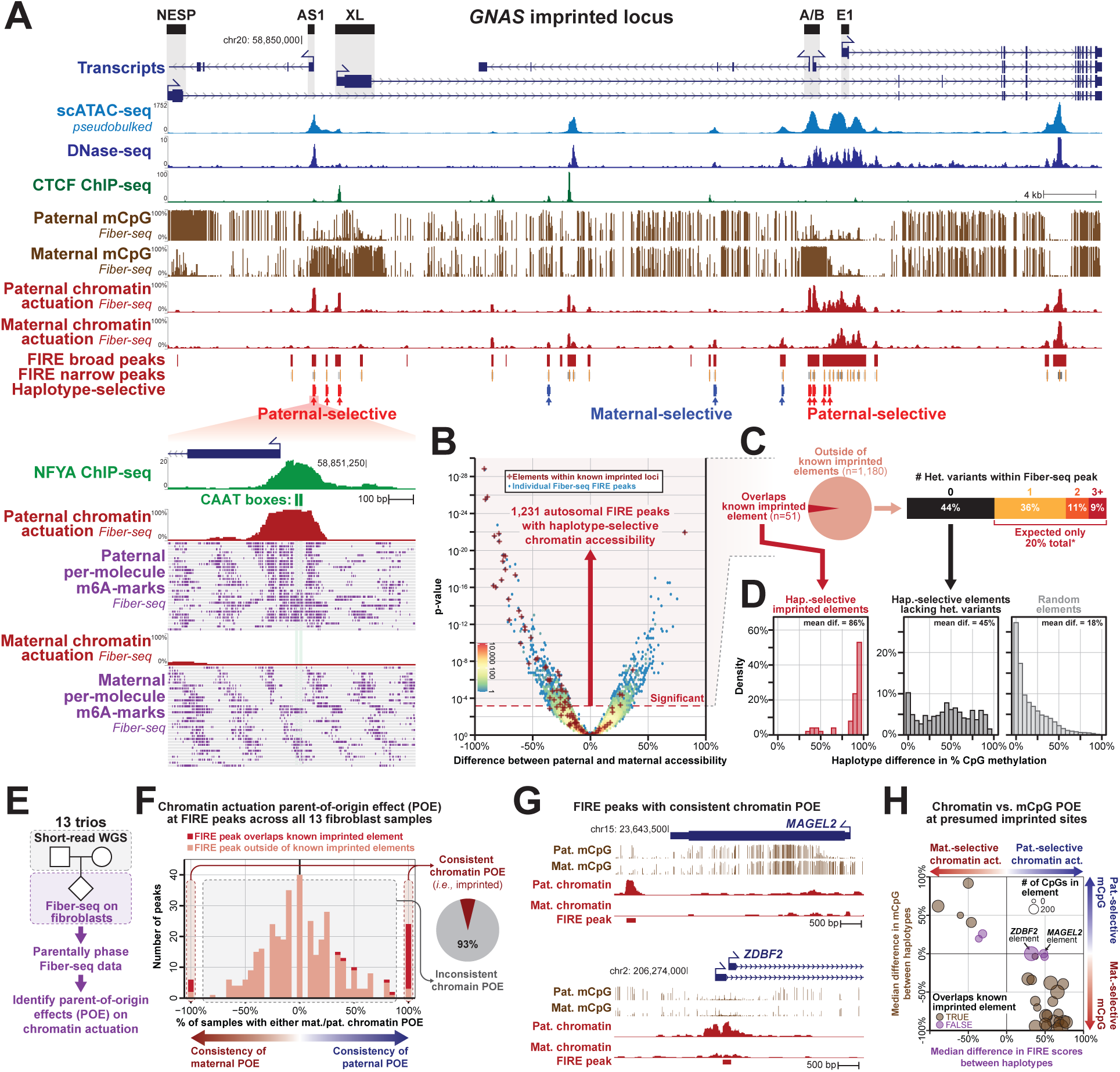
Haplotype-selective chromatin accessibility. **A)** The *GNAS* imprinted locus comparing the relationship between *GNAS* isoforms, scATAC-seq, DNase–seq, CTCF ChIP-seq, NFYA ChIP-seq, mCpG, FIRE percent actuation, FIRE peaks, and maternally (blue) or paternally (red) haplotype-selective FIRE peaks in GM12878. Fiber-seq captures haplotype-selective chromatin architectures in both mCpG and chromatin actuation. **B)** Difference between maternal and paternal accessibility for FIRE haplotype-selective peaks stratified by p-value (two-sided Fisher’s exact test). The dashed line indicates genome-wide significance after applying a 5% FDR correction (Benjamini-Hochberg), and the red plus marks indicate known imprinted sites. **C)** Stratification of haplotype-selective peaks by imprinting status and the number of genetic variants within each peak. **D)** Histogram of the haplotype differences in percent CpG methylation for haplotype-selective peaks within imprinted sites, sites without genetic variants, and non-haplotype-selective peaks. **E)** Schematic of sequencing 13 trios with parental short-reads for phasing and Fiber-seq on the probands to identify parent of origin effects (POE) in chromatin. **F)** Distribution of haplotype-selective chromatin accessibility (HSCA) peaks showing the fraction of fibroblast samples with consistent maternal or paternal bias. Dark red in the histogram indicates previously identified imprinted sites; the pie chart shows the proportion of sites with consistent POE. **G)** Browser views of two genomic regions (*MAGEL2* and *ZDBF2*) demonstrating consistent POE in fibroblasts. **H)** Relationship between parental bias in CpG methylation (y-axis) and FIRE actuation (x-axis). Purple points represent new imprinted sites, with three showing consistent POE in FIRE without evidence of differential CpG methylation.

To further validate this, we performed Fiber-seq on 13 fibroblast cell lines for which parental genomic sequencing data was available to phase chromatin patterns to the maternal and paternal haplotype (**Fig. 3E**). Using this, we identified 30 FIRE peaks that showed consistent parent-of-origin haplotype-selective chromatin across the 13 fibroblast cell lines (**Fig. 3F**), 5 of which were previously unannotated as being within imprinted elements as defined using CpG methylation patterns. Notably, all 5 of these elements are within genomic regions known to be imprinted but do not directly overlap a differentially methylated CpG region (**Fig. 3G,H**), indicating that parent-of-origin chromatin accessibility at these sites is independent of the local CpG methylation. Overall, this demonstrates that only a fraction of the haplotype-selective peaks in a cell are caused by previously uncharacterized imprinted elements, and that elements with consistent parent-of-origin chromatin accessibility can be maintained even in the absence of underlying parent-of-origin CpG methylation.

### Haplotype-selective chromatin marks disease-associated loci and elements

GM12878 cells contain 3.2 million autosomal heterozygous variants that distinguish each haplotype, and we next sought to evaluate which of these genetic variants contribute to the formation of haplotype-selective chromatin in GM12878 cells. Overall, HSCA elements are enriched in directly overlapping heterozygous genetic variants, with 56% of all autosomal elements with HSCA containing at least one heterozygous genetic variant (**Fig. 3C**), and some containing over 10 heterozygous genetic variants. To determine whether the encompassed heterozygous variants may in fact be causal of the haplotype-selective chromatin signal, we quantified the overlap of these elements with variants known to have genome-wide significant associations with different human traits and diseases (*i.e.,* lead GWAS variants). Overall, elements with HSCA showed a 6-fold enrichment (36 versus 6) in overlapping lead GWAS variants compared to random non-HSCA FIRE peaks (**Fig. 4A**), indicating that haplotype-selective chromatin at these elements may, in fact, be directly resulting from the effect of these underlying genetic variants.

**Figure 4.**
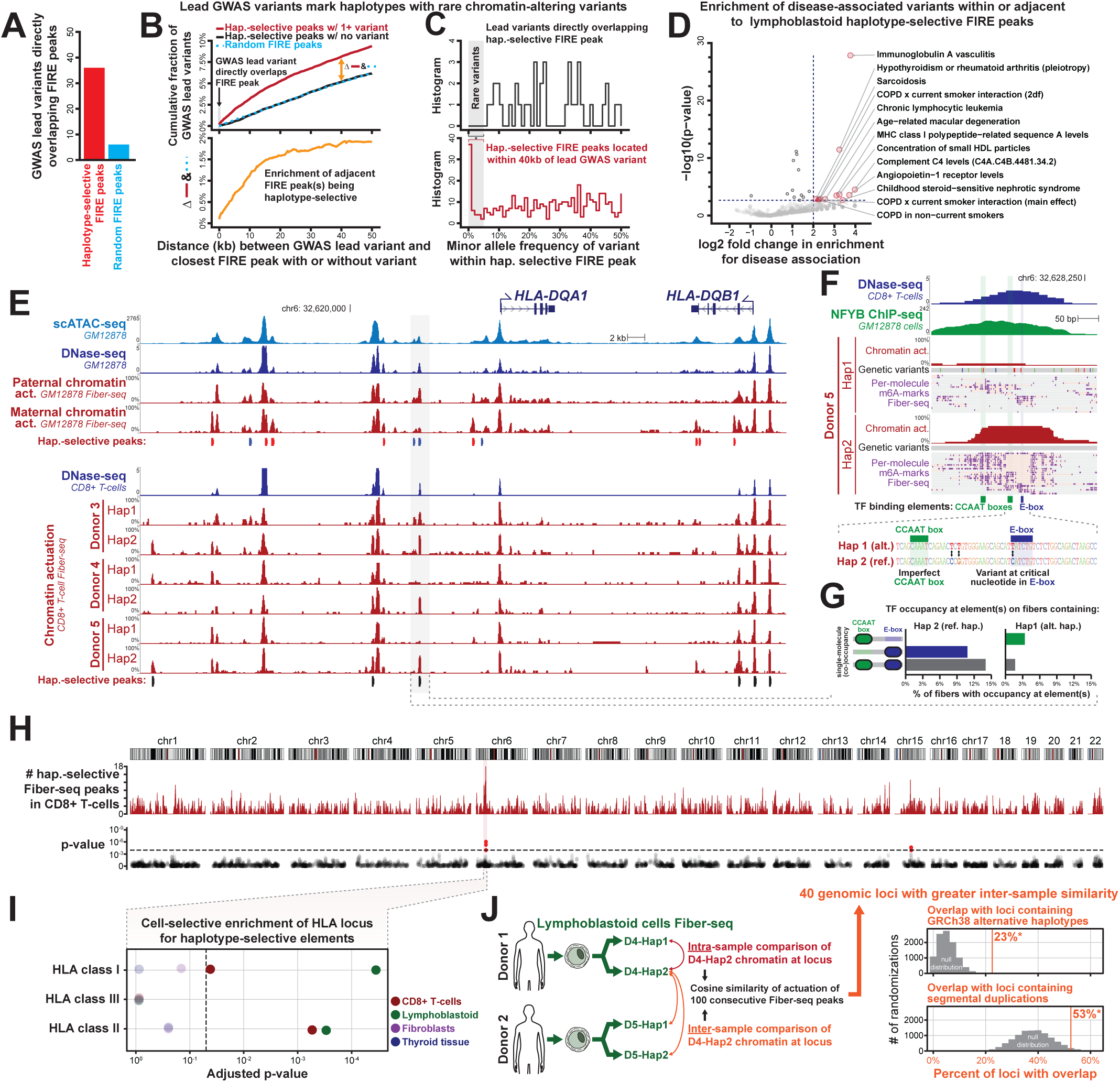
Haplotype-selective chromatin in the major histocompatibility complex. **A)** The number of haplotype-selective peaks (red) or random windows (blue) in GM12878 that overlap GWAS variants that are heterozygous in GM12878. **B)** Top, the fraction of lead GWAS variants that can be found within a specific distance (kbp) of: a haplotype selective peak with a genetic variant (red), a haplotype selective peak without genetic variants, and a random set of FIRE peaks of the same size. Bottom, the difference in the fraction of GWAS within a specific distance to a haplotype selective variant with a genetic variant versus a random set of FIRE peaks of the same size. **C)** Histogram of minor allele frequencies for GWAS lead variants that directly overlap haplotype-selective FIRE peaks. Red bars indicate variants within 40 kbp of a haplotype-selective FIRE peak containing a genetic variant. **D)** Enrichment of disease-associated variants within 40 kbp of haplotype-selective FIRE peaks for different disease associations. The x-axis shows the log2 fold enrichment, and the y-axis represents the p-value of a two-sided Fisher’s exact test. **E)** Haplotype-selective sites in the *HLA-DQA1*/*HLA-DQB1* locus for CD8+ T-cells sequenced to ∼30-fold coverage across three individuals. **F)** Haplotype-selective Fiber-seq patterns showing disruption of single-molecule Fiber-seq TF occupancy and chromatin actuation in haplotype 1 of Donor 5 (Hap 1. Alt) compared to the reference haplotype, which is haplotype 2 in Donor 5 (Hap 2. Ref). **G)** Histogram of the percent of fibers with TF occupancy at the CCAAT box and E-box across both haplotypes of Donor 5. Inaccessible fibers make up the remaining percentage of molecules not shown. **H)** Ideogram of the number of haplotype-selective sites in high-coverage CD8+ T-cells and the two-sided Fisher’s exact significance of the enrichment of haplotype-selective chromatin (*Methods*). **I)** Cell-selective enrichment of HLA class I, II, and III for haplotype-selective elements (two-sided Fisher’s exact test). **J)** Schematic of testing intra- versus inter-sample cosine similarity between four haplotypes from two donors (GM12878 and COLO829BL) and enrichment of inter-sample similarity within GRCh38 alternative haplotypes (p < 1e-04; permutation test n=10,000; *Methods*) and segmental duplications (p = 0.0357; permutation test n=10,000; *Methods*).

However, most lead GWAS variants are not thought to be disease-causal but rather are thought to be tagging a haplotype that contains neighboring variants that are mediating the disease or trait association, including neighboring rare variants with a large effect size (*36*). To evaluate whether these lead GWAS variants are tagging a putatively functional variant along the same haplotype, we quantified the incidence of haplotype-selective chromatin peaks within 500 kbp surrounding lead GWAS variants in GM12878. This demonstrated that lead GWAS variants were enriched for being located within 40 kbp of a peak with haplotype-selective chromatin (**Fig. 4B**). Notably, unlike lead GWAS variants that directly overlap elements with HSCA, these adjacent haplotype-selective peaks preferentially contained rare genetic variants with a minor allele frequency (MAF) of less than 5%, indicating that Fiber-seq is identifying rare variants along these haplotypes that are potentially mediating the disease/trait associations (**Fig. 4C**). Consistent with this, we found that the disease associations of these GWAS lead variants preferentially localized adjacent to GM12878 haplotype-selective chromatin elements were significantly enriched (p < 0.05, two-sided Fisher’s exact test) for diseases or traits related to the biological function of lymphoblast cells (**Fig. 4D, Supplemental Table 3**).

The per-molecule and near-nucleotide resolution of Fiber-seq permitted the dissection of the specific TF binding elements and variants that are likely mediating these disease associations. For example, haplotype-selective Fiber-seq patterns within the *MOG1/HLA-F* locus demonstrated that among the lead SNPs associated with platelet counts, which is frequently an immune-mediated phenotype, only two appear to be present within putative regulatory elements, with one (rs4713235) directly disrupting single-molecule Fiber-seq TF occupancy at a CTCF binding element (**Fig. S10**). Furthermore, the 75 kb region surrounding the *HLA-DQA1* gene contains 3,199 heterozygous genetic variants that distinguish the two haplotypes in GM12878 (**Fig. S11A**). However, TF footprinting using Fiber-seq in GM12878 cells and primary T-cells enabled the identification of specific variants mediating these haplotype-selective chromatin features (**Fig. 4E**). For example, TF footprint patterns in the Fiber-seq data exposed that among the 54 variants within a 1 kb region overlapping an *HLA-DQA1* promoter-proximal HSCA FIRE peak, rs9271894, which is associated with Celiac disease risk in biobank PheWAS studies (*37*) (OR 2.84, p-value 1e-250), appears to be likely driving this HSCA signal. Specifically, this *HLA-DQA1* promoter-proximal HSCA FIRE peak contained two TF footprints with altered occupancy between haplotypes, one encompassing a predicted high-affinity E-box and one encompassing a predicted low-affinity CCAAT box (**Fig. 4F,G**). However, in these samples with HSCA only the TF footprint encompassing the E-box also contained a heterozygous variant (rs9271894), which disrupts a central base within the E-box. Furthermore, single-molecule protein footprint co-occupancy analyses (*15*) of these two TF footprints demonstrated that TF occupancy at the CCAAT box is cooperatively dependent upon TF occupancy at the E-box (**Fig. 4G**), suggesting a central role of this E-box, and potentially of rs9271894 in mediating this HSCA signal. To investigate whether rs9271894 is driving this HSCA signal, we performed plasmid Fiber-seq (*38*) using a 400 bp region encompassing this FIRE peak, as plasmid Fiber-seq is a scalable approach for directly measuring how non-coding variants alter chromatin accessibility. Plasmid Fiber-seq demonstrated that altering rs9271894 alone was sufficient to alter chromatin accessibility of this peak (**Fig. S12, Supplemental Table 4**). Together, these findings prioritize rs9271894 as the likely functional variant mediating this non-coding *HLA-DQA1* Celiac disease association.

### The HLA locus contains the highest rate of chromatin epigenomic diversity

Next, we sought to evaluate whether certain genomic loci are preferentially marked by haplotype-selective chromatin, irrespective of their disease association. To accomplish this, we quantified the enrichment of haplotype-selective peaks within rolling windows of 100 peaks (*Methods*) using Fiber-seq data from lymphoblastoid cells, activated and sorted CD8^+^ T-cells, fibroblasts, and thyroid tissue from multiple unrelated donors. After removing imprinting regions, this demonstrated that the MHC region on chromosome 6 contained the most haplotype-selective chromatin in the entire human genome in CD8^+^ T-cells (corrected p-value 0.0013, Benjamini-Hochberg corrected FDR < 5%) (**Fig. 4I**). Notably, whereas this enrichment for haplotype-selective chromatin was selectively localized to the MHC region HLA class II and HLA class I loci in lymphoblastoid cells, CD8^+^ T-cells only showed this localization within the HLA Class II locus (**Fig. 4J**), reflecting differences in the biological roles of these loci in these distinct immune cell types. In contrast, although fibroblasts and thyroid tissue show numerous actuated elements within the MHC region (**Fig. S11B,C**), these samples did not show any enrichment for haplotype-selective chromatin within the HLA locus (**Figs. 4J, S8C**), indicating that although many MHC proteins are ubiquitously expressed across various cell types, the marked genetic diversity within the MHC region is only associated with haplotype-selective chromatin patterns in select cell types.

Given the marked divergence in chromatin accessibility between both haplotypes within the MHC region, we next sought to evaluate whether the haploid chromatin accessibility within this region showed more variability between both haplotypes within the same individual or between individuals. To accomplish this, we compared haploid Fiber-seq data from the COLO829BL and GM12878 lymphoblastoid cell lines, which were derived from two separate donors of different ages and sex (**Fig. 4K**). While the far majority of the haploid genome showed greater variability in chromatin accessibility between both donors, as would be expected, we did identify 40 extended genomic loci, including the MHC region, that showed greater chromatin accessible variability between both haplotypes within the same donor. Notably, these 40 genomic loci were significantly enriched for containing alternative haplotypes, like the HLA locus (permutation test n=10,000, p<0.0001) and segmental duplications (permutation test n=10,000, p<0.0357). This demonstrates that genomic regions containing some of the most genetically diverse human haplotypes (*39–41*) also show the most haplotype-selective chromatin patterns, consistent with the hypothesis that selective pressures and/or consequences of these diverse haplotypes are also at the level of altered gene regulatory patterns (*42*).

### Somatic autosomal epimutations mark the chromatin landscape of cell lines and tissue

We next sought to understand the mechanistic basis underlying haplotype-selective chromatin at the 44% of GM12878 elements with HSCA that did not contain a genetic variant directly within the peak (*i.e.,* ‘HSCA-noVar’ elements) (**Fig. 3C**). Notably, unlike GM12878 elements with HSCA that overlap a heterozygous genetic variant (*i.e.,* ‘HSCA-withVar’ elements), HSCA-noVar elements did not show any enrichment for being adjacent to lead GWAS variants (**Fig. 4B, S9C**), suggesting that haplotype-selective chromatin at HSCA-noVar elements may not be genetically determined. Furthermore, HSCA-noVar elements overwhelmingly are not previously undiscovered imprinting sites (**Fig. 3F**), raising the possibility that haplotype-selectivity at these sites may arise from somatic epimutations causing random autosomal mono-allelic chromatin accessibility. To test this hypothesis, we determined the stability of haplotype-selectivity at these elements between multiple tissues derived from the same individual, under the assumption that somatic epimutations would preferentially show divergent chromatin patterns between different tissues that arose from the same zygote. Specifically, we performed Fiber-seq on a lymphoblastoid cell line (COLO829BL) and melanoma cell line (COLO829T) derived from the same donor (**Fig. 5A**). To ensure consistent haplotype phasing between these two samples, we leveraged a near telomere-to-telomere assembly of COLO829BL using Fiber-seq, ultra-long Oxford Nanopore, and Hi-C (contig N50 140.01 Mbp with 18 chromosomes having telomere-to-telomere scaffolds), enabling us to unambiguously phase 84.2% of all Fiber-seq reads from COLO829BL and 83.9% of Fiber-seq reads from COLO829T. Overall, we found that COLO829BL had 710 peaks with genome-wide significant HSCA and that the haplotype-selectivity of these peaks was in strong agreement across two COLO829BL Fiber-seq replicates, including at HSCA-noVar elements (Pearson’s correlation > 0.9, p-value < 2.2e-16, two-sided *t*-test) (**Fig. 5B**). 29% (202/710) of these HSCA elements in COLO829BL also demonstrated some degree of chromatin actuation in the COLO829T cells, enabling us to evaluate the stability of HSCA at these 202 elements (**Fig. 5C**). We find that for imprinted sites (*R*=0.87) or COLO829BL HSCA-withVar elements (*R*=0.76), there is a strong correlation in the HSCA between both cell types (*e.g.,* elements selective to haplotype 1 in COLO829BL are also selective to haplotype 1 in COLO829T) (**Fig. 5D,E**), consistent with imprinted and HSCA-withVar elements largely having deterministic chromatin actuation dependent upon the underlying haplotype. In contrast, COLO829BL HSCA-noVar elements had markedly divergent haplotype-selective actuation between the two cell types (*R*=0.15) (**Fig. 5D,E**) indicating that this class of elements overall displays a non-deterministic pattern of chromatin actuation that appears to be occurring irrespective of their underlying haplotype, findings consistent with haplotype selective-chromatin arising from somatic epimutations.

**Figure 5.**
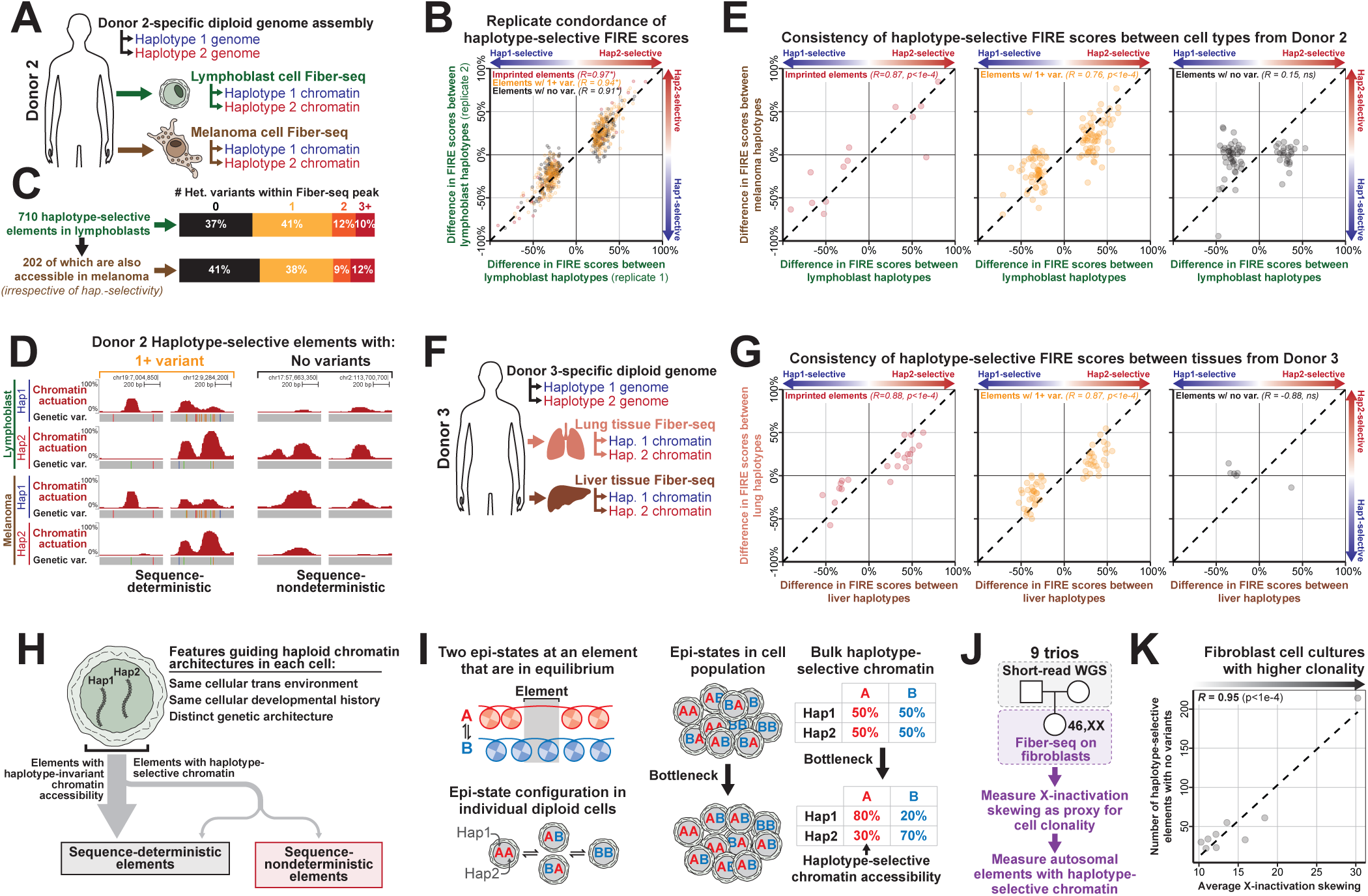
Deviation of haplotype-selective chromatin across cell types. **A)** Schematic of the sequencing of donor 2 of lymphoblast and melanoma cell lines. **B)** Replicate concordance of haplotype-selective percent actuation difference across two lymphoblast replicates for imprinted elements (red), elements with genetic variants between haplotypes (orange), and elements without genetic variants (black). Shown is the Pearson’s correlation; all correlations are significant with p < 2.2e−16 (two-sided t-test). **C)** The overlap of 710 lymphoblast haplotype-selective peaks with genetic variants and the subset of those peaks (202) that also overlap peaks within the melanoma cell line. **D)** Example of shared haplotype-selective elements and unique haplotype-selective elements between lymphoblast and melanoma cells. **E)** Concordance of haplotype-selective peaks between lymphoblast and melanoma cells within imprinted sites, sites with genetic variants, and sites without genetic variants (Pearson’s correlation; two-sided t-test: left p = 1e-4, middle p < 2.2e−16, right p = 0.21). **F)** Experimental design showing sequencing of liver and lung primary tissues from donor 3. **G)** Comparison of haplotype-selective peak concordance between liver and lung primary tissues, analyzed as in panel **E**. **H)** Conceptual model distinguishing haplotype-invariant elements from haplotype-selective elements with deterministic and non-deterministic patterns. **I)** Model illustrating how population bottlenecking in cell populations may generate additional haplotype-selective chromatin accessibility. **J)** Experimental approach for sequencing nine female fibroblast cell lines, using X-inactivation allelic skewing as a proxy for cell clonality. **K)** Correlation analysis between the number of haplotype-selective FIRE peaks without genetic variants and the average allelic skew across all X chromosome promoters, excluding the pseudoautosomal regions (Pearson’s correlation; two-sided t-test p-value = 7.9e−05).

To further evaluate whether autosomal HSCA-noVar elements have non-sequence deterministic patterns of chromatin accessibility, we used AlphaGenome (*43*) to predict haplotype-resolved FIRE signal at HSCA-noVar and HSCA-withVar elements within GM12878 cells (*43*). At HSCA-withVar elements, AlphaGenome-predicted DNase differences strongly correlated with measured FIRE actuation differences between both haplotypes (*R*=0.69), with 90.6% of peaks having concordant directionality between predicted and observed haplotype accessibility differences (**Fig. S13**). In contrast, at HSCA-noVar elements, AlphaGenome predicted minimal DNase differences between the two haplotypes (**Fig. S13**), consistent with the current sequence-based models being unable to predict epigenetic differences that arise independently of sequence, such as somatic epimutations.

To validate these findings in primary human tissues, we performed Fiber-seq on primary lung and liver tissue that were obtained from a separate single individual. Furthermore, we generated a diploid genome assembly from this individual to consistently phase the chromatin data between the two tissues. Overall, we found that the lung tissue had 88 peaks with genome-wide significant HSCA (**Fig. 5F**), with 98.9% (87/88) of these HSCA elements in lung also demonstrating some degree of chromatin actuation in the liver tissue. Consistent with the COLO829 data, imprinted elements (*R*=0.88) and HSCA-withVar elements (*R*=0.87) showed a strong correlation in the haplotype-selective chromatin actuation between both tissues. In contrast, lung HSCA-noVar elements showed markedly divergent haplotype-selective chromatin actuation between the two tissues (*R*=-0.88) (**Fig. 5G**), consistent with haplotype-selective chromatin at these HSCA-noVar elements arising from somatic epimutations (**Fig. 5H**).

These findings raise the prospect that some autosomal accessible chromatin elements somatically acquire non-deterministic chromatin actuation patterns that are mitotically stable (**Fig. 5I**), and as humans are diploid, this results in each haplotype having a divergent chromatin actuation pattern. One prediction of this model would be that bottlenecked samples would have higher rates of elements with HSCA, as a greater proportion of the cells within the Fiber-seq reaction would have arisen from the same founder cell with the somatic epimutation. To test this, we performed Fiber-seq on 46,XX fibroblast cell lines and measured both the number of HSCA-noVar elements in each sample, as well as the degree of skewing of X chromosome inactivation (XCI), which can be used as a proxy for how bottlenecked the sample is. Consistent with this prediction, we observed that fibroblast lines with greater evidence of clonal bottlenecking demonstrated more HSCA-noVar elements (*R*=0.95) (**Fig. 5J,K**). Together, these findings indicate that the autosomal accessible chromatin landscape of a cell is molded by somatic epimutations that create mitotically stable and non-genetically deterministic chromatin alterations.

### Somatically inactivated elements and domains along the X chromosome

The X chromosome in 46,XX cells is subjected to mitotically stable random XCI, creating an inactive X (Xi) and active X (Xa) chromosome within the same cell that is largely non-genetically deterministic (*44*, *45*). XCI represents an extreme case of mitotically stable somatic epimutations, and we sought to utilize Fiber-seq to explore the impact of XCI across every regulatory element along the X chromosome. To accomplish this, we applied Fiber-seq to two 46,XX cell lines [GM12878 (previously introduced) and MAN_1877] that have allelic XCI, enabling us to accurately map the chromatin architecture along the Xi and Xa based on parental haplotypes (**Fig. 6A**). Full-length long-read transcript sequencing of GM12878 cells showed that 99% of the long non-coding RNA *XIST* transcripts (*46–49*) originate from the maternal haplotype (**Fig. S14**), confirming that the Xi is invariantly the maternal haplotype in GM12878 cells. Using the Fiber-seq data, we categorized FIRE elements along the X chromosome as to whether they were preferentially somatically silenced on the Xi (*i.e.,* ‘Xa-selective’ accessibility) or Xa (*i.e.,* ‘Xi-selective’ accessibility), or showed similar chromatin accessibility along both the Xa and Xi (*i.e.,* ‘shared’).

**Figure 6.**
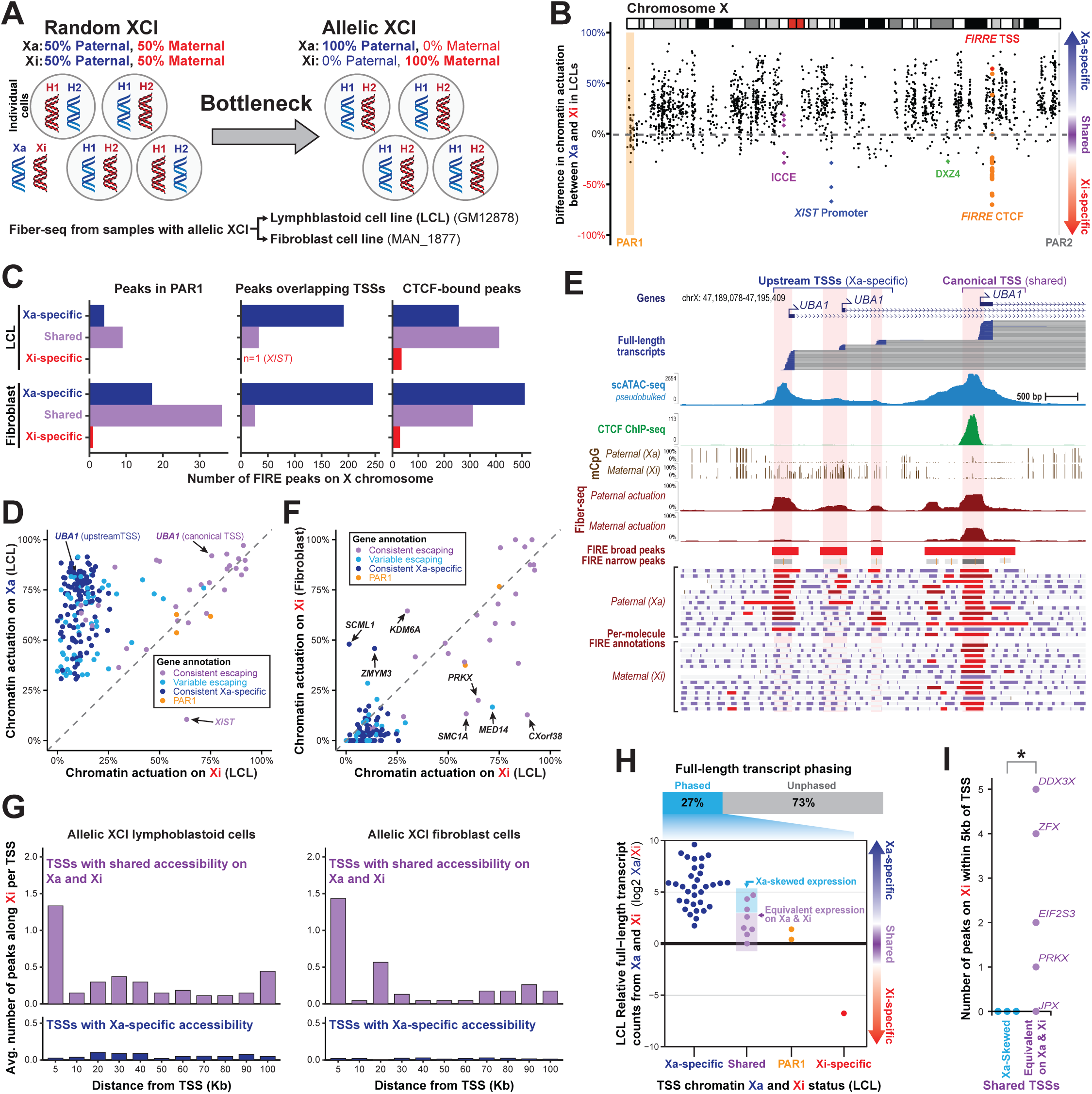
Haplotype-specific features of X chromosome inactivation (XCI). **A)** Schematic of culture-derived XCI skewing. **B)** Chromosome-wide comparison between percent actuation of the paternal (Xa) and maternal (Xi) haplotypes at each FIRE peak in LCL cells (GM12878). Pseudoautosomal regions (PAR1 & PAR2) are highlighted in orange and gray, respectively. **C)** Counts of FIRE peaks categorized as Xa-specific, Xi-specific, or Shared between both haplotypes for LCL (top) and fibroblast cells (bottom). FIRE elements are stratified by their location within or outside of PAR1 (left), and non-PAR1 elements are further subsetted to those that overlap a CTCF site (middle) or TSS (right). **D)** Scatterplot of LCL Fiber-seq percent actuation on the Xi (x-axis) and Xa (y-axis) for each TSS. Points are colored by XCI escape annotations from previous studies (*55*, *56*). **E)** *UBA1* promoter region comparing full-length transcript reads, scATAC-seq, CTCF ChIP-seq, mCpG, FIRE percent actuation, FIRE peaks, and representative Fiber-seq reads from the paternal (Xa) and maternal (Xi) haplotypes in LCLs (GM12878). **F)** Scatterplot of Fiber-seq percent actuation on the Xi in LCL (x-axis) and fibroblast (y-axis) cells. Points are colored as in panel **D**. **G)** The average number of escaping non-TSS FIRE peaks in LCL (left) and fibroblast (right) cells by absolute distance from TSSs. Counts are displayed separately for escaping TSSs (top, purple) and inactivated TSSs (bottom, blue). **H)** Full-length LCL transcript expression differences between the Xa and Xi for genes phased by Fiber-seq and displayed in **D**. Count differences are displayed as log2 fold-change between the haplotypes. Genes are stratified by the Fiber-seq classifications of their TSS FIRE peaks as in **c**. **I)** The number of escaping LCL non-TSS FIRE peaks within 5 Kb of each TSS in the shared category in **H**. Shared TSSs were grouped into high or low log2 fold-change in expression, highlighted with blue and purple in **H** (\**p*L=L0.04031; one-sided Wilcoxon rank sum test).

We observed that chromatin accessibility outside of the pseudoautosomal region 1 (PAR1) was predominantly impacted by XCI, with 63% of elements being Xa-specific, 34% being shared, and only 2.6% being Xi-specific elements (**Fig. 6B-D**). Aside from the *XIST* promoter, the majority (73.3%) of ‘Xi-selective’ elements were within *ICCE* (Inactive-X CTCF-binding Contact Element) (*50*), DXZ4 (*51*, *52*), and *FIRRE* (Functional Intergenic Repeating RNA Element) loci (*53*) – and 78.9% of these were CTCF elements (**Fig. S15**), which play a role in organizing three-dimensional mega-domains on the Xi (*54*). 66% of ‘shared’ elements were also CTCF elements, with the majority (59%) of X chromosome CTCF sites having similar accessibility along both the Xa and Xi. Notably, boundaries separating chromatin domains subjected to XCI and escaping XCI often contained CTCF elements with protein occupancy on the Xi (**Fig. S16**). For example, *UBA1* has four distinct accessible TSSs in GM12878 cells, with the three upstream TSSs being Xa-specific, and the canonical downstream TSS escaping XCI (**Fig. 6E**). The canonical XCI escaping *UBA1* TSS is bookended by an upstream CTCF element that shows single-molecule TF occupancy on both the Xa and Xi (90% and 59% occupancy, respectively) (**Fig. 6E**). Overall, these findings demonstrate that CTCF occupancy along the human X chromosome preferentially escapes XCI mediated somatic silencing and that CTCF occupancy along chromosome X may serve as boundary elements insulating chromatin domains from XCI, features that have been previously proposed in mice (*52*).

### Escaping promoter-proximal elements enable productive transcription from the Xi

16% of TSSs for transcribed genes in GM12878 cells were also marked by ‘shared’ elements (**Fig. 6C**). Notably, these TSSs largely corresponded to genes known to escape XCI based on prior transcript sequencing data (*55*, *56*) (**Fig. 6D**), as well as paired GM12878 long-read transcript data. However, the GM12878 transcript sequencing data could only phase 27% of these genes owing to the paucity of heterozygous coding genetic variation along the human chromosome X in GM12878 cells. We sought to directly test the relationship between promoter accessibility and transcript production on the Xi using the paired haplotype-phased long-read transcript and Fiber-seq data. As expected, we observed highly skewed expression in genes with Xa-specific or Xi-specific TSS accessibility (**Fig. 6H**). However, genes with similar TSS accessibility between the Xa and Xi showed heterogeneous expression, with some genes having similar transcript levels between the Xa and Xi (n=5), and others showing Xa-biased expression to a similar degree to that of genes with Xa-specific TSSs (n=3).

Given the widespread accessibility that we observed along the Xi, we hypothesized that the escape of promoter-proximal regulatory elements may augment productive transcription from the Xi. To address this, we computed the average number of escaping non-TSS elements within 5-10 Kbp bins away from escaping or inactivated TSSs (**Fig. 6G**). This revealed enrichment of escaping regulatory elements within 100 Kbp of escaping TSSs, with the largest difference (51-to 71-fold increase) within 5 Kbp of the escaping TSS. Notably, genes that have similar TSS accessibility and transcript levels between the Xa and Xi had significantly more promoter-proximal escape elements compared with genes that have similar TSS promoter accessibility but higher transcription from the Xa (12 vs 0, p-value = 0.04, one-sided Wilcoxon rank sum test; **Fig. 6I**). Overall, these results demonstrate that promoter accessibility is necessary but often not sufficient for robust gene expression from Xi, and that the simultaneous escape of promoter-proximal regulatory elements plays a pivotal role in maximizing the transcriptional potential of escape genes.

## Discussion

We demonstrate how Fiber-seq can enable a comprehensive view of chromatin accessibility across the 6 Gbp diploid human genome with single-molecule and single-haplotype precision at near-nucleotide resolution. Specifically, we present a novel semi-supervised machine learning tool (FIRE) and show that FIRE provides a more comprehensive and quantitative snapshot of a cell’s accessible chromatin landscape, overcoming previously known and unknown biases inherent to our existing catalog of regulatory elements. Importantly, FIRE enables the accurate *de novo* construction of a cell’s haplotype-resolved chromatin-accessible landscape directly using long-read sequencing data, enabling the advent of synchronous multi-omic long-read sequencing that can fine-map the chromatin mechanisms by which disease-associated haplotypes cause human disease (*15*, *16*), as well as resolve chromatin architectures within complex genomic loci like segmental duplications (*57*), centromeres (*58*, *59*), and transposons (*24*).

Overall, we find that genomic loci marked by the most genetic diversity within the human population (*39–41*, *60*) also contain the most haplotype-selective chromatin in the human genome. This finding indicates that genetic diversity at these loci is likely directly impacting gene regulatory patterns. As changes in gene regulation are known to play a dominant role in human speciation (*42*), it is likely that these haplotype-resolved maps of gene regulation will improve our understanding of the specific genetic variants that make us humans. For example, we show that Fiber-seq can localize the specific regulatory elements and TF binding events that underlie human disease associations, including those within the HLA locus. The HLA locus has been definitively established as the primary genetic locus underlying human-specific immune responses and autoimmune disease risk. However, the HLA locus is one of the most genetically complex loci in the human genome, with genetic variants occurring once every 20 bases along many disease-associated haplotypes – a rate of genetic diversity that stifles short-read-based approaches. Although it is largely assumed that only protein-coding variants mediate HLA locus autoimmune disease risk, emerging data have indicated that non-coding variants play a critical role in autoimmunity through altering HLA gene expression (*61–63*), which in turn can dramatically impact how immune cells respond to antigens. We show that Fiber-seq can pinpoint specific genetic variants within the HLA locus that drive disease associations, exposing that the Celiac disease-associated SNP rs9271894 disrupts cooperative TF occupancy at an E-box and adjacent CCAAT box immediately upstream of *HLA-DQA1* (**Fig. 4H**), which is one of the primary immune genes associated with Celiac disease (*64*). Although the exact impact of this variant on immune cell function is unknown, Fiber-seq nonetheless enabled us to localize which of the thousands of variants within the HLA region actually show a molecular phenotype at the chromatin level. Given these findings, we anticipate that the broader application of Fiber-seq will dramatically alter how we study and understand the contribution of non-coding variation within the HLA locus to human disease risk and speciation.

It is well known that the establishment of X inactivation differs dramatically between mice and humans, both in terms of the order of events and the epigenetic silencing mechanisms (*45*). Although short-read epigenetic maps of the Xi and Xa can be readily established using hybrid mice, this is not readily accomplishable in humans owing to the lack of genetic diversity along chromosome X. We demonstrate that Fiber-seq enables the precise delineation of chromatin elements along the Xa and Xi in humans, revealing widespread chromatin accessibility across the Xi and exposing that promoter accessibility along the Xi is necessary but often not sufficient for robust gene expression from Xi. Epigenetic editing approaches are actively being investigated for the treatment of X-linked diseases (*65*), and we anticipate that Fiber-seq will provide a roadmap for these translational efforts.

Outside of the X chromosome, we find that ∼99% of autosomal accessible elements show nearly identical levels of chromatin accessibility between the two haplotypes in these bulk measurements. This demonstrates that, on average, chromatin accessibility is largely deterministic (*i.e.,* a reflection of a cell’s genomic sequence, cellular trans environment, and developmental history), indicating the potential of computational tools for accurately predicting the vast majority of a cell’s steady-state chromatin accessibility landscape (*43*). As transvection is not known to impact human gene regulation, the chromatin patterns along each haplotype are independently formed during a cell’s and organism’s life span, a system ripe for stochastic deviations in the accessibility pattern between both haplotypes as a result of epigenetic noise and subsequent memory (*66*). As described above, Fiber-seq exposed numerous elements with HSCA that appear to be primarily driven by underlying heterozygous genetic variants. However, ∼40% of elements with HSCA lacked an underlying genetic variant. We show that these elements largely appear to have non-deterministic behavior (*i.e.,* their chromatin accessibility deviates between both haplotypes in a manner that cannot be explained by the genetic architecture of their underlying haplotype). Importantly, we prove that these elements are not imprinted sites and show that elements with non-deterministic behavior can be observed in primary human tissues. Together, these findings demonstrate that autosomal accessible chromatin elements can somatically acquire non-deterministic chromatin actuation patterns that are mitotically stable and divergent between haplotypes, a pattern consistent with these being sites of somatic epimutations (*67*). Somatic epimutations are becoming increasingly recognized using CpG methylation and transcript sequencing (*68*, *69*), but have been challenging to systematically study at the level of chromatin (*70*) owing to the inability to accurately haplotype phase short sequencing reads at elements that lack underlying heterozygous variants within a sample – a challenge that is overcome using Fiber-seq. Notably, we find that clonal expansions increase the rate of somatic epimutations, consistent with these somatic epimutations arising from founder events. Together, these findings indicate that the autosomal accessible chromatin landscape of a cell is molded by somatic epimutations that create mitotically stable and non-genetically deterministic chromatin alterations.

Autosomal monoallelic expression is a well-documented feature of normal human tissues (*71*) and can influence disease risk (*72*). While this phenomenon is thought to arise from somatic epimutations, with roles established in both rare (*73*) and common (*74*) diseases, current catalogs are largely based on CpG methylation, owing to the haplotype-resolving power of single-molecule methylation assays. Here, we show that autosomal, mitotically stable, haplotype-selective regulatory patterns can arise independently of CpG methylation (**Fig. 3E-H**), and that most regulatory elements consistent with somatic epimutations lack pronounced haplotype-specific CpG methylation differences (**Fig. 3C**), though we cannot exclude that differential methylation at distal regulatory elements may contribute to haplotype-selective accessibility at some loci. These findings suggest that epigenetic memory at these loci is primarily encoded in local chromatin architecture rather than CpG methylation or the trans environment. More broadly, our results reveal that somatic epimutations are pervasive and impose an epigenetic ceiling on the deterministic influence of genetic variation in shaping the human diploid epigenome.

## Limitations of the Study

Several limitations should be considered when using FIRE or, more broadly, when interpreting Fiber-seq data. First, it is important to ensure appropriate data quality and coverage, especially when comparing multiple datasets. Specifically, the power to detect FIRE peaks (FDR < 0.05) is directly related to coverage. While a strength of Fiber-seq is that molecules are sequenced regardless of accessibility, this also necessitates genome-wide sequencing, which can result in variance in coverage between samples or experiments. Consequently, comparing accessibility signal at a union set of peaks between samples is often more informative than simply comparing the number of intersecting peaks. Of note, deeply sequenced samples often are powered to identify FIRE peaks with less than 15% chromatin actuation, but the biological significance of these lowly actuated peaks remains to be seen. Second, FIRE infers the presence of regulatory elements from m6A modification patterns but cannot directly identify which TFs or protein complexes are bound. Confidently resolving specific-factor occupancy requires complementary approaches, such as ChIP-seq/CUT&RUN/DiMeLo-seq or motif-guided footprint analysis. Third, our characterization of somatic epimutations is inherently conservative: we define HSCA-noVar elements as those lacking any overlapping heterozygous variant and require genome-wide significance, meaning the true prevalence of non-genetically determined haplotype-selective chromatin may be higher than reported. Moreover, while we demonstrate that these elements are mitotically stable, non-deterministic, and amplified by clonal bottlenecking, we do not propose a specific molecular mechanism for their establishment or maintenance. Although we observe somatic epimutations in primary liver and lung tissue, our most detailed characterization relies on GM12878, a clonal cell line in which generation and long-term culture may have reshaped clonal dynamics. Finally, the FIRE model was trained on human GM12878 Fiber-seq data using paired DNase-seq and CTCF ChIP-seq peaks as positive labels. Although we demonstrate generalization to other TFs, cell types, and organisms (*24*), regulatory elements that systematically diverge from human DNase-seq or CTCF ChIP-seq patterns may be underrepresented using FIRE.

## Supplemental figure legends

**Figure S1. Motivation for the FIRE model: heterogeneity in single-molecule data.**

**A)** Global mean m6A rate (total m6A calls divided by total A+T base pairs) for each of the 21 GM12878 Fiber-seq libraries used in training of the FIRE model, prepared with Hia5 concentrations ranging from 50U to 500U.

**B)** Heterogeneity of m6A concentrations between reads in the Fiber-seq experiments used in the training of the FIRE model.

**C)**. Distribution of the length of methyltransferase-sensitive patches (MSPs) across the Fiber-seq experiments used in training the FIRE model.

**D)** Performance of different classification thresholds for identifying actuated regulatory elements on chromosome 20. True positives (TP; y-axis) are defined as MSPs overlapping DNase I hypersensitive sites (DHSs), and false positives (FP; x-axis) are MSPs outside of DHSs. Each point represents a different threshold value applied to either the FIRE estimated precision score (red) or MSP length alone (gray). The threshold for the final point was used for downstream analyses (94.9% estimated precision for FIRE; see **Supplemental Note**).

**E)** Distribution of MSP length within DNase peaks and outside of DNase peaks (grey), stratified by inferred FIRE elements (red) and nucleosomal linker regions (purple).

**Figure S2. Training of the FIRE model.**

**A)** Schematic of training Fiber-seq inferred regulatory elements (FIREs) using XGBoost within the Mokapot framework.

**B)** Schematic of windows along single reads used for calculating features in the FIRE model. Descriptions of each feature are listed in **Supplemental Table 1**.

**C)** Ranking the importance of features in the XGBoost model using the feature score (F score), which sums up how many times each feature is split on within the model. Note that feature bins were labeled 0-8, making the 4^th^ bin the central bin of the MSP.

**Figure S3. FIRE elements and their underlying m6A calls.** UCSC genome browser screenshots of a low accessibility regulatory element **(A)**, a high accessibility element **(B)**, and two sites without regulatory elements **(C,D)**. Shown in each panel in order are: the gene models, ATAC signal, percent of Fiber-seq reads with FIRE elements, wide and narrow FIRE peak calls, raw m6A calls for individual Fiber-seq reads, and FIRE calls for individual Fiber-seq reads in the same order.

**Figure S4. The aggregate FIRE score and peak calling with the FIRE method.**

**A)** Schematic of calculating a false discovery rate (FDR) for the aggregated FIRE score. top) the aggregated FIRE score across multiple Fiber-seq molecules. middle) aggregate FIRE score for Fiber-seq reads with a shuffled start position within the chromosomes. bottom) FDR track calculated from the null distribution of FIRE scores and peak calls using a 5% FDR threshold.

**B)** Number of base pairs genome-wide with an aggregate FIRE score above a threshold (x-axis) for both the observed FIRE elements (red) and the shuffled Fiber-seq reads (gray). Dashed lines are drawn at 1% and 5% FDR thresholds.

**C)** The FDR score (y-axis) vs FIRE score threshold (x-axis) for 130-fold Fiber-seq coverage of G12878 and 30-fold coverage.

**D)** The correlation of FIRE scores within FIRE peaks between 130-fold Fiber-seq coverage of G12878 (x-axis) and 30-fold coverage (y-axis).

**E)** Correlation of FIRE percent actuation and DNase-peak signal within GM12878 for FIRE peaks faceted by FIRE peak size (left to right) and whether GC correction was (bottom) or was not (top) applied (Pearson’s correlation; p < 2.2e−16 two-sided t-test).

**Figure S5. Nucleosome calling is robust to parameter choice and does not substantially affect FIRE scores.**

**A)** Nucleosome length distributions from GM12878 Fiber-seq data across a 3x3 grid of ‘ft add-nucleosomes’ parameter combinations. Columns vary coupled minimum nucleosome length (nl) and minimum combined nucleosome length (cnl); rows vary minimum distance added (md). The selected default parameters (nl=75, cnl=100, md=25; red) produce a modal nucleosome length near the canonical 147 bp (gray dashed line) with the highest nucleosome yield (red dashed horizontal line) and smooth ∼10 bp periodic structure consistent with nucleosome breathing. All other parameter combinations are shown in black.

**B)** Nucleosome length distributions using default parameters validated across additional samples, including cell lines (A549, K562, THP-1), a reference genome (CHM13, HG002), primary tissues (liver, lung), and tumor/normal pairs (COLO829BL). All samples show a consistent mode near 147 bp with preserved breathing periodicity (y-axes scaled independently).

**C)** FIRE score robustness to nucleosome calling parameters. Each panel shows a hex-bin density plot comparing per-peak FIRE scores computed with the default nucleosome calling parameters (x-axis) versus an alternative parameter combination (y-axis) in GM12878. Pearson correlation coefficients (r) are annotated; the dashed diagonal marks perfect agreement. All nine parameter combinations yield r > 0.99, demonstrating that FIRE element classification is largely insensitive to nucleosome calling parameters. P-values are from a two-sided Pearson correlation *t-test*.

**Figure S6. Quantitative agreement between FIRE and DNase-seq chromatin accessibility across ENCODE cell lines.**

**A)** Hexbin density plots of DNase signal (log10) versus FIRE percent actuation for all union peaks, faceted by cell line (rows: A549, Jurkat, THP-1) and peak category (columns: FIRE-only, shared, DNase-only). Pearson correlation and the linear regression fit (red line) are shown in each panel.

**B)** Fiber-seq coverage distribution at DNase-only peaks, stratified by cell line. Peaks are colored by whether they exceed 15% FIRE actuation (red) or not (blue). Coverage is clamped at 150+.

**C)** Linear regression of DNase signal on FIRE percent actuation for shared peaks, stratified by peak size (0-200 bp, 200-250 bp, >250 bp). Regression equations and Pearson R values are shown per size group. Labeled points indicate predicted DNase signal at 50% actuation. Larger peaks show steeper slopes and stronger correlations, consistent with DNase sensitivity scaling with element width.

**D)** DNase signal distributions (violin plots with median crossbars) binned by FIRE percent actuation for shared and FIRE-only peaks, stratified by peak size (0-200 bp vs >200 bp). DNase signal increases monotonically with FIRE actuation, and larger elements show consistently higher DNase signal at matched actuation levels.

**E)** Median fold difference in DNase signal between large (>200 bp) and small (0-200 bp) peaks at each FIRE actuation bin. The dashed red line indicates parity (fold difference = 1). Median fold differences across all bins are annotated per cell line.

**F)** Correlation between CTCF ChIP-seq enrichment (ENCFF843VHC) and FIRE percent actuation at K562 CTCF peaks. Each hexbin is colored by the log10 number of peaks. Pearson correlation coefficient and p-value are shown in the upper left, computed using a two-sided t-test (n=44,722).

**Figure S7. Validation and unique features of FIRE elements.**

**A)** Correlation of FIRE percent actuation and DNase-peak signal within K562 for FIRE peaks faceted by FIRE peak size (left to right; Pearson’s correlation; p < 2.2e−16 two-sided t-test).

**B)** Pairwise correlation plots among FIRE, DNase-seq, and ATAC-seq signal at shared K562 peaks. Left: DNase signal versus FIRE % actuation. Middle: ATAC signal versus FIRE % actuation. Right: DNase signal versus ATAC signal. Short-read signal axes are log10-scaled. Pearson R values are shown per panel.

**C)** Association between the % of cells with single-cell ATAC-seq signal, versus the size of the regulatory element for only those elements that show a FIRE actuation of >70%. As expected, there is a strong association between element width and scATAC-seq signal, but this does not approach 100%, demonstrating that even at very large elements, there is a limit of detection in scATAC-seq signal.

**D)** Distribution of the GC content for unique FIRE peaks (top) and FIRE peaks shared with ATAC or DNase (bottom) stratified by peak size. Labels indicate the median GC content and the number of elements plotted in the distribution.

**E)** Enrichment of transcription (TF) factor motifs within FIRE only peaks versus all FIRE peaks. Enrichments and p-values calculated using the HOMOR software.

**F)** The proportion of methylated adenines over total adenines (either strand). All fibers with a FIRE element overlapping FIMO-predicted REST motif (green) are split by those with an overlapping footprint called by ft footprint (orange) and those that are not footprinted (blue). The x-axis represents the position relative to the start of the REST motif.

**Figure S8. Relationship between sequence composition and chromatin accessibility at FIRE peaks.**

**A)** Percentage of bases in A or T homopolymers (>= 2) versus Accessibility (%), with marginal histograms.

**B)** Count of bases in A or T homopolymers versus Accessibility (%), with marginal histograms (count capped at 100).

**C)** GC content versus FIRE accessibility (%), max DNase signal, and max ATAC signal, with linear regression fits (black lines). Pearson correlation coefficients (R) are shown in each panel. Hex bin color indicates point density.

**Figure S9. Genetic and chromatin diversity in GM12878 Fiber-seq across the major histocompatibility complex (MHC).**

**A)** Correlation analysis between CTCF ChIP-seq signal and three chromatin accessibility measures: ATAC-seq (left), DNase-seq (middle), and FIRE (right). Values represent Pearson’s correlation coefficients with p-values determined by two-sided t-tests.

**B)** Histogram of centered absolute difference in percent accessibility between replicates (blue) and haplotypes (red) for COLO829BL (Wilcoxon rank-sum test; W = 6.461e+09; p-value < 2.2e-16).

**C)** GWAS variant enrichment near haplotype-selective chromatin accessibility (HSCA) peaks. (Top) Cumulative fraction of heterozygous GWAS risk variants in GM12878 as a function of distance to the nearest FIRE peak. GWAS variants were sourced from the PICS2/GWAS Catalog and filtered to heterozygous sites in GM12878. Peak categories were randomly balanced to equal sample sizes before computing distances. Solid lines denote observed peak sets: all HSCA peaks (dark red) and the subset harboring heterozygous variants (orange). Dashed lines denote three control sets: randomly sampled FIRE peaks (light blue), FIRE peaks matched to the heterozygous SNV count distribution of HSCA peaks (green), and genomic-position-shuffled HSCA peaks (gray). Results are shown for all GWAS variants (left) and excluding the MHC region (chr6: 28.5–33.5 Mbp; right). (Bottom) Number of heterozygous GWAS risk variants directly overlapping peaks, normalized per 1,000 peaks to account for differences in set size. P-values are two-sided Fisher’s exact tests comparing each control category to HSCA.

**Figure S10. Haplotype-selective Fiber-seq patterns within the MOG1/HLA-F locus.**

**A)** Haplotype-selective Fiber-seq patterns within the *MOG1/HLA-F* locus for 4 CD8+ cell lines. Annotated with red lines are lead GWAS SNPs associated with platelet counts.

**B)** Fiber-seq results for an individual (donor 3) homozygous for the rs29269 SNP show that the SNP falls between regulatory elements and does not disrupt them.

**C)** Fiber-seq results for an individual heterozygous (donor 1) for the rs4713235 SNP show that the SNP directly disrupts single-molecule Fiber-seq transcription factor occupancy at a CTCF binding element.

**D)** FIRE haplotype-selective peaks for donors 1 and 3 (two-sided Fisher’s exact test).

**E)** Chromatin accessibility for each haplotype across donors 1 (het), 3 (homo ref.), and 4 (homo alt.) at position rs4713235 (two-sided Fisher’s exact test).

**Figure S11. Genetic and chromatin diversity in GM12878 Fiber-seq across the major histocompatibility complex (MHC).**

**A)** IGV screenshot of the genetic diversity at the *HLA-DRB1* locus for phased GM12878 Fiber-seq reads.

**B)** Density of FIRE peaks for different samples across chromosome 6, where the color represents the cell type of the sample. Fibroblast samples are orange, Thyroid samples are blue, CD8+ samples are red, Lymphoblastoid cell lines are green, and the grey box highlights the MHC region.

**C)** Ideogram depicting enrichment of haplotype-selective sites across chromosomes in sliding windows of 100 FIRE peaks for multiple cell types. Statistical significance was determined using a two-sided Fisher’s exact test with Benjamini-Hochberg correction for multiple testing (*Methods*).

**Figure S12. Nucleosome occupancy analysis of plasmid Fiber-seq constructs.**

**A)** Alignment of the four plasmid Fiber-seq constructs. Top track shows chromatin actuation derived from CD8+ T-cell Fiber-seq data. The four promoter sequences are shown aligned to the Hap 2 reference sequence (blue), with black tick marks indicating variant positions.

**B)** Differential nucleosome occupancy between H2_ref and each comparison construct (H1_wild, 1_edit (rs9271894), 3_edit) across a 3 kb region centered on the promoter. Each subplot shows the nucleosome occupancy of H2_ref minus the indicated sample after NPR-matched subsampling. The blue shaded region denotes the promoter. Positive values (blue fill) indicate higher nucleosome occupancy in H2_ref; negative values (colored fill) indicate higher occupancy in the comparison sample. Dashed lines mark the positions of variants. All three comparisons show a consistent increase in nucleosome occupancy for promoter sequences containing the C>T E-box variant.

**C)** Zoomed view of the promoter region from (**B**).

**D)** Pairwise differential nucleosome occupancy among H1_wild, 1_edit (rs9271894), and 3_edit within the promoter region, demonstrating near-equivalent nucleosome occupancy across these constructs.

**Figure S13. Correlation between FIRE haplotype accessibility differences and AlphaGenome-predicted DNase sensitivity differences.**

**A)** Hexbin density plots of FIRE percent actuation difference (hap1 - hap2) versus AlphaGenome-predicted DNase difference (hap1 - hap2) for regulatory elements without sequence variants (left) and with one or more SNVs or SVs (right). Pearson R and Spearman rho are shown. Dashed red line indicates a linear regression fit; dashed gray lines mark zero on each axis. X-axis values are clamped to [-0.5, 0.5] for display; correlation values and regression lines are computed on unclamped values. Imprinted loci are excluded.

**B)** Stacked bar plot showing the fraction of peaks where the AlphaGenome-predicted DNase difference has the same sign as the FIRE actuation difference (concordant), the opposite sign (discordant), or is exactly zero, stratified by variant status. Peaks with zero FIRE difference are excluded.

**C)** Same comparison as (**A**), stratified by imprinting status (columns) and variant status (rows). Per-group Pearson R and Spearman rho values are shown.

**D)** Same comparison as (**A**), stratified by total variant count (SNVs + SVs): 0 variants, 1 variant, 2-4 variants, and 5+ variants. Correlation increases with variant count, indicating that sequence differences between haplotypes increasingly explain both FIRE accessibility and predicted DNase sensitivity differences.

**Figure S14. Haplotype-specific chromatin and gene expression of XIST.**

**A)** *XIST* promoter region comparing GM12878 scATAC-seq, CTCF ChIP-seq, mCpG, FIRE percent accessibility, FIRE peaks, and representative Fiber-seq reads from the paternal (Xa) and maternal (Xi) haplotypes.

**B)** Barplot showing *XIST* transcript counts for the GM12878 paternal (Xa) and maternal (Xi) haplotypes.

**Figure S15. Haplotype-specific chromatin at Inactive X structural elements.** GM12878 browser tracks for the repeat-rich regions, which are known to organize the Xi into distinct three-dimensional mega-domains via Xi-specific CTCF binding sites: **A)** DXZ4, **B)** FIRRE (Functional Intergenic Repeating RNA Element), and **C)** ICCE (Inactive-X CTCF-binding Contact Element) loci. Shown are tracks for scATAC-seq, CTCF ChIP-seq, mCpG, FIRE percent accessibility, and FIRE peaks. Short-read CTCF data is not available for DXZ4 due to mapping limitations within segmental duplications.

**Figure S16. Prevalence of XCI escape by proximity to escaping CTCF elements.**

Barplots showing the increased prevalence of escaping FIRE peaks in proximity to escaping CTCF sites compared with inactivated CTCF sites. Peaks are grouped by genomic distance from CTCF sites in 25 kb bins and represent the difference in the proportions of peaks escaping XCI when proximal to either escaping or inactivated CTCF sites. Escape prevalence was calculated using all non-CTCF peaks (left) and only TSS containing non-CTCF peaks (right).

## Methods

### Cell Culture

GM12878 were purchased from the Coriell Institute for Medical Research and cultured in RPMI 1640 Medium supplemented with 2mM L-glutamine,10% fetal bovine serum, and 100 I.U./mL penicillin/100 μg/mL streptomycin. Cells were maintained in a 37 °C humidified incubator under 5% carbon dioxide. Cell cultures were split every 3-5 days.

### Fiber-seq Library Preparation and Sequencing

Cells were permeabilized and treated with Hia5 enzyme as previously described (*75*). Specifically, 1 million cells were washed with PBS and then resuspended in 60 μl Buffer A (15 mM Tris, pH 8.0; 15 mM NaCl; 60 mM KCl; 1mM EDTA, pH 8.0; 0.5 mM EGTA, pH 8.0; 0.5 mM Spermidine) and 60 μl of cold 2X Lysis buffer (0.1% IGEPAL CA-630 in Buffer A for GM12878 and GM24385; 0.2% IGEPAL CA-630 in Buffer A for UDN318336 fibroblasts) was added and mixed by gentle flicking then kept on ice for 10 minutes. Samples were then pelleted, the supernatant removed, and then resuspended in 57.5 μl Buffer A and moved to a 25°C thermocycler. 0.5 μl of Hia5 MTase (100 U) and 1.5 μl 32 mM S-adenosylmethionine (NEB B9003S) (0.8 mM final concentration) were added, then carefully mixed by pipetting the volume up and down 10 times with wide bore tips. The reactions were incubated for 10 minutes at 25°C, then stopped with 3 μl of 20% SDS (1% final concentration) and transferred to new 1.5 mL microfuge tubes. High molecular weight DNA was then extracted using the Promega Wizard HMW DNA Extraction Kit A2920. PacBio SMRTbell libraries were then constructed using the Fiber-seq treated gDNA following the manufacturer’s SMRTbell prep kit 3.0 procedure (https://www.pacb.com/wpcontent/uploads/Procedure-checklist-Preparing-whole-genome-and-metagenome-libraries-usingSMRTbell-prep-kit-3.0.pdf).

### Nucleosome and MSP (methyltransferase sensitive patch) and calling

Nucleosome calling is performed by identifying stretches of DNA that are protected from Hia5 (i.e. do not have m6A signal). The rate of false positive m6A calls in nucleosomes is very low when using fibertools (*26*) allowing for a heuristic to perform as well as or better than our previous HMM caller (*75*). There are three parameters in our heuristic nucleosome calling that can be adjusted: the minimum nucleosome length (n, default 75), the minimum combined nucleosome length (c, default 100), and the minimum extension to nucleosome length (e, default 25). These three parameters impact the four phases in nucleosome calling.

1. Call all regions that have no m6A events for at least n bases a candidate nucleosome.
2. Call all regions of size c or more that have only one internal m6A (putative false positive) a candidate nucleosome.
3. Extend the length of nucleosomes identified in phases one and two if by spanning one additional m6A e bases of unmodified sequence would be added to the nucleosome length.
4. Recursively apply step three till no nucleosomes change.

After nucleosomes are called MSPs are operationally defined as all the regions between nucleosome footprints in the Fiber-seq data. This process is automated through the fibertools subcommand “add-nucleosomes”.

### Training of Fire-seq inferred regulatory elements

For training data, we generated 21 different GM12878 Fiber-seq experiments with a range of under- and over-methylated experimental conditions to ensure we captured a broad range of percent m6A (**Fig. S1**; 5.8-13.3%) to ensure our model could generalize to new samples with varying levels of m6A. We merged these sequencing results and randomly selected 10% of the Fiber-seq reads from 100,000 randomly selected DNase I and CTCF ChIP-seq peaks for mixed-positive labels and 100,000 equally sized regions that did not overlap DNase or CTCF Chip-seq for negative labels. We then generated features for each of these MSPs, including length, log fold enrichment of m6A, the A/T content, and windowed measures of m6A around the MSP (**Supplemental Table 1**) and held out 20% of the Fiber-seq reads to be used as test data.

To carry out semi-supervised training, we used an established method, Mokapot (*25*), which we summarize below. In the first round of semi-supervised training, Mokapot identifies the feature that best discriminates between our mixed-positive and negative labels and then selects a threshold for that feature such that the mixed-positive labels can be discriminated from the negative labels with 95% estimated precision (defined below). The subset of mixed-positive labels above this threshold is then used as an initial set of positive labels in training an XGBoost model with five-fold cross-validation (*28*). Then this process is iteratively repeated, using the learned prediction from the previous iteration’s model to create positive labels at 95% estimated precision, until the number of positive identifications at 95% precision in the validation set ceases to increase (15 iterations, **Fig. S1**). A more detailed and descriptive outline of the FIRE method and its motivation is provided in the **Supplemental Note**.

### Estimated precision of individual FIRE elements

We cannot compute the precision associated with a particular XGBoost score because we do not have access to a set of clean-positive labels. Instead, we define a notion of “estimated precision” using a balanced held-out test set of mixed-positive and negative labels (20% of the data). We defined the estimated precision (*EP*) of a FIRE element to be

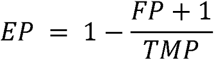

where *TMP* is the number of “true” identifications from the mixed positive labels with at least that element’s XGBoost score, and *FP* is the number of false positive identifications from negative labels with at least that element’s XGBoost score. We add a pseudo count of one to the numerator of false positive identifications so as to prevent liberal estimates for smaller collections of identifications (*25*).

### Aggregate FIRE score calculation

The FIRE score (*S_g_*) for a position in the genome (*g*) is calculated using the following formula:

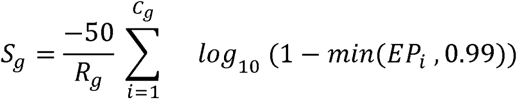

where *C_g_* is the number of FIRE elements at the *g*^th^ position, *R_g_* is the number of Fiber-seq reads at the *g*^th^ position, and *EP_i_* is the estimated precision of the *i*^th^ FIRE element at the *g*^th^ position. The estimated precision of each FIRE element is thresholded at 0.99, such that the FIRE score takes on values between 0 and 100. Regions covered by less than four FIRE elements (i.e., if *C_g_* < 4) are not scored and are given a value of negative one (**Fig. S2**).

### Regions of unreliable coverage

Regions with unreliable coverage were defined as regions with Fiber-seq coverage that deviate from the median coverage by five standard deviations, where a standard deviation is defined by the Poisson distribution (i.e., the square root of the mean coverage).

### FIRE score FDR calculation

We shuffle the location of an entire Fiber-seq read by selecting a random start position within the chromosome and relocating the entire read to that start position. Fiber-seq reads originating from regions with unreliable coverage (defined above) are not shuffled, and reads from regions with reliable coverage are not shuffled into regions with unreliable coverage (bedtools shuffle - chrom -excl, v2.31.0) (*76*). We then compute FIRE scores associated with the shuffled genome. Recalling that the FIRE score for the *g*^th^ position in the genome is denoted *S_g_*, we divide the number of bases that have shuffled FIRE scores above *S_g_* by the number of bases that have un-shuffled FIRE scores above *S_g_*. This provides an estimate of the FDR associated with the FIRE score *S_g_*.

### Peak calling

Peaks are called by identifying FIRE score local-maxima that have FDR values below a 5% threshold and at least 10% actuation (*C_g_* / *R_g_*). Adjacent local maxima that share 50% of the underlying FIRE elements or have 90% reciprocal overlap are merged into a single peak, using the higher of the two local maxima. Then, the start and end positions of the peak are determined by the median start and end positions of the underlying FIRE elements. We also calculated and reported wide peaks by taking the union of the FIRE peaks and all regions below the FDR threshold and then merged the resulting regions that were within one nucleosome (147 bp) of one another.

### Short-read accessibility data and comparisons

DNase I hypersensitivity sites and bigWig tracks for GM12878 were downloaded from the ENCODE data portal. We used accession ENCFF762CRQ for DNase peaks and ENCFF960FMM for the bigWig track (Meuleman et al., 2020). For CTCF CHiP-seq peaks, we used the union of ENCFF356LIU and ENCFF960ZGP (*77*). When intersecting short-read and FIRE peaks, we required a 1 bp overlap, and when measuring the short-read signal within a FIRE peak, we used the maximum values of the short-read signal across the FIRE peak. scATAC-seq data was downloaded in fastq format from the ENCODE portal (**Supplemental Table 5**). Fastq data from each experiment were processed using cellranger-atac count, and the outputs were aggregated using cellranger-atac aggr (10x Genomics v.2.1.0). Aligned fragments from passing cell barcodes were intersected with FIRE peaks using bedmap (v2.4.41) (*78*). Shuffled regions were generated using Bedtools shuffle (v2.31.0). scATAC-seq peaks were called using MACS2 callpeak (v2.2.7.1, parameters: -g hs -q 0.01 --nomodel --shift -75 --extsize 150 --keep-dup all -B --SPMR). scATAC-seq percent accessibility values were computed for each element in the FIRE and shuffled peak sets as the percentage of cell barcodes with at least 1 fragment overlapping that respective peak region.

### FIRE Peaks Enrichment Transcription Factor Motifs

To determine the transcription factor (TF) binding motifs enriched in FIRE peaks we used Homer v3.12 (http://homer.ucsd.edu/homer/motif/) to discover TF binding sites represented in target sequences and compared them to a chosen background. Using findMotifsGenome.pl, we determined the enrichment of all FIRE peaks and all ATAC peaks to the whole genome as a background, then we compared ATAC-overlapping FIRE peaks and non-ATAC-overlapping FIRE peaks with all FIRE peaks as the background. We determined the REST motif genome coordinates in the non-ATAC-overlapping FIRE peaks for aggregate footprinting analysis using annotatePeaks.pl.

### Haplotype-selective peaks

For peaks with at least 10 Fiber-seq reads on both haplotypes, we tested the difference in percent actuation in each haplotype (fraction of reads with FIRE elements) using a two-sided Fisher’s exact test. Specifically, the inputs for the test are the number of FIRE elements in haplotype one, the number of Fiber-seq reads without FIRE elements in haplotype one, the number of FIRE elements in haplotype two, and the number of Fiber-seq reads without FIRE elements in haplotype two. We then apply an FDR correction (Benjamini-Hochberg) to correct for multiple testing and select elements with a correct p-value less than or equal to 0.05 to be haplotype selective peaks.

### The FIRE pipeline

The FIRE pipeline (https://github.com/fiberseq/FIRE v0.0.4) was applied to aligned and phased Fiber-seq BAM files with m6A predictions called using Fibertools-rs (v0.4) (*26*). The FIRE pipeline is a Snakemake (*79*) workflow that applies the FIRE model to individual reads, calculates the aggregate FIRE scores, computes the FDR, calls peaks, and identifies haplotype-selective peaks. The only required inputs for the FIRE pipeline are an aligned Fiber-seq BAM file and the reference genome used for alignment.

### Phasing and identification of genomic variants

Variant calling for single nucleotide variants (SNVs) and insertions/deletions (Indels) was conducted using DeepVariant version 1.5.0 (*80*), while structural variants (SVs) were identified using pbsv (https://github.com/PacificBiosciences/pbsv). The Fiber-seq reads were haplotype-phased through a specialized pipeline available on GitHub (https://github.com/mrvollger/k-mer-variant-phasing) (*16*). This pipeline employs SNVs detected by DeepVariant and utilizes the HiPhase (*81*) variant-based phaser to organize reads into phase blocks. These blocks are then attributed to either the maternal or paternal haplotype using parental short-read genome sequencing in conjunction with the trio k-mer-based phaser meryl (k=31) when available (*82*).

### Identification of CpG methylation

Base-level CpG methylation was called using jasmine (PacBio), and the percent CpG methylation at each genomic position was identified from a pileup of reads using pb-CpG-tools (https://github.com/PacificBiosciences/pb-CpG-tools).

### Imprinted loci

Imprinted loci were defined using the 143 differentially methylated regions identified in 12 B-lymphocyte cell lines by Akbari et al. (*83*). Novel imprinted elements in the 13 fibroblast samples were identified by taking the union of all haplotype-selective peaks across all 13 samples and identifying sites where at least 10 of the samples showed significant chromatin actuation. Subsequently, we analyzed these elements for sites where all samples showed the same direction of maternal or paternal skew and classified these as putative novel imprinted elements.

### Identifying regions enriched in haplotype-selective elements

To test for regions enriched for haplotype-selective elements, we took consecutive windows containing 100 FIRE peaks (sliding by 10 peaks) and compared the number of nominally significant haplotype-selective elements in the 100-peak window to the number of haplotype-selective peaks genome-wide using a two-sided Fisher’s exact test.

### Intra- versus Inter-sample sample similarity in FIRE actuation

To measure the similarity between two haplotypes, we took genomic regions containing 100 peaks and measured the cosine similarity of the percent actuation between the two haplotypes. We then repeated this process across the genome in 100 peak windows, sliding 10 peaks at a time and comparing every unique pair of haplotypes across our two samples.

### Testing for enrichment of inter-sample similarity within SDs and alternative haplotypes

To test for the enrichment of regions with greater inter-sample similarity, we compared the percent of sites intersecting with SDs or alternative haplotypes to 10,000 random shufflings of windows of the same size to calculate an empirical p-value.

### Genome assembly of COLO829BL and estimation of the number of T2T scaffolds

To create a *de novo* assembly for COLO829BL, we used Verkko (v1.4.1) with down-sampled PacBio HiFi reads (60-fold) and ONT ultra-long reads (60-fold) where ultra-long reads were down-sampled in descending order based on read length to retain the longest reads (*4*). Additionally, we used 30-fold Illumina Hi-C data within Verkko to allow for long-range phasing of the assembly. To estimate the number of telomere-to-telomere (T2T) scaffolds, we required that a single scaffold covered more than 95% of a specific chromosome based on T2T-CHM13v2.0 alignment and that there were more than 20 occurrences of telomere sequences with 500bp of both ends of the contig.

### Genome assembly of ST001

To create a de novo assembly for ST001, we used hifiasm (v0.19.9) with down-sampled PacBio HiFi reads selected to retain the longest 70-fold of reads. The initial PacBio HiFi dataset combined the Fiber-seq liver and lung reads from ST001. We then created consistent phasing by aligning the Fiber-seq reads (pbmm2 v1.13.0) back to the diploid assembly of ST001 and assigned haplotypes based on alignment to either the H1 or H2 assembly.

### Plasmid construction

To generate plasmid constructs, the pEF-GFP backbone was linearized by PCR using Repliqa HiFi ToughMix (Quantabio, 95200) with an annealing temperature of 61°C for 30 cycles. The PCR product was purified using a Monarch PCR & DNA Cleanup Kit (NEB, T1130) and treated with DpnI (NEB, R0176) for 1 hour at 37°C to remove methylated template, followed by a second column purification. Gene fragments (gBlocks) encoding the H1 wild-type, 1-edit, 3-edit, and H2 reference sequences were synthesized by IDT with terminal homology arms compatible with the linearized backbone. Constructs were assembled by Gibson Assembly using Gibson Assembly Master Mix (NEB, E2611) at a 3:1 molar ratio of insert to backbone and incubated at 50°C for 1 hour.

### Bacterial transformation and plasmid preparation

Gibson assembly reactions were transformed into NEB 5-alpha competent cells (NEB, C2987) following the manufacturer’s protocol and plated on LB agar supplemented with 100 µg/mL carbenicillin. Single colonies were inoculated into 5 mL LB with 100 µg/mL carbenicillin and grown overnight at 37°C with shaking at 250 rpm. Plasmid DNA was isolated using the Monarch Spin Plasmid Miniprep Kit (NEB, T1110) and sequence-verified by Plasmidsaurus. To remove endogenous m6A methylation, verified plasmids were re-transformed by electroporation into the methyltransferase-deficient strain ER2796 (1.8 kV) and recovered in SOC medium. Transformants were selected on LB with 100 µg/mL carbenicillin, cultured overnight, miniprepped, and sequence-verified as above. Verified constructs were pooled at equimolar concentrations.

### Cell culture and transfection

2×10L HEK293T cells were seeded in a 10 cm dish in 12 mL DMEM (Thermo Fisher, 11965076) supplemented with 10% FBS (Sigma-Aldrich, F0926) and 1% penicillin-streptomycin (Thermo Fisher, 15070063). Twenty-four hours after seeding, cells were transfected with 10 µg of the equimolar plasmid pool using Lipofectamine LTX (Thermo Fisher, 15338100) in Opti-MEM (Thermo Fisher, 31985070) according to the manufacturer’s protocol. Forty-eight hours post-transfection, cells were harvested by trypsinization (0.25% trypsin-EDTA; Thermo Fisher, 25200072), counted, and pelleted at 350 × *g* for 5 minutes.

### Plasmid Fiber-seq

Cell pellets were resuspended in Buffer A (15 mM Tris-HCl pH 8.0, 15 mM NaCl, 60 mM KCl, 1 mM EDTA pH 8.0, 0.5 mM EGTA pH 8.0, 0.5 mM spermidine) at a concentration of 2×10L cells per 60 µL. Nuclei were extracted by addition of an equal volume of 2x lysis buffer (Buffer A supplemented with 0.05% IGEPAL CA-630) and incubated on ice for 10 minutes. Nuclei were pelleted at 350 × *g* for 5 minutes at 4°C, resuspended in 60 µL Fiber-seq Master Mix (Buffer A, 0.8 mM SAM, 200 U Hia5), and incubated at 25°C for 10 minutes. The reaction was quenched by addition of SDS to a final concentration of 1% (*38*).

### Plasmid DNA isolation and library preparation

Methylated plasmid DNA was isolated from treated nuclei using the Monarch Plasmid Miniprep Kit and eluted in 35 µL water. Contaminating genomic DNA was removed by treatment with Exonuclease V (NEB, M0345) at 37°C for 1 hour, followed by heat inactivation at 70°C for 30 minutes and column purification. Plasmids were linearized with KpnI-HF (NEB, R3142), and sequencing libraries were prepared using the SMRTbell Prep Kit 3.0 (PacBio) according to the manufacturer’s protocol. Libraries were sequenced on a PacBio Revio at the University of Washington PacBio Sequencing Services.

### Differential nucleosome occupancy analysis

To compare chromatin architecture across plasmid constructs, we performed pairwise differential footprint analyses on Fiber-seq data from four plasmid samples (H1_wild, H2_ref, 1_edit, 3_edit, **Supplemental Table 4**). Each sample was processed using fibertools to obtain m6A methylation data, followed by FiberHMM to call nucleosome footprints (≥80 bp) on individual molecules. Reads were aligned to their respective plasmid reference sequences using pbmm2.

To normalize for chromatin state, reads were subsampled across all four samples to a shared nucleosome-per-read (NPR) distribution by taking the minimum read count at each NPR value. Per-position nucleosome occupancy was then computed as the fraction of subsampled reads with a nucleosome footprint overlapping each base in the plasmid sequence. To account for a 3 bp internal deletion in H1_wild relative to the other constructs, the corresponding positions were removed from the H1_wild occupancy array to restore positional alignment. To identify regions of differential nucleosome positioning, we subtracted the per-position occupancy of each sample from H2_ref (the reference haplotype), and separately compared H1_wild, 1_edit, and 3_edit to one another to assess equivalence among constructs carrying the functional variant.

### X chromosome peaks

FIRE peaks used in XCI analyses were filtered to remove peaks overlapping ENCODE 2020 hg38 blacklist regions (accession: ENCFF356LFX). Transcriptional start site (TSS) FIRE peaks were identified as peaks intersecting both GENCODE v45 TSSs (padded to 20 bp) and ENCODE CAGE-seq peaks (**Supplemental Table 5**). CTCF FIRE peaks were identified as peaks intersecting ENCODE CTCF ChIP-seq peaks (**Supplemental Table 5**). Intersections were performed using Bedtools intersect (v2.31.0). Peaks included in XCI analyses were filtered to meet the following phasing and coverage criteria: <= 25% variance from the mean coverage, < 35% of reads assigned as unphased, and at least 10X coverage from each haplotype. In addition, GM12878 peaks surrounding the MTMR1 TSS (chrX:150,631,401-150,738,995) and a fibroblast sample peak at the PDHA1 TSS (chrX:19,343,661- 19,343,996) were manually filtered due to substantial disagreement in between the variant and k-mer phasing methods. Peaks were required to be actuated on at least 30% of Fiber-seq reads on the Xa and/or Xi.

### XCI Classifications

X chromosome FIRE peaks with a percent actuation difference of >= 50% between haplotypes were classified as “Xa-specific” or “Xi-specific” if the peaks were actuated on >= 30% of paternal or maternal Fiber-seq reads, respectively. FIRE Peaks were classified as “shared” if the peaks were actuated on >= 30% of Fiber-seq reads on both haplotypes or >= 30% of reads on one haplotype with a percent difference of < 50% between haplotypes. Transcriptional start site (TSS) FIRE peaks actuated on less than 30% of maternal Fiber-seq reads were classified as inactivated. TSS FIRE peaks were classified as fully escaping XCI if >= 30% of maternal Fiber-seq reads were actuated and the percent actuation difference between haplotypes was less than 25%.

### Iso-seq analysis

Full-length 3’ transcriptomic iso-seq data was generated in a previous study (*16*). Reads were clustered by isoform using Isoseq cluster 2 (PacBio v3.99.99) and aligned to hg38 using pbmm2 (PacBio v1.11.99, parameters: --sort --min-gap-comp-id-perc 95.0 --min-length 50 --sample --report-json mapping_stats.report.json --preset ISOSEQ). Phased variant calls were used to haplotag clustered isoforms using WhatsHap v1.6 (*84*). Clustered isoforms were then collapsed using Isoseq collapse v4.0.0 (PacBio). Isoforms were annotated using Pigeon v1.2.0 (PacBio). Isoforms and genes with >= 35% of transcripts not assigned to a haplotype were classified as unphased.

### Promoter-proximal escape quantification

TSS FIRE peaks were grouped by escape status (fully escaping or inactivated). For each group, non-TSS FIRE peaks within 100 Kb in either direction (upstream and downstream) of each TSS were counted in 5 Kb distance bins. Counts within each bin were then normalized by the number of TSSs in the respective group.

### CTCF footprinting

X chromosome CTCF FIRE peaks (overlapping ENCODE CTCF ChIP-seq peaks) that also fully overlap a FIMO-predicted CTCF motif (*85*) (v5.5) were used for footprinting analyses. We decided to limit CTCF footprinting to the most highly bound portions of CTCF, modules two and three (*86*). FIRE elements that completely overlapped modules two and three of a motif were centered using *fibertools* ft center (*26*), and the number of m6A observed within the motif was quantified. FIRE-contained motifs with <= 1 m6A were classified as bound.

## Data availability

Processed epigenetic (FIRE) results for all samples are publicly available through https://doi.org/10.5281/zenodo.14511246, as well as the Fiber-seq data portal https://stergachislab.github.io/Fiber-seq-publication-data/. Per-sample information is provided in **Supplemental Table 6**. All raw and processed sequencing data generated for GM12878 (PRJNA1233341) and K562 (SRX20077598, SRX20077599, and SRX20077600) have been submitted to the NCBI Sequence Read Archive (https://www.ncbi.nlm.nih.gov/sra/). Restrictions apply to the availability of some genetic data generated or analyzed during this study to preserve subject confidentiality. Genomic data for these samples are accessible to the scientific community through the UDN, GREGoR, All of US, and SMaHT consortia, with per-sample information and accessions provided in **Supplemental Table 6**. Individuals interested in accessing this should submit a data access request to the relevant consortia. Cell lines obtained from the National Institute of General Medical Sciences Human Genetic Cell Repository at the Coriell Institute for Medical Research include GM12878.

## Code availability

FIRE is available as a Snakemake pipeline on GitHub (https://github.com/fiberseq/FIRE) and Zenodo (https://zenodo.org/records/11075226). Fibertools is available on GitHub (https://github.com/fiberseq/fibertools-rs) and Zenodo (https://zenodo.org/records/10850620). The code used for phasing long-read data is available as a Snakemake pipeline on GitHub (https://github.com/mrvollger/k-mer-variant-phasing) and Zenodo (https://zenodo.org/records/10655527). The code for making figures and tables is available on GitHub (https://github.com/mrvollger/fire-figures) and Zenodo (https://zenodo.org/doi/10.5281/zenodo.10681988).

## Acknowledgments

The authors thank Dr. John Stamatoyannopoulos for his assistance in reviewing this manuscript and Christine M. Disteche for feedback regarding the XCI escape portion of this study. We thank Alan Beggs, Monica Wojcik, Anne O’Donnell Luria, Gail Jarvik, and Katrina Dipple for providing the fibroblast cell lines. The following cell lines/DNA samples were obtained from the NIGMS Human Genetic Cell Repository at the Coriell Institute for Medical Research: GM12878. A.B.S. holds a Career Award for Medical Scientists from the Burroughs Wellcome Fund and is a Pew Biomedical Scholar. This study was supported by National Institutes of Health (NIH) grants 1DP5OD029630, UM1DA058220, 1U01HG013744, and OT2OD002748 to A.B.S., a Brotman Baty Institute Catalytic Collaboration Grant to A.B.S., NIH grants 1DP2AI183504 and 1U01AI176320 to J.P.R., as well as a Crohn’s and Colitis Foundation Senior Research Award #1158945 to J.P.R. This study was also possible thanks to support from Google.org. Additionally, M.R.V. and S.C.B. were supported by a training grant (T32) from the NIH (2T32GM007454-46). M.R.V. was also supported by a Pathway to Independence award from the National Institute of General Medical Sciences (1K99GM155552-01).

## Author contributions

Conceptualization and design: M.R.V., E.G.S., and A.B.S. Experimental design and execution: E.G.S., J.R., B.J.M. and A.B.S. Computational experiments: M.R.V., E.G.S., D.D., B.J.M., A.E.S., and S.J.N. Data generation: E.G.S., J.R., K.M.M., C.H., S.C.B., Y.M., N.L.P., B.J.M., D.G.M, G.H.G, K.H., J.G.M., M.C., E.E.E, J.T.B., D.J.M. and A.B.S. Genome Assembly: W.T.H., Y.K., and E.E.E. Conceptualization and design of the semi-supervised training regime and peak calling: M.R.V., D.M.W., W.E.F., W.S.N., and A.B.S. FIRE implementation: M.R.V. Supplemental material organization: M.R.V., E.G.S., and A.B.S. Display items: M.R.V., E.G.S., and A.B.S. Manuscript writing: M.R.V., E.G.S., and A.B.S. with input from all authors.

