## Supplemental Figures for "Somatic epimutations cap genetic determinism in the human diploid chromatin epigenome"

### Figure S1

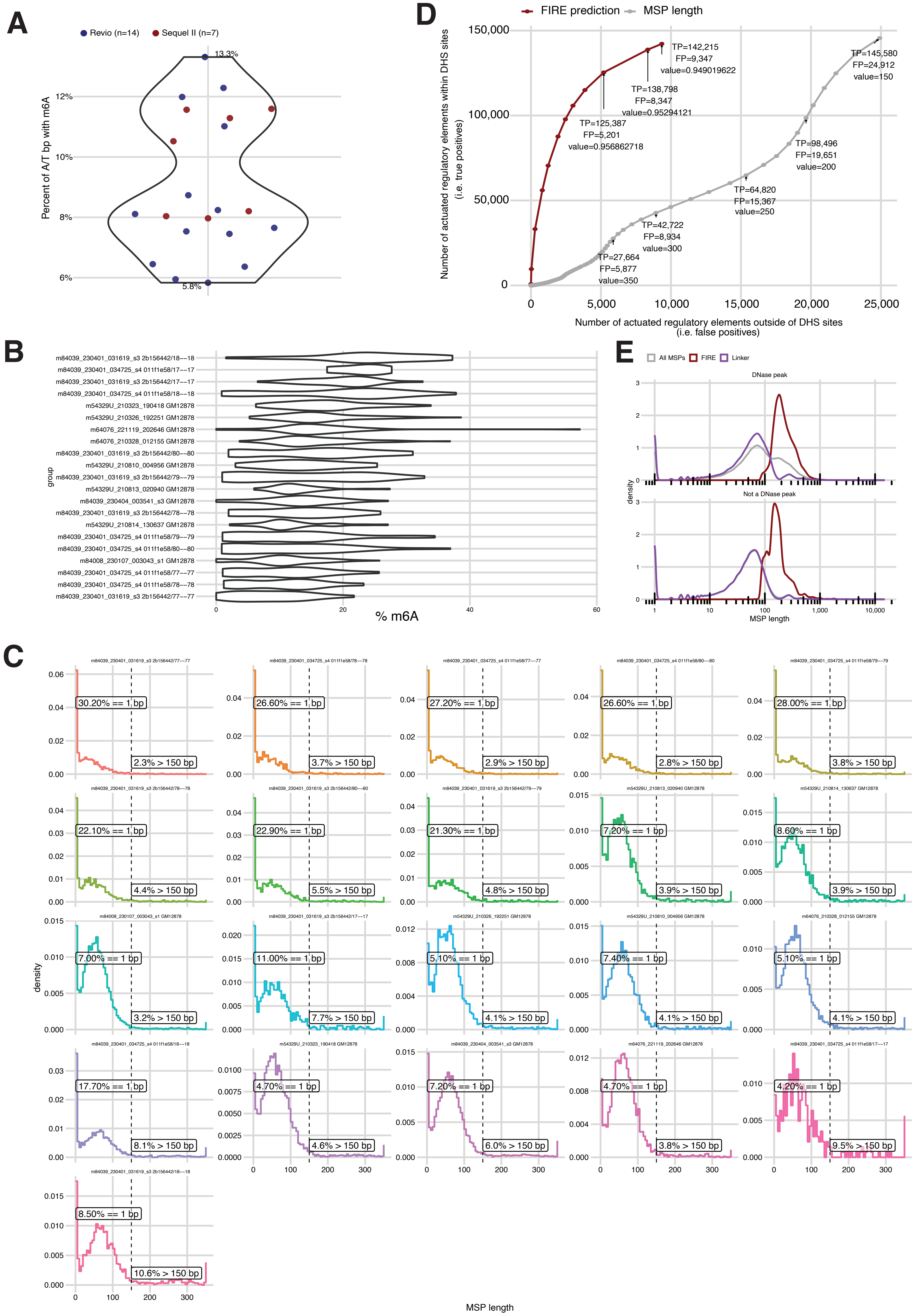
A
Revio (n=14)
Sequel II (n=7)
Percent of A/T bp with m6A
13.3%
5.8%
B
% m6A
group
C
density
MSP length
D
FIRE prediction
MSP length
Number of actuated regulatory elements within DHS sites (i.e. true positives)
Number of actuated regulatory elements outside of DHS sites (i.e. false positives)
TP=142,215 FP=9,347 value=0.949019622
TP=138,798 FP=8,347 value=0.95294121
TP=125,387 FP=5,201 value=0.956862718
TP=145,580 FP=24,912 value=150
TP=98,496 FP=19,651 value=200
TP=64,820 FP=15,367 value=250
TP=42,722 FP=8,934 value=300
TP=27,664 FP=5,877 value=350
E
All MSPs
FIRE
Linker
DNase peak
Not a DNase peak
density
MSP length

### Figure S2

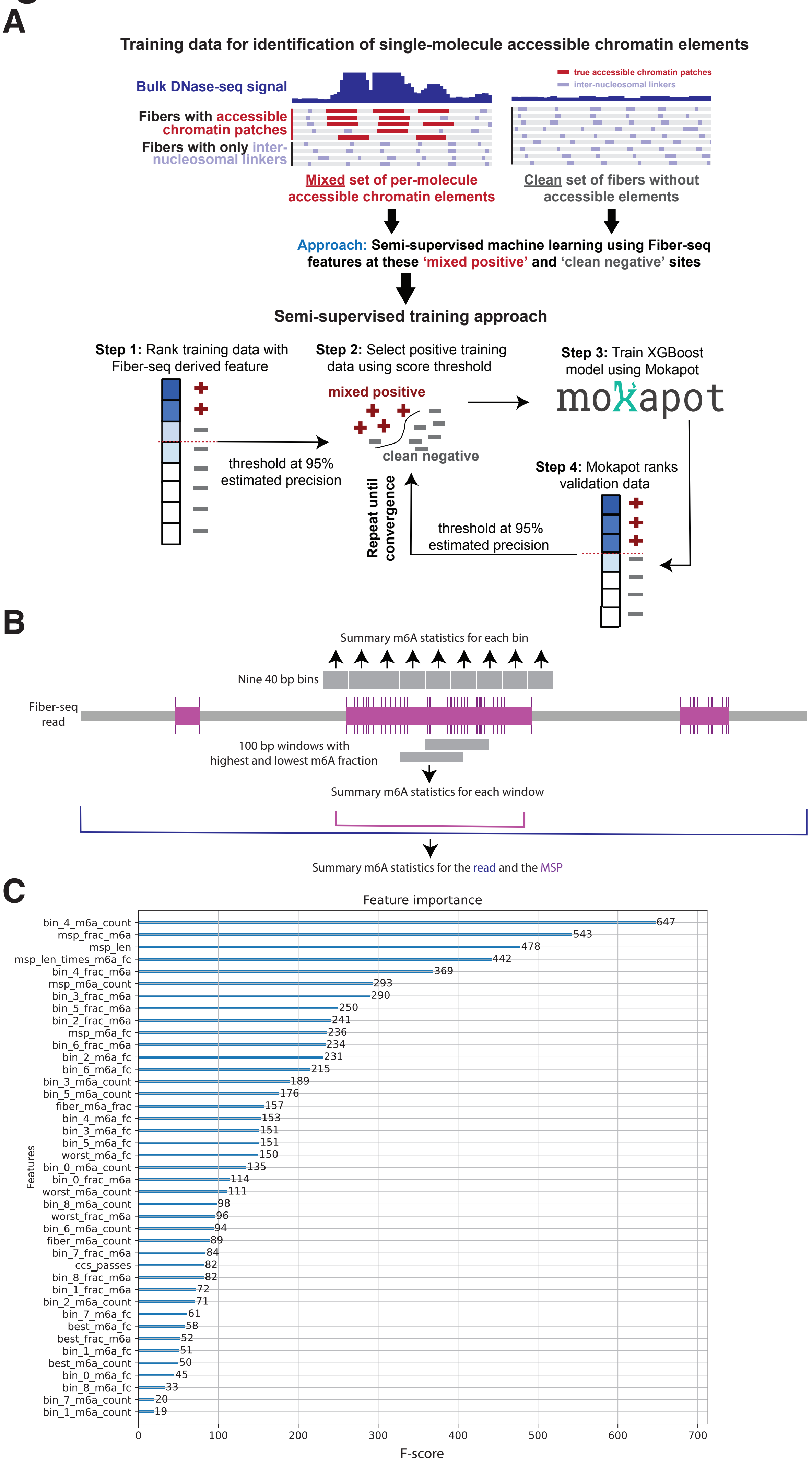
A
Training data for identification of single-molecule accessible chromatin elements
Bulk DNase-seq signal
true accessible chromatin patches
inter-nucleosomal linkers
Fibers with accessible chromatin patches
Fibers with only inter-nucleosomal linkers
Mixed set of per-molecule accessible chromatin elements
Clean set of fibers without accessible elements
Approach: Semi-supervised machine learning using Fiber-seq features at these 'mixed positive' and 'clean negative' sites
Semi-supervised training approach
Step 1: Rank training data with Fiber-seq derived feature
threshold at 95% estimated precision
Step 2: Select positive training data using score threshold
mixed positive
clean negative
Step 3: Train XGBoost model using Mokapot
mokapot
Step 4: Mokapot ranks validation data
threshold at 95% estimated precision
Repeat until convergence
B
Summary m6A statistics for each bin
Nine 40 bp bins
Fiber-seq read
100 bp windows with highest and lowest m6A fraction
Summary m6A statistics for each window
Summary m6A statistics for the read and the MSP
C
Feature importance
bin_4_m6a_count 647
msp_frac_m6a 543
msp_len 478
msp_len_times_m6a_fc 442
bin_4_frac_m6a 369
msp_m6a_count 293
bin_3_frac_m6a 290
bin_5_frac_m6a 250
bin_2_frac_m6a 241
msp_m6a_fc 236
bin_6_frac_m6a 234
bin_2_m6a_fc 231
bin_6_m6a_fc 215
bin_3_m6a_count 189
bin_5_m6a_count 176
fiber_m6a_frac 157
bin_4_m6a_fc 153
bin_3_m6a_fc 151
bin_5_m6a_fc 151
worst_m6a_fc 150
bin_0_m6a_count 135
bin_0_frac_m6a 114
worst_m6a_count 111
bin_8_m6a_count 98
worst_frac_m6a 96
bin_6_m6a_count 94
fiber_m6a_count 89
bin_7_frac_m6a 84
ccs_passes 82
bin_8_frac_m6a 82
bin_1_frac_m6a 72
bin_2_m6a_count 71
bin_7_m6a_fc 61
best_m6a_fc 58
best_frac_m6a 52
bin_1_m6a_fc 51
best_m6a_count 50
bin_0_m6a_fc 45
bin_8_m6a_fc 33
bin_7_m6a_count 20
bin_1_m6a_count 19
Features
0
100
200
300
400
500
600
700
F-score

### Figure S3

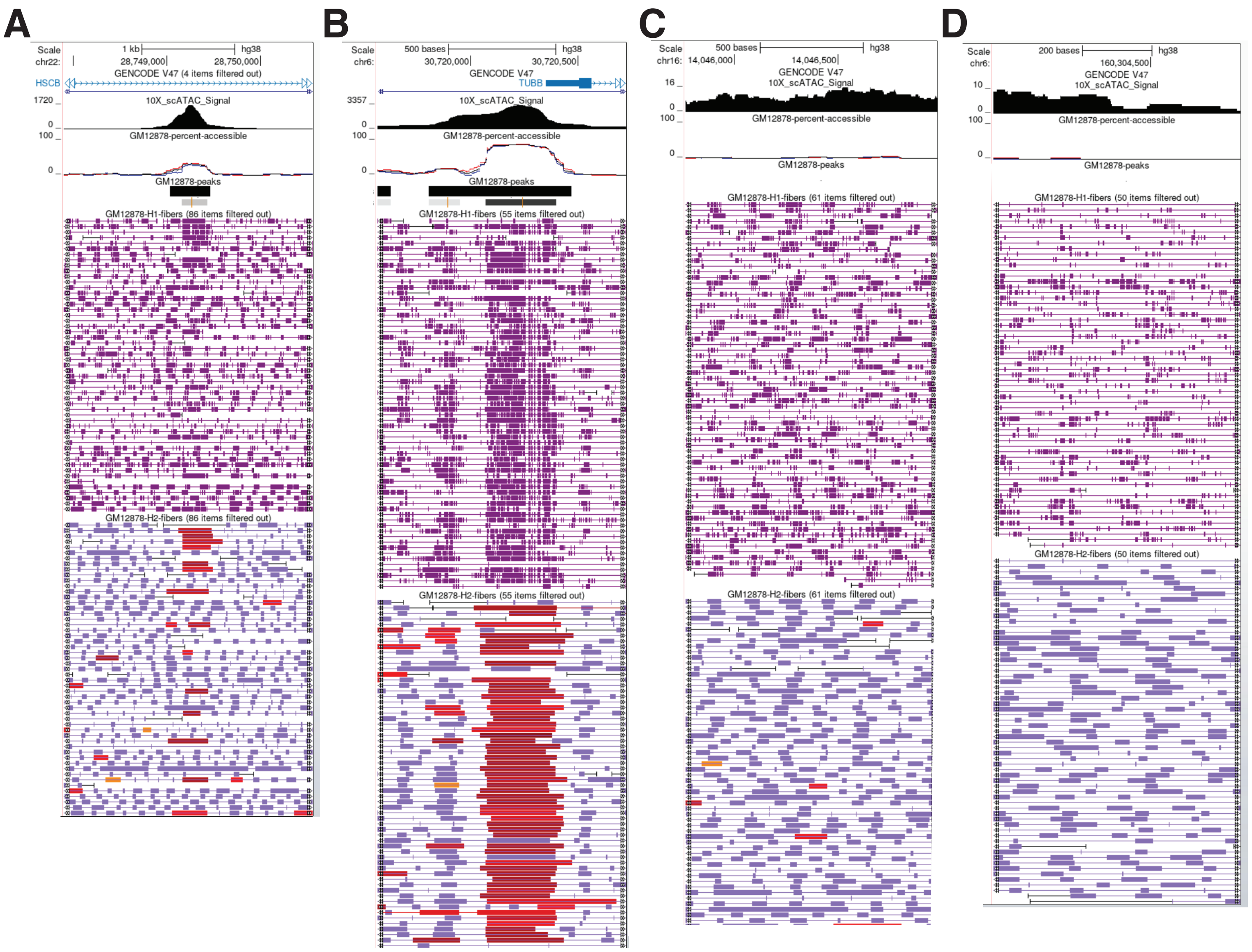
A
Scale 1 kb hg38
chr22: 28,749,000 28,750,000
GENCODE V47 (4 items filtered out)
HSCB
1720 _ 10X_scATAC_Signal
0 _
100 _ GM12878-percent-accessible
0 _
GM12878-peaks
GM12878-H1-fibers (86 items filtered out)
GM12878-H2-fibers (86 items filtered out)
B
Scale 500 bases hg38
chr6: 30,720,000 30,720,500
GENCODE V47
TUBB
3357 _ 10X_scATAC_Signal
0 _
100 _ GM12878-percent-accessible
0 _
GM12878-peaks
GM12878-H1-fibers (55 items filtered out)
GM12878-H2-fibers (55 items filtered out)
C
Scale 500 bases hg38
chr16: 14,046,000 14,046,500
GENCODE V47
16 _ 10X_scATAC_Signal
0 _
100 _ GM12878-percent-accessible
0 _
GM12878-peaks
GM12878-H1-fibers (61 items filtered out)
GM12878-H2-fibers (61 items filtered out)
D
Scale 200 bases hg38
chr6: 160,304,500
GENCODE V47
10 _ 10X_scATAC_Signal
0 _
100 _ GM12878-percent-accessible
0 _
GM12878-peaks
GM12878-H1-fibers (50 items filtered out)
GM12878-H2-fibers (50 items filtered out)

### Figure S4

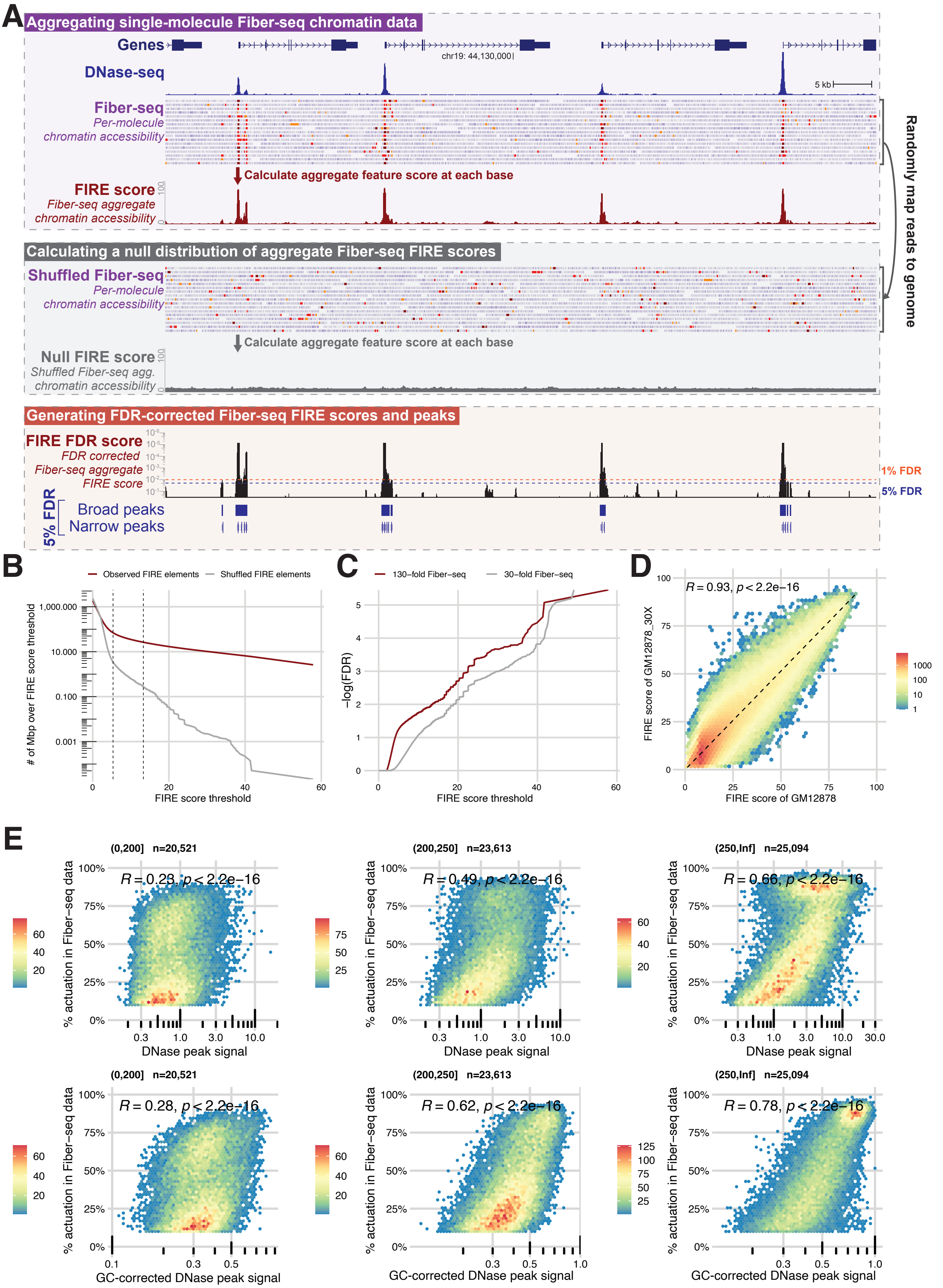

A
Aggregating single-molecule Fiber-seq chromatin data
Genes
chr19: 44,130,000
DNase-seq
5 kb
Fiber-seq
Per-molecule chromatin accessibility
Calculate aggregate feature score at each base
FIRE score
Fiber-seq aggregate chromatin accessibility
Randomly map reads to genome
Calculating a null distribution of aggregate Fiber-seq FIRE scores
Shuffled Fiber-seq
Per-molecule chromatin accessibility
Calculate aggregate feature score at each base
Null FIRE score
Shuffled Fiber-seq agg. chromatin accessibility
Generating FDR-corrected Fiber-seq FIRE scores and peaks
FIRE FDR score
FDR corrected Fiber-seq aggregate FIRE score
1% FDR
5% FDR
5% FDR
Broad peaks
Narrow peaks
B
Observed FIRE elements
Shuffled FIRE elements
### of Mbp over FIRE score threshold
FIRE score threshold
C
130–fold Fiber–seq
30–fold Fiber–seq
−log(FDR)
FIRE score threshold
D
R = 0.93, p < 2.2e−16
FIRE score of GM12878_30X
FIRE score of GM12878
E
(0,200] n=20,521
R = 0.23, p < 2.2e−16
(200,250] n=23,613
R = 0.49, p < 2.2e−16
(250,Inf] n=25,094
R = 0.66, p < 2.2e−16
% actuation in Fiber−seq data
DNase peak signal
(0,200] n=20,521
R = 0.28, p < 2.2e−16
(200,250] n=23,613
R = 0.62, p < 2.2e−16
(250,Inf] n=25,094
R = 0.78, p < 2.2e−16
GC-corrected DNase peak signal

### Figure S5

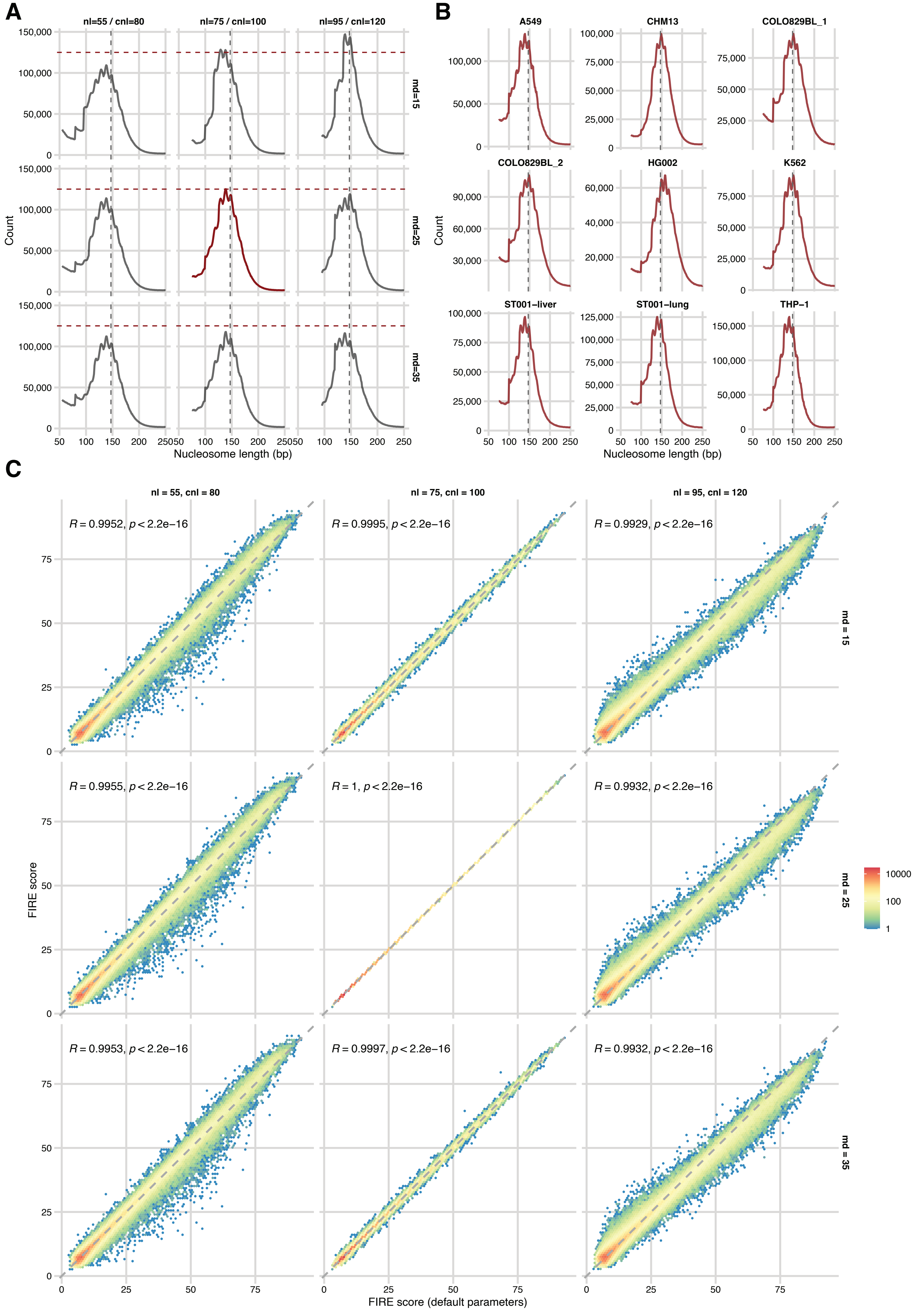
A
nl=55 / cnl=80
nl=75 / cnl=100
nl=95 / cnl=120
md=15
md=25
md=35
Count
Nucleosome length (bp)
B
A549
CHM13
COLO829BL_1
COLO829BL_2
HG002
K562
ST001−liver
ST001−lung
THP−1
Count
Nucleosome length (bp)
C
nl = 55, cnl = 80
nl = 75, cnl = 100
nl = 95, cnl = 120
R = 0.9952, p < 2.2e−16
R = 0.9995, p < 2.2e−16
R = 0.9929, p < 2.2e−16
R = 0.9955, p < 2.2e−16
R = 1, p < 2.2e−16
R = 0.9932, p < 2.2e−16
R = 0.9953, p < 2.2e−16
R = 0.9997, p < 2.2e−16
R = 0.9932, p < 2.2e−16
md = 15
md = 25
md = 35
FIRE score
FIRE score (default parameters)
10000
100
1

### Figure S6

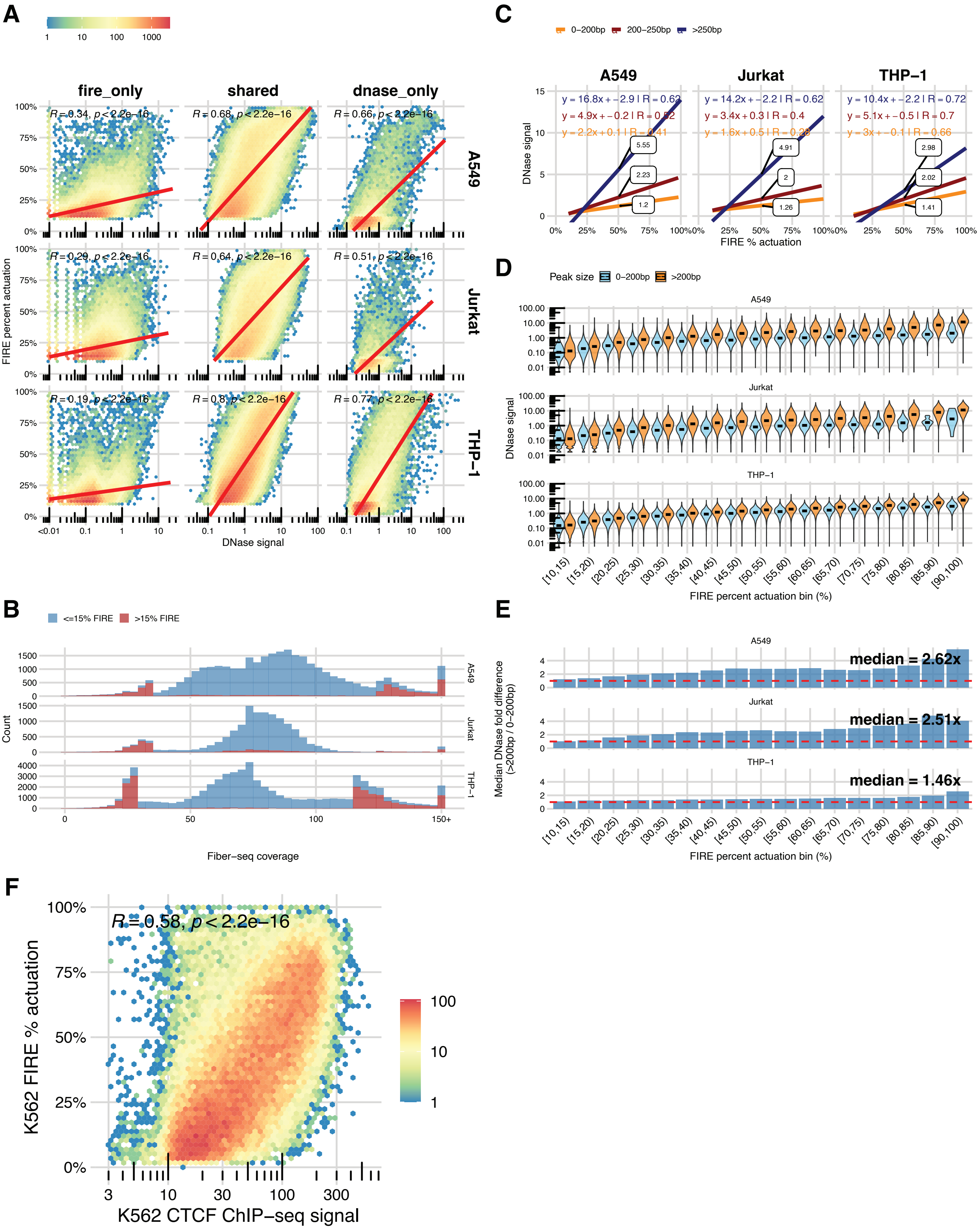

A
B
C
D
E
F
fire_only
shared
dnase_only
A549
Jurkat
THP−1
FIRE percent actuation
DNase signal
<=15% FIRE
>15% FIRE
Count
Fiber−seq coverage
0−200bp
200−250bp
>250bp
FIRE % actuation
Peak size
FIRE percent actuation bin (%)
Median DNase fold difference (>200bp / 0−200bp)
median = 2.62x
median = 2.51x
median = 1.46x
R = 0.58, p < 2.2e−16
K562 FIRE % actuation
K562 CTCF ChIP−seq signal

### Figure S7

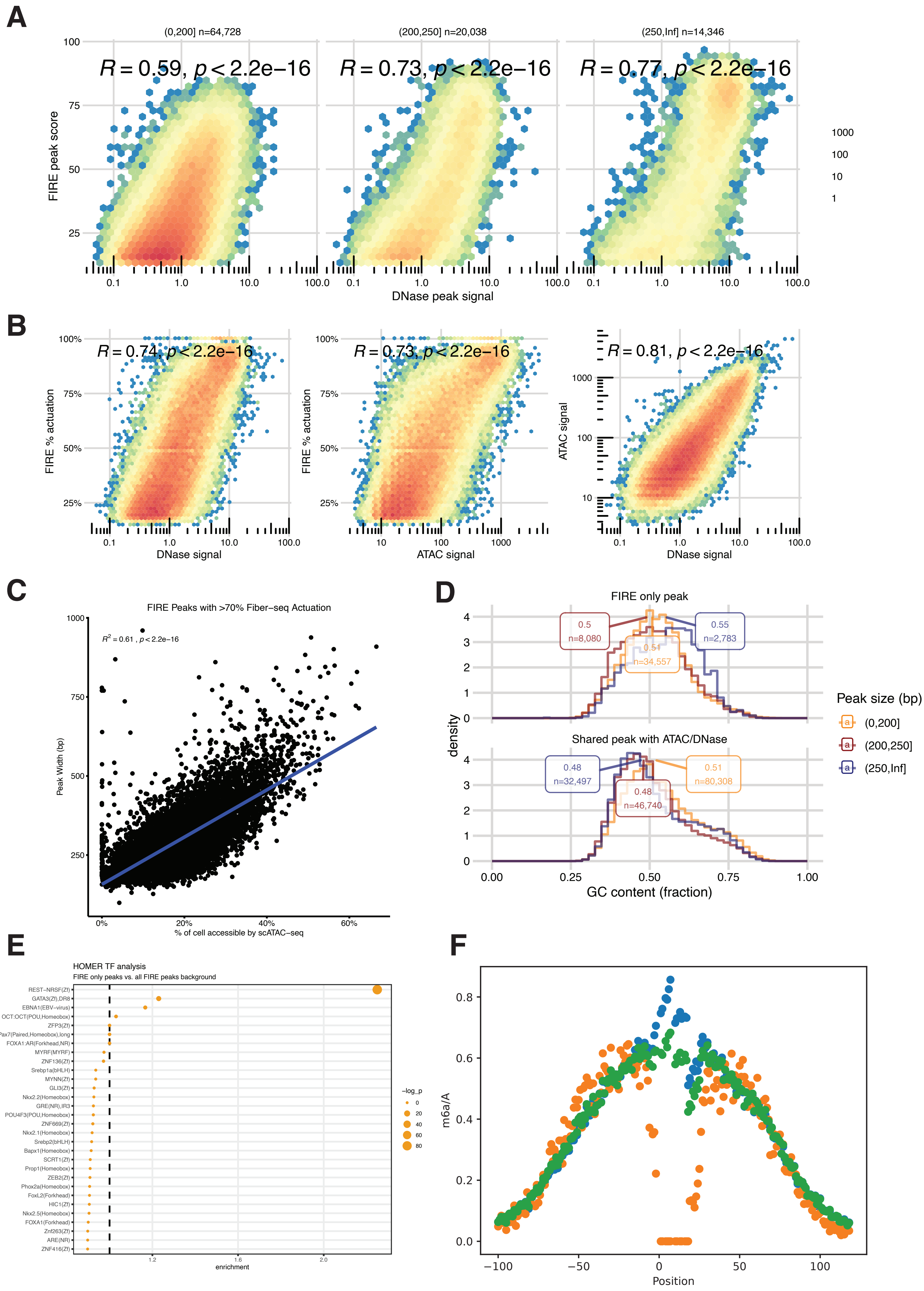

A
(0,200] n=64,728
(200,250] n=20,038
(250,Inf] n=14,346
R = 0.59, p < 2.2e−16
R = 0.73, p < 2.2e−16
R = 0.77, p < 2.2e−16
FIRE peak score
DNase peak signal
1000
100
10
1
B
R = 0.74, p < 2.2e−16
R = 0.73, p < 2.2e−16
R = 0.81, p < 2.2e−16
FIRE % actuation
DNase signal
ATAC signal
C
FIRE Peaks with >70% Fiber−seq Actuation
R² = 0.61 , p < 2.2e−16
Peak Width (bp)
% of cell accessible by scATAC−seq
D
FIRE only peak
Shared peak with ATAC/DNase
0.5 n=8,080
0.55 n=2,783
0.51 n=34,557
0.48 n=32,497
0.51 n=80,308
0.48 n=46,740
density
GC content (fraction)
Peak size (bp)
(0,200]
(200,250]
(250,Inf]
E
HOMER TF analysis
FIRE only peaks vs. all FIRE peaks background
REST−NRSF(Zf)
GATA3(Zf),DR8
EBNA1(EBV−virus)
OCT:OCT(POU,Homeobox)
ZFP3(Zf)
Pax7(Paired,Homeobox),long
FOXA1:AR(Forkhead,NR)
MYRF(MYRF)
ZNF136(Zf)
Srebp1a(bHLH)
MYNN(Zf)
GLI3(Zf)
Nkx2.2(Homeobox)
GRE(NR),IR3
POU4F3(POU,Homeobox)
ZNF669(Zf)
Nkx2.1(Homeobox)
Srebp2(bHLH)
Bapx1(Homeobox)
SCRT1(Zf)
Prop1(Homeobox)
ZEB2(Zf)
Phox2a(Homeobox)
FoxL2(Forkhead)
HIC1(Zf)
Nkx2.5(Homeobox)
FOXA1(Forkhead)
Znf263(Zf)
ARE(NR)
ZNF416(Zf)
TF
enrichment
−log_p
F
m6a/A
Position

### Figure S8

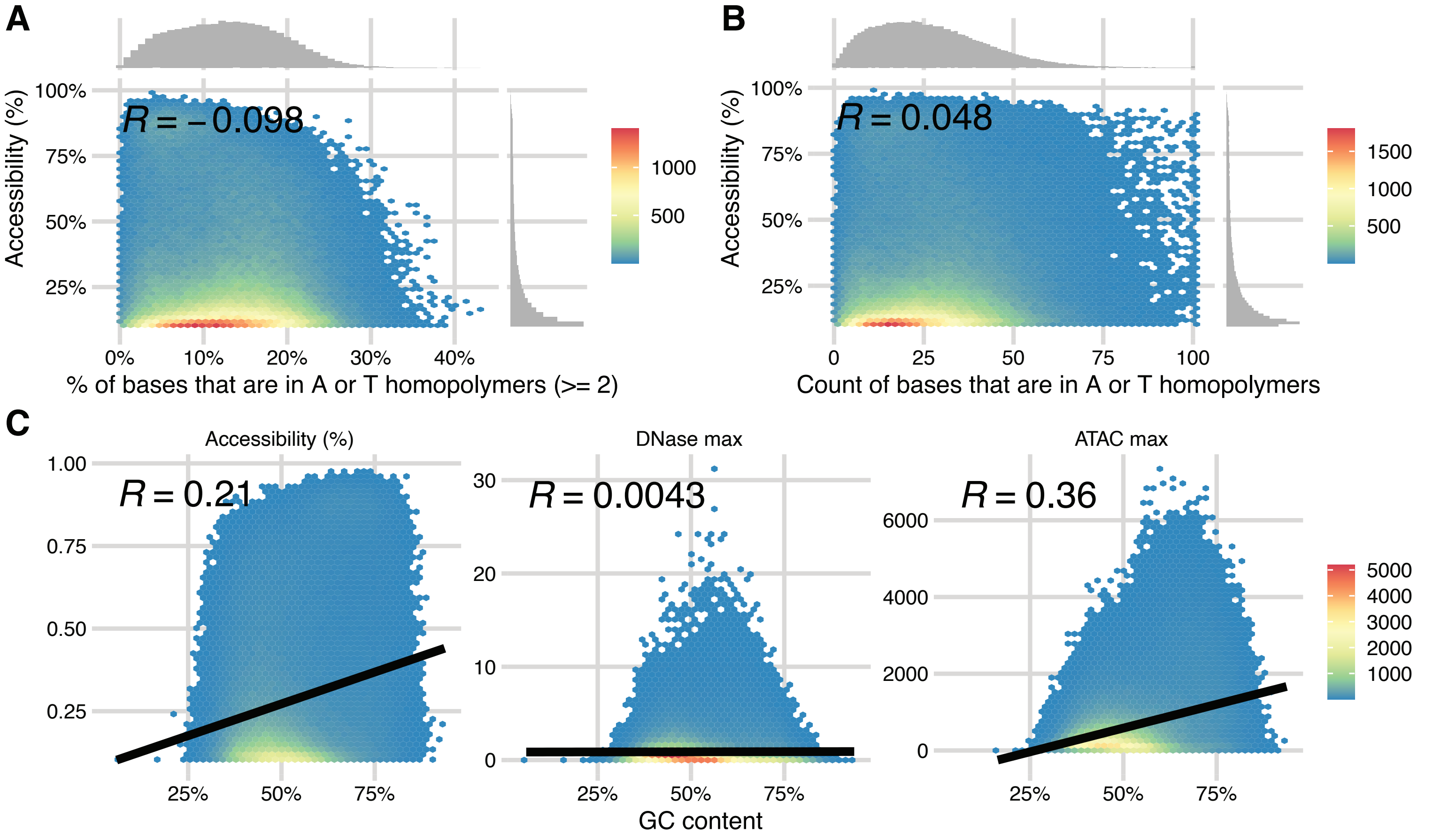

A
B
C
R = −0.098
R = 0.048
Accessibility (%)
100%
75%
50%
25%
0%
10%
20%
30%
40%
% of bases that are in A or T homopolymers (>= 2)
1000
500
0
25
50
75
100
Count of bases that are in A or T homopolymers
1500
Accessibility (%)
DNase max
ATAC max
R = 0.21
R = 0.0043
R = 0.36
1.00
0.75
0.50
0.25
30
20
10
0
6000
4000
2000
GC content
5000
4000
3000
2000
1000

### Figure S9

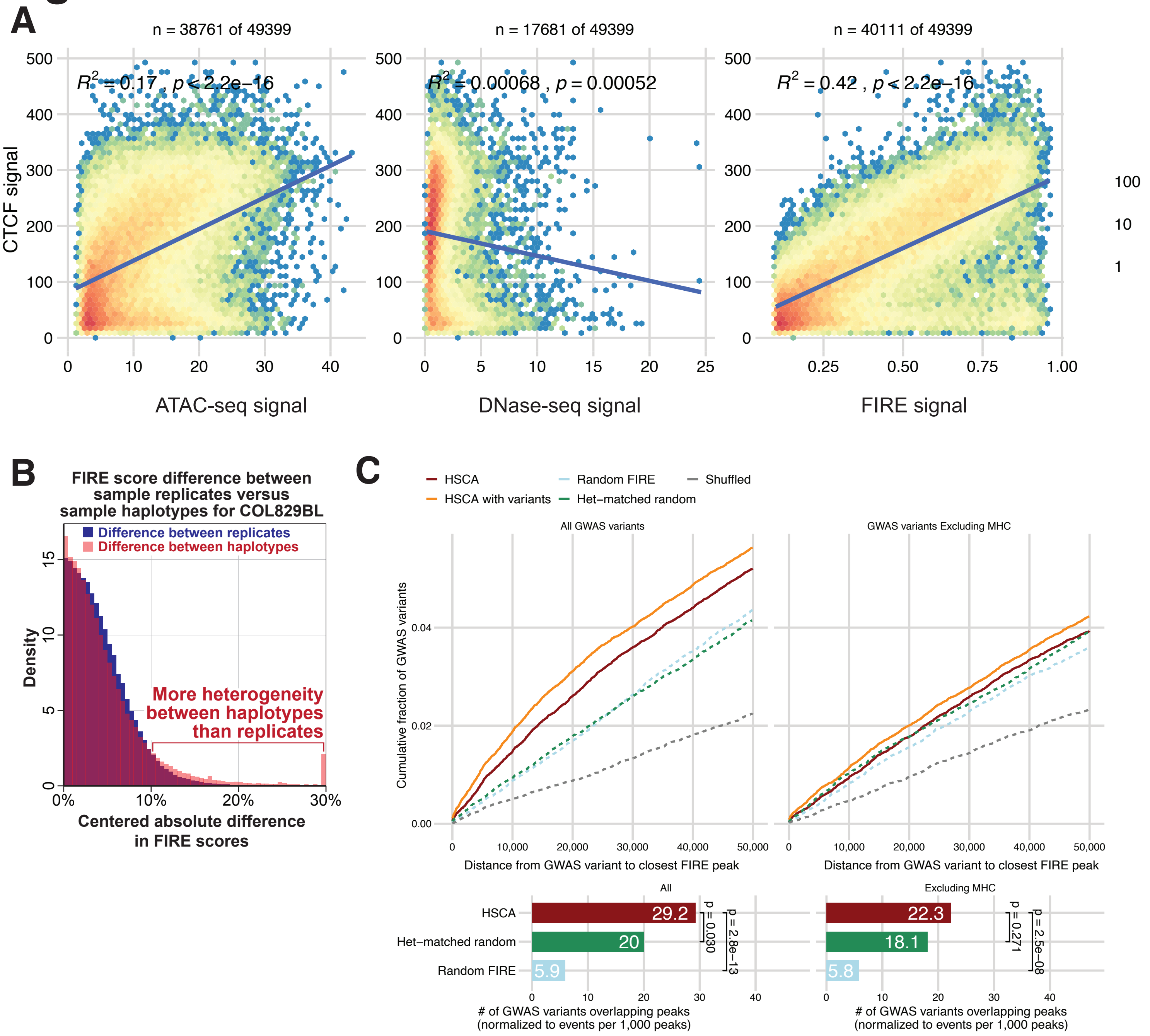
A
n = 38761 of 49399
n = 17681 of 49399
n = 40111 of 49399
$R^2 = 0.17$ , $p < 2.2e-16$
$R^2 = 0.00068$ , $p = 0.00052$
$R^2 = 0.42$ , $p < 2.2e-16$
CTCF signal
ATAC-seq signal
DNase-seq signal
FIRE signal
100
10
1
B
FIRE score difference between sample replicates versus sample haplotypes for COL829BL
Difference between replicates
Difference between haplotypes
Density
More heterogeneity between haplotypes than replicates
Centered absolute difference in FIRE scores
C
HSCA
HSCA with variants
Random FIRE
Het−matched random
Shuffled
All GWAS variants
GWAS variants Excluding MHC
Cumulative fraction of GWAS variants
Distance from GWAS variant to closest FIRE peak
All
Excluding MHC
HSCA
Het−matched random
Random FIRE
29.2
20
5.9
22.3
18.1
5.8
p = 0.030
p = 2.8e−13
p = 0.271
p = 2.5e−08
### of GWAS variants overlapping peaks (normalized to events per 1,000 peaks)

### Figure S10

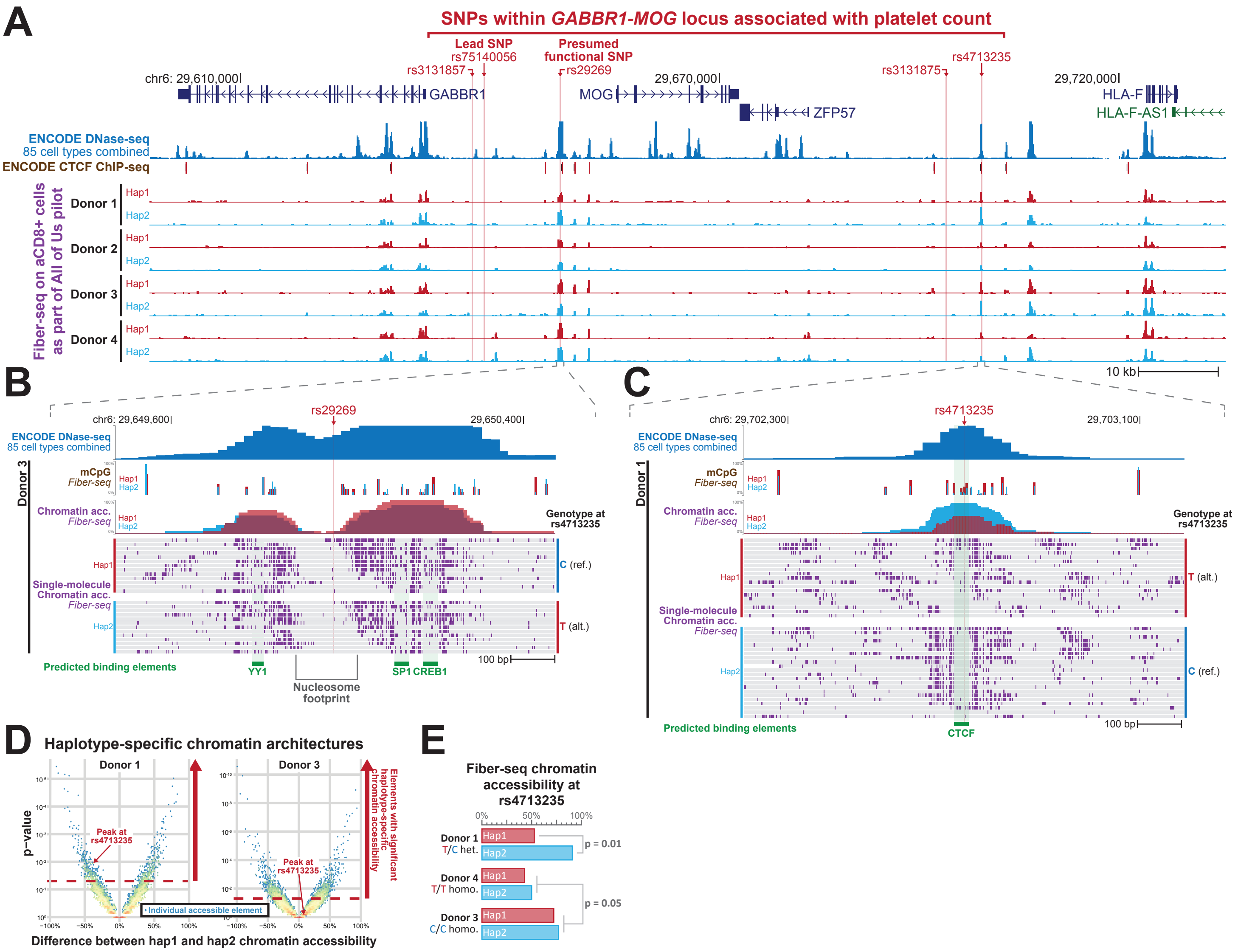

A
SNPs within GABBR1-MOG locus associated with platelet count
Lead SNP rs75140056
Presumed functional SNP
rs3131857
rs29269
rs3131875
rs4713235
chr6: 29,610,000
29,670,000
29,720,000
GABBR1
MOG
ZFP57
HLA-F
HLA-F-AS1
ENCODE DNase-seq
85 cell types combined
ENCODE CTCF ChIP-seq
Fiber-seq on aCD8+ cells as part of All of Us pilot
Donor 1
Donor 2
Donor 3
Donor 4
Hap1
Hap2
10 kb
B
rs29269
chr6: 29,649,600
29,650,400
mCpG Fiber-seq
Chromatin acc. Fiber-seq
Genotype at rs4713235
C (ref.)
T (alt.)
Single-molecule Chromatin acc. Fiber-seq
Predicted binding elements
YY1
Nucleosome footprint
SP1
CREB1
100 bp
C
rs4713235
chr6: 29,702,300
29,703,100
T (alt.)
C (ref.)
CTCF
D
Haplotype-specific chromatin architectures
Donor 1
Donor 3
p-value
Peak at rs4713235
Individual accessible element
Elements with significant haplotype-specific chromatin accessibility
Difference between hap1 and hap2 chromatin accessibility
E
Fiber-seq chromatin accessibility at rs4713235
Donor 1 T/C het.
Donor 4 T/T homo.
Donor 3 C/C homo.
p = 0.01
p = 0.05

### Figure S11

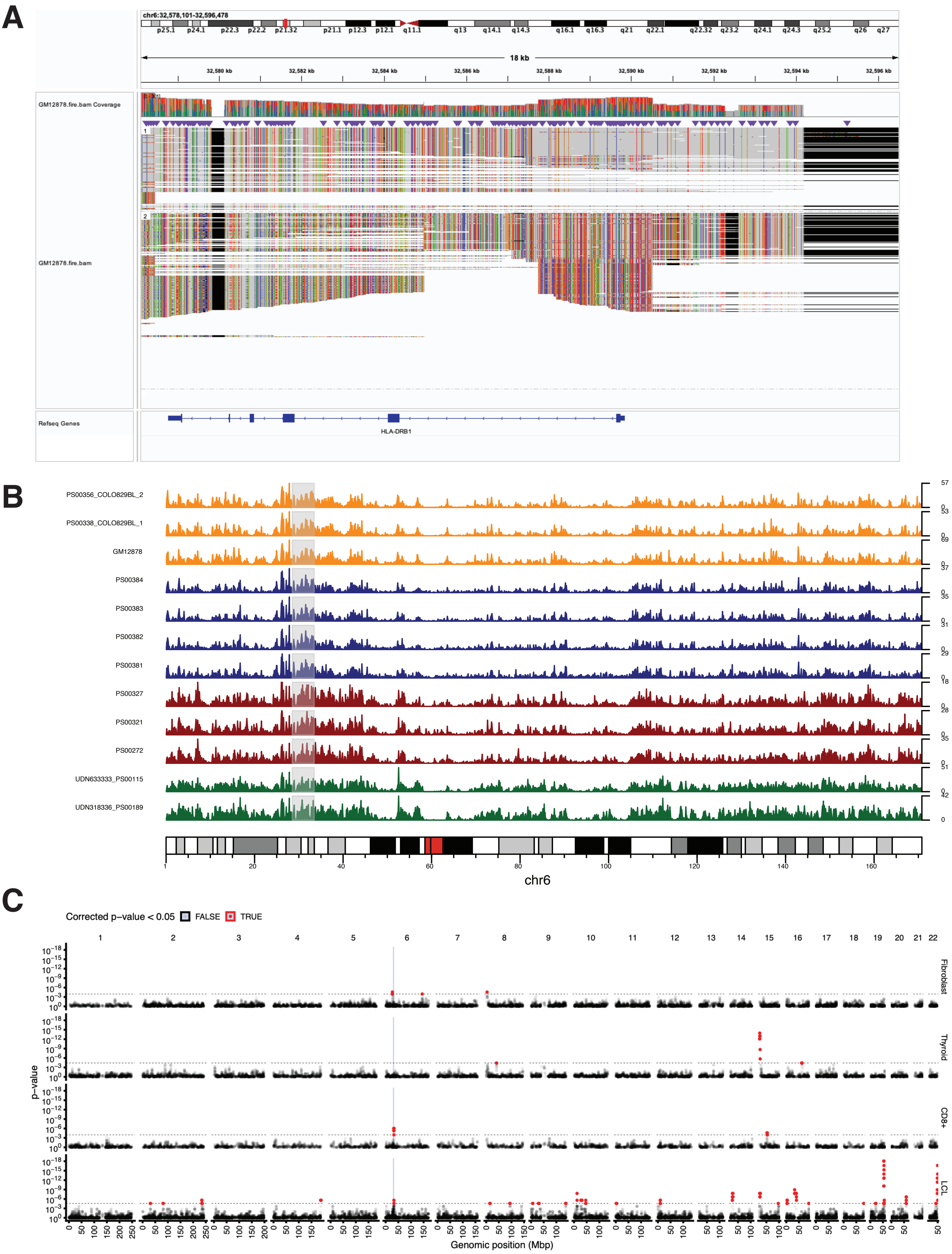

A
chr6:32,578,101-32,596,478
18 kb
GM12878.fire.bam Coverage
GM12878.fire.bam
Refseq Genes
HLA-DRB1
B
PS00356_COLO829BL_2
PS00338_COLO829BL_1
GM12878
PS00384
PS00383
PS00382
PS00381
PS00327
PS00321
PS00272
UDN633333_PS00115
UDN318336_PS00189
chr6
C
Corrected p-value < 0.05
FALSE
TRUE
p-value
Fibroblast
Thyroid
CD8+
LCL
Genomic position (Mbp)

### Figure S12

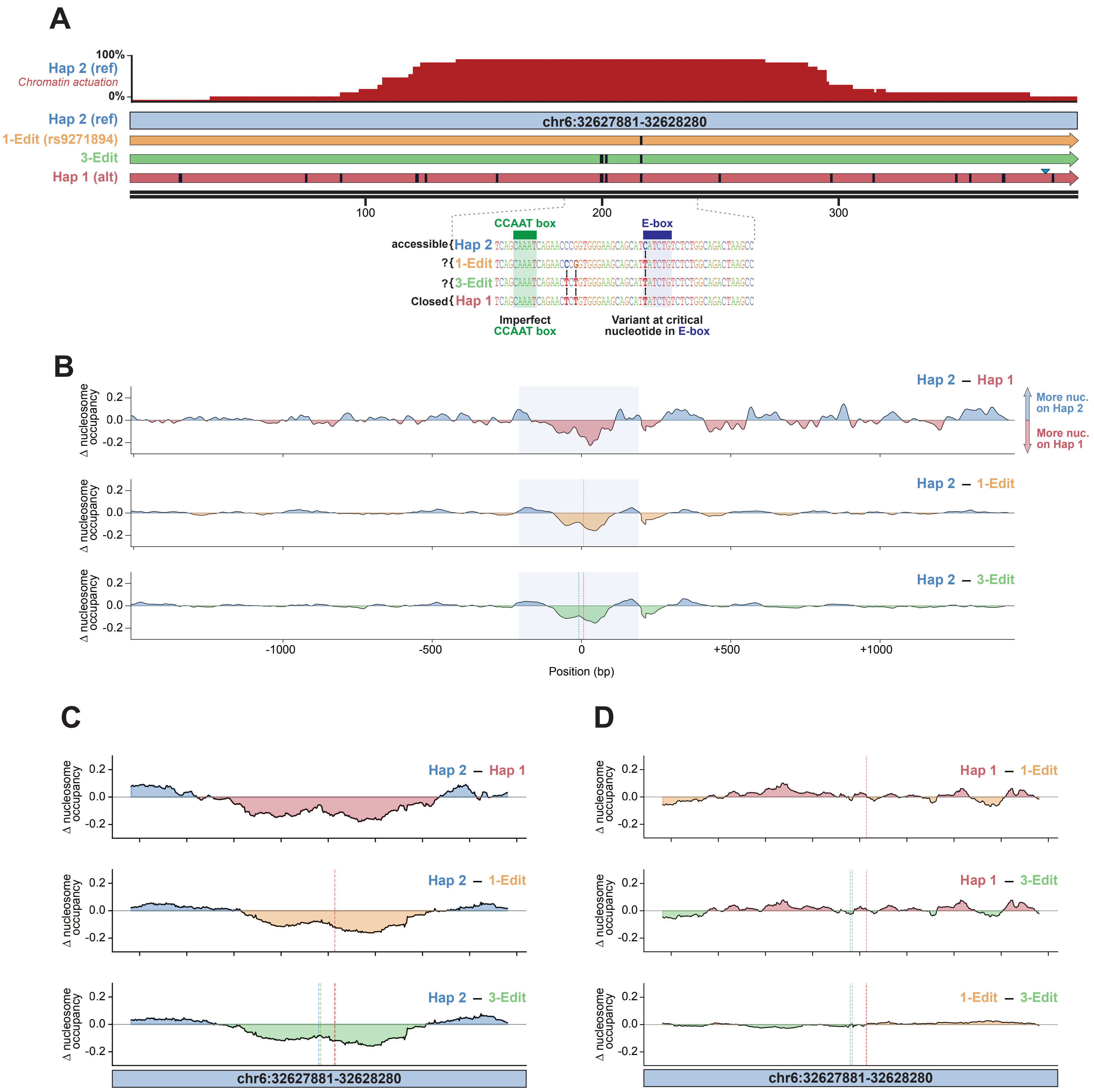

A
100%
0%
Hap 2 (ref)
Chromatin actuation
Hap 2 (ref)
chr6:32627881-32628280
1-Edit (rs9271894)
3-Edit
Hap 1 (alt)
100
200
300
CCAAT box
E-box
accessible{Hap 2 TCAGCAAATCAGAACCCGGTGGGAAGCAGCATCATCTGTCTCTGGCAGACTAAGCC
?{1-Edit TCAGCAAATCAGAACCCGGTGGGAAGCAGCATTATCTGTCTCTGGCAGACTAAGCC
?{3-Edit TCAGCAAATCAGAACTCTGTGGGAAGCAGCATTATCTGTCTCTGGCAGACTAAGCC
Closed{Hap 1 TCAGCAAATCAGAACTCTGTGGGAAGCAGCATTATCTGTCTCTGGCAGACTAAGCC
Imperfect CCAAT box
Variant at critical nucleotide in E-box
B
Δ nucleosome occupancy
0.2
0.0
-0.2
Hap 2 – Hap 1
More nuc. on Hap 2
More nuc. on Hap 1
Hap 2 – 1-Edit
Hap 2 – 3-Edit
-1000
-500
0
+500
+1000
Position (bp)
C
Hap 2 – Hap 1
Hap 2 – 1-Edit
Hap 2 – 3-Edit
chr6:32627881-32628280
D
Hap 1 – 1-Edit
Hap 1 – 3-Edit
1-Edit – 3-Edit
chr6:32627881-32628280

### Figure S13

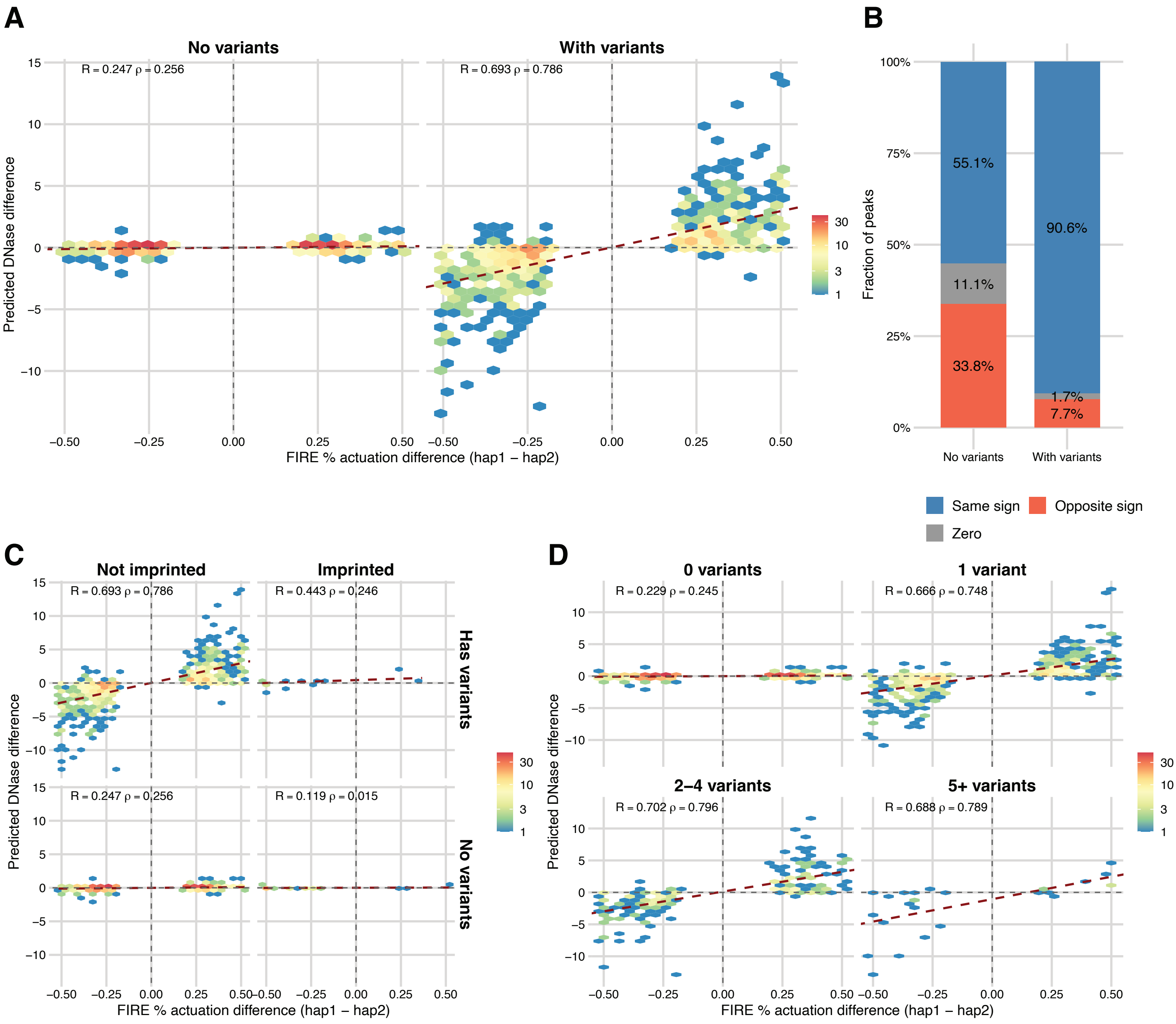
A
No variants
With variants
R = 0.247 ρ = 0.256
R = 0.693 ρ = 0.786
Predicted DNase difference
FIRE % actuation difference (hap1 – hap2)
B
Fraction of peaks
55.1%
11.1%
33.8%
90.6%
1.7%
7.7%
No variants
With variants
Same sign
Opposite sign
Zero
C
Not imprinted
Imprinted
Has variants
No variants
R = 0.693 ρ = 0.786
R = 0.443 ρ = 0.246
R = 0.247 ρ = 0.256
R = 0.119 ρ = 0.015
D
0 variants
1 variant
2–4 variants
5+ variants
R = 0.229 ρ = 0.245
R = 0.666 ρ = 0.748
R = 0.702 ρ = 0.796
R = 0.688 ρ = 0.789

### Figure S14

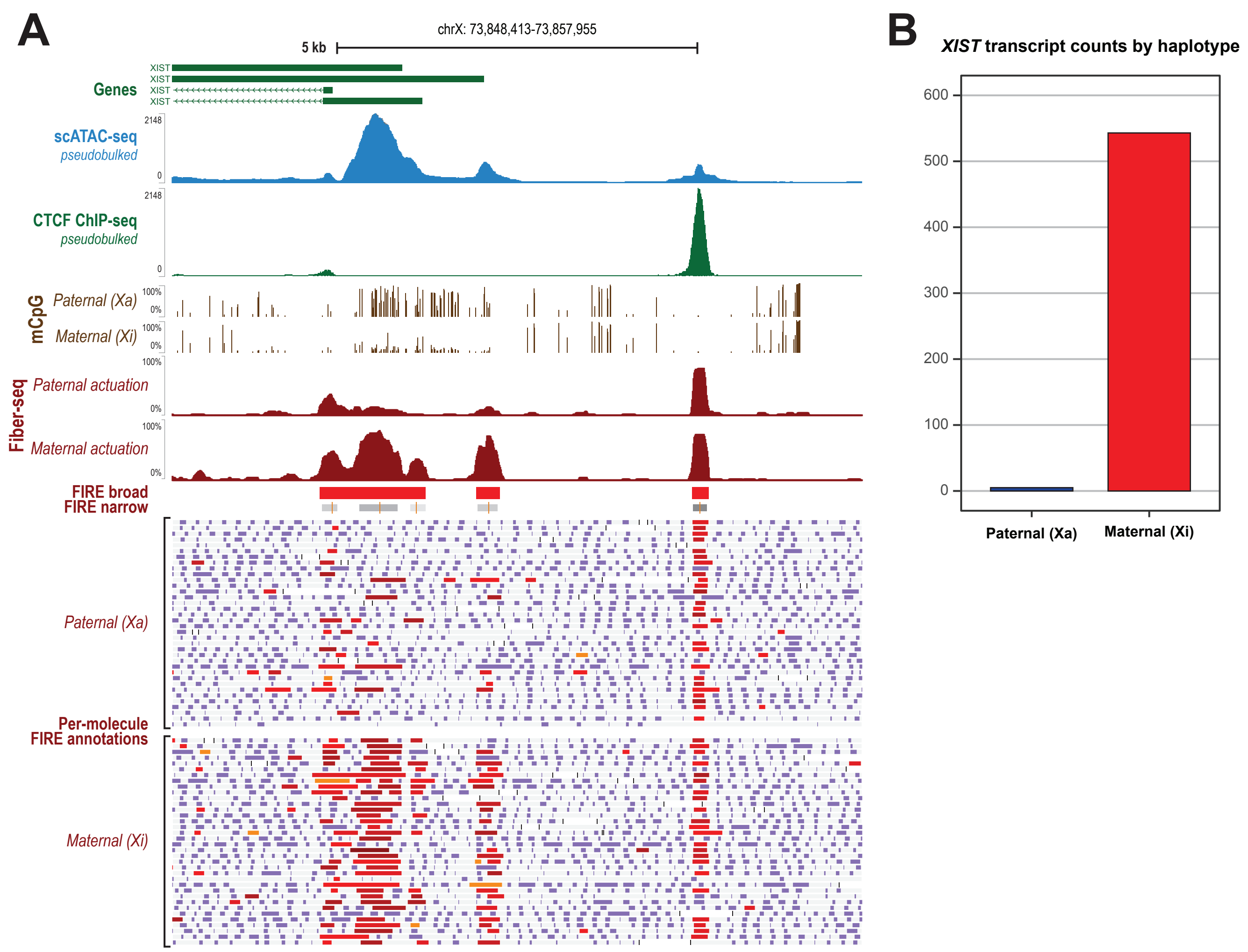
A
chrX: 73,848,413-73,857,955
5 kb
Genes
XIST
XIST
XIST
XIST
scATAC-seq
pseudobulked
2148
0
CTCF ChIP-seq
pseudobulked
2148
0
mCpG
Paternal (Xa)
Maternal (Xi)
100%
0%
Fiber-seq
Paternal actuation
Maternal actuation
FIRE broad
FIRE narrow
Per-molecule FIRE annotations
Paternal (Xa)
Maternal (Xi)
B
XIST transcript counts by haplotype
600
500
400
300
200
100
0
Paternal (Xa)
Maternal (Xi)

### Figure S15

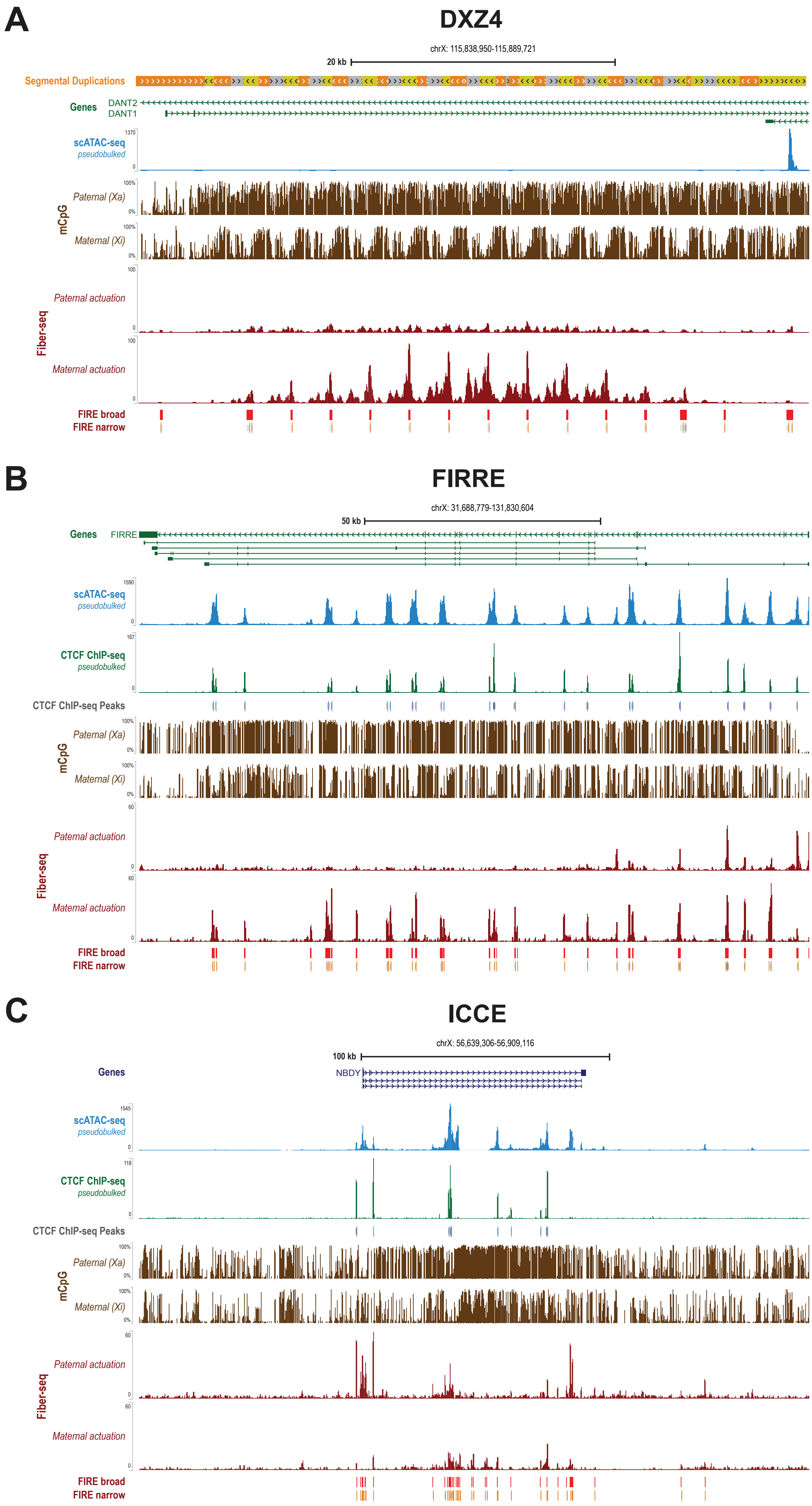
A
DXZ4
chrX: 115,838,950-115,889,721
20 kb
Segmental Duplications
Genes
DANT2
DANT1
scATAC-seq
pseudobulked
1370
0
mCpG
Paternal (Xa)
Maternal (Xi)
100%
0%
Fiber-seq
Paternal actuation
Maternal actuation
100
FIRE broad
FIRE narrow
B
FIRRE
chrX: 31,688,779-131,830,604
50 kb
Genes
FIRRE
scATAC-seq
pseudobulked
1590
CTCF ChIP-seq
pseudobulked
167
CTCF ChIP-seq Peaks
mCpG
Paternal (Xa)
Maternal (Xi)
Fiber-seq
Paternal actuation
Maternal actuation
60
FIRE broad
FIRE narrow
C
ICCE
chrX: 56,639,306-56,909,116
100 kb
Genes
NBDY
scATAC-seq
pseudobulked
1545
CTCF ChIP-seq
pseudobulked
118
CTCF ChIP-seq Peaks
mCpG
Paternal (Xa)
Maternal (Xi)
Fiber-seq
Paternal actuation
Maternal actuation
60
FIRE broad
FIRE narrow

### Figure S16

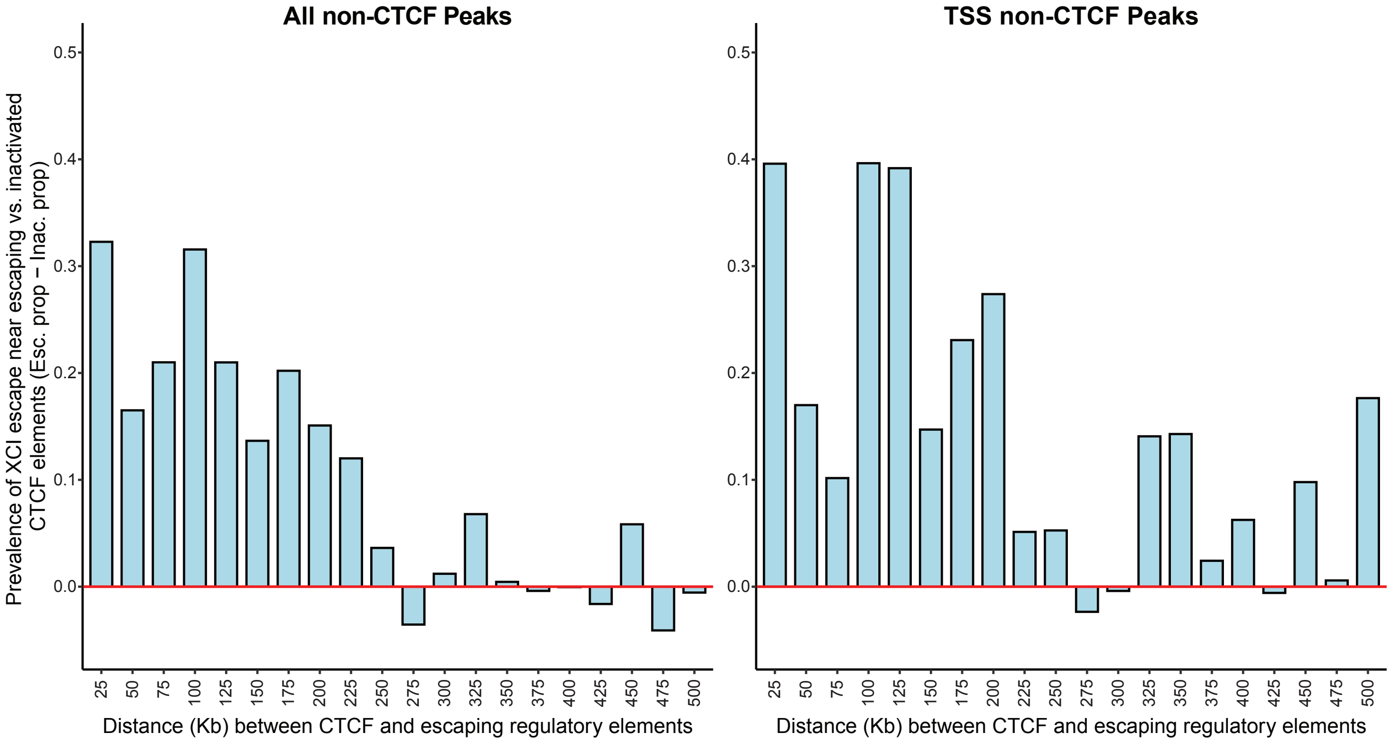

All non-CTCF Peaks
TSS non-CTCF Peaks
Prevalence of XCI escape near escaping vs. inactivated CTCF elements (Esc. prop − Inac. prop)
0.5
0.4
0.3
0.2
0.1
0.0
25
50
75
100
125
150
175
200
225
250
275
300
325
350
375
400
425
450
475
500
Distance (Kb) between CTCF and escaping regulatory elements
