## Supplemental Note for "Somatic epimutations cap genetic determinism in the human diploid chromatin epigenome"

### Supplemental Note: Description of the FIRE Methodology

#### Overview

Fiber-seq Inferred Regulatory Elements (FIRE) is a computational framework for identifying putative regulatory elements from Fiber-seq data at both the single-molecule and aggregate levels. The FIRE pipeline operates in two stages: (1) a semi-supervised machine learning classifier that assigns each methyltransferase-sensitive patch (MSP) on an individual Fiber-seq read a probability of being an actuated regulatory element, and (2) an aggregation and peak-calling procedure that combines single-molecule classifications across reads to identify regulatory element peaks genome-wide and estimate a false discovery rate (FDR) for each peak. Here, we provide a comprehensive description of each component of the FIRE methodology, including the nucleosome and MSP calling procedure, the construction of training data, the feature engineering, the semi-supervised learning framework, the estimated precision metric, the aggregate FIRE score, the FDR estimation, and the peak-calling algorithm.

#### Nucleosome and MSP calling

Fiber-seq relies on the non-specific N6-adenine methyltransferase Hia5 to stencil protein occupancy footprints along individual DNA molecules. Regions protected from Hia5 methylation correspond to nucleosome footprints, while regions accessible to methylation (methyltransferase-sensitive patches, or MSPs) correspond to either internucleosomal linker regions or actuated regulatory elements. Accurately delineating nucleosome positions is therefore a prerequisite for defining MSPs.

We use a heuristic nucleosome-calling algorithm implemented in fibertools-rs (ft add-nucleosomes) that is governed by three parameters: (1) a minimum nucleosome length *n* (default: 75 bp), (2) a minimum combined nucleosome length *c* (default: 100 bp), and (3) a minimum extension length *e* (default: 25 bp). The algorithm proceeds in four phases:

1. Identify all regions with no m6A events spanning at least *n* bases as candidate nucleosomes.
2. Identify all regions of at least *c* bases that contain only one internal m6A call (a putative false positive) as additional candidate nucleosomes.
3. Extend nucleosomes identified in phases 1 and 2 if spanning one additional m6A would add at least *e* bases of unmodified sequence to the nucleosome length.
4. Recursively apply step 3 until no nucleosome boundaries change.

After nucleosome calling, MSPs are operationally defined as all regions between adjacent nucleosome footprints on a given Fiber-seq read. This heuristic approach performs comparably to our previous hidden Markov model-based caller while being substantially faster, enabling genome-scale applications. The default parameters were selected to match the known size distribution of nucleosome core particles (~147 bp) while allowing for partial protection events.

##### Nucleosome calling is robust to parameter choice

To verify that FIRE scores are not sensitive to the specific nucleosome calling parameters, we systematically varied all three parameters across a 3×3 grid: minimum nucleosome length $n \in \{55, 75, 95 \text{ bp}\}$, minimum combined nucleosome length $c \in \{80, 100, 120 \text{ bp}\}$, and minimum extension length $e \in \{15, 25, 35 \text{ bp}\}$ (Fig. X).

The default parameters ($n$ = 75, $c$ = 100, $e$ = 25) produce a nucleosome length distribution with the highest yield and a mode near the canonical 147 bp, with smooth ~10 bp periodic sub-structure consistent with nucleosome breathing dynamics (Fig. S5A). This distribution is consistent across diverse sample types — including cell lines (A549, K562, THP-1), CHM13, HG002, primary tissues (liver, lung from donor ST001), and a tumor/normal pair (COLO829BL) — all showing a mode near 147 bp with preserved breathing periodicity (Fig. SXB).

Critically, FIRE scores are largely insensitive to the choice of nucleosome calling parameters. Comparing per-peak FIRE scores computed with the default parameters against each of the eight alternative parameter combinations, all pairwise Pearson correlations exceed $R$ = 0.99 (Fig. S5C). This demonstrates that the FIRE classifier learns features of the m6A signal within MSPs that are robust to reasonable variation in how MSP boundaries are defined, rather than relying on precise nucleosome boundary placement.

#### Training data construction

##### Fiber-seq training datasets

To ensure that the FIRE model generalizes across the range of experimental conditions encountered in practice, we generated 21 separate GM12878 Fiber-seq experiments, varying the global m6A methylation rate from 5.8% to 13.3% (Fig. S1A-C). This over 2.3-fold range in methylation rate captures the full spectrum of under- and over-methylated conditions that can arise from differences in enzyme concentration, incubation time, and cell permeability during the Fiber-seq protocol. By training on this diverse set of experiments, the final model learns to identify regulatory elements irrespective of the global methylation rate of a given sample.

##### Label assignment

Because no ground-truth single-molecule annotations of regulatory elements exist, we constructed training labels by leveraging the extensive short-read epigenomic data available for GM12878. Specifically:

**Mixed-positive labels.** MSPs that overlapped GM12878 DNase-seq hypersensitive sites (DHS; ENCODE accession ENCFF762CRQ), GM12878 CTCF ChIP-seq peaks (union of ENCFF356LIU and ENCFF960ZGP), or DHS hotspots were assigned a positive label. We randomly selected 100,000 peaks from the merged accessibility data (merging peaks within 147 bp, i.e., one nucleosome distance) and labeled MSPs with at least 25% reciprocal overlap as positive. Critically, these labels are “mixed-positive” rather than “clean-positive” because a regulatory element identified by bulk methods may show heterogeneous actuation at the single-molecule level: at any given locus, some individual fibers will have an actuated regulatory element while others will have an internucleosomal linker region. Thus, MSPs assigned a positive label based on genomic location represent a mixture of true regulatory elements and linker regions.

**Negative labels.** We identified 100,000 equally-sized genomic regions that did not overlap any DNase-seq or CTCF ChIP-seq peaks, excluding gaps in the hg38 reference assembly, segmental duplications, alternate contigs, and sex chromosomes. Negative regions were spatially shuffled using bedtools shuffle (-chrom -seed 42) to maintain chromosome structure while ensuring no overlap with positive regions.

**Sampling and independence.** From the 21 merged Fiber-seq datasets, we randomly sampled 10% of Fiber-seq reads overlapping the selected positive and negative regions. To enforce statistical independence between training examples and prevent the model from learning fiber-wide features rather than MSP-specific features, we selected at most one MSP per fiber per label class. MSPs shorter than 85 bp with a negative label were excluded from the training set to avoid trivially classified short linker regions that would inflate apparent model performance.

##### Train/test split

The labeled dataset was randomly split 80/20 (seed = 42) into training and held-out test sets. Both sets were independently class-balanced by downsampling the majority class to match the minority class, resulting in approximately equal numbers of mixed-positive and negative examples in each partition.

#### Feature engineering

The FIRE model operates on a set of features computed for each MSP that capture the local m6A modification pattern, independent of the underlying genomic sequence or position. This reference-free design is critical for enabling the model to generalize across cell types, complex genomic regions absent from the reference genome, and non-human organisms.

##### Feature categories

We compute the following categories of features for each MSP (Supplemental Table 3) and show an illustration in Figure S2B:

**MSP-level features.** These capture global properties of the MSP:

- *msp_len*: Length of the MSP in base pairs.
- *msp_m6a*: Number of m6A sites within the MSP.
- *msp_m6a_frac*: Fraction of adenine/thymine (AT) base pairs with m6A modification within the MSP.
- *msp_fc*: Log fold-change of the m6A fraction within the MSP relative to the overall m6A fraction of the entire Fiber-seq read. This normalization is essential for the model's ability to generalize across samples with different global methylation rates, as it transforms the absolute m6A signal into a relative enrichment score.
- *msp_len_times_m6a_fc*: The product of MSP length and the log fold-change of m6A. This composite feature captures the intuition that true regulatory elements tend to be both longer and more enriched for m6A than internucleosomal linkers. This feature serves as the initial discriminative feature in the first iteration of the semi-supervised learning procedure.

**Fiber-level reference features.** These provide context about the overall methylation state of the read:

- *fiber_m6a_count*: Total number of m6A sites across the entire Fiber-seq read.
- *fiber_m6a_frac*: Overall fraction of AT base pairs with m6A across the entire read.
- *ccs_passes*: Number of circular consensus sequencing (CCS) passes, which reflects sequencing quality.

**Windowed features (9 bins of 40 bp each).** To capture the spatial distribution of m6A modification within and around each MSP, we divide the MSP into nine 40 bp windows. Bins are zero-indexed from 0 to 8, so bin 4 is the center window, positioned at the MSP midpoint. For each window, we compute:

- *bin_X_m6a_count*: Number of m6A sites.
- *bin_X_m6a_frac*: Fraction of AT base pairs with m6A.
- *bin_X_m6a_fc*: Log fold-change of the window's m6A fraction relative to the whole-read m6A fraction.

These windowed features enable the model to learn the characteristic spatial patterns of m6A enrichment that distinguish regulatory elements from linker regions. For example, regulatory elements typically show broad, relatively uniform m6A enrichment across the MSP, whereas linker regions may show more heterogeneous or sparse modification patterns.

**Best/worst 100 bp window features.** We also identify the 100-bp sub-windows within the MSP with the highest ("best") and lowest ("worst") m6A signal. For each, we compute m6A count, m6A fraction, and m6A fold-change. These features capture the dynamic range of m6A modification within a single MSP, which can be informative for distinguishing regulatory elements (which tend to have consistently high m6A) from linker regions (which may have isolated m6A clusters).

##### Feature normalization for cross-sample generalization

A key design principle of the FIRE feature set is the use of fold-change normalization relative to the whole-read m6A level. Features such as *msp_fc*, *bin_X_m6a_fc*, and *msp_len_times_m6a_fc* express the local m6A enrichment relative to the fiber-level baseline. This normalization, in addition to the varied training data, helps ensure that the model can generalize to new samples with different global methylation rates without recalibration, because the signal is the *relative* enrichment of m6A at regulatory elements rather than only the absolute number of m6A modifications.

#### Semi-supervised learning with Mokapot

##### Motivation for a semi-supervised approach

The fundamental challenge in training the FIRE classifier is that our training labels are inherently noisy. MSPs labeled as positive based on overlap with DNase-seq peaks or CTCF ChIP-seq peaks represent a mixture of truly actuated regulatory elements and internucleosomal linker regions that happen to fall within bulk-defined peaks. This mixed-positive/clean-negative label structure precludes standard supervised learning, which assumes clean labels in both classes. Instead, we adopt a semi-supervised approach that can iteratively refine noisy positive labels while maintaining clean negative labels.

##### The Mokapot framework

We use the Mokapot framework, originally developed for peptide identification in mass spectrometry, which implements the semi-supervised strategy described by Käll et al. The key insight is that iterative retraining with progressively refined positive labels can yield a classifier that outperforms what would be achievable with the original noisy labels alone.

The semi-supervised training proceeds as follows:

1. **Initialization (Iteration 1).** Mokapot identifies the single feature that best discriminates between the mixed-positive and negative labels. In our implementation, this is *msp_len_times_m6a_fc*, which captures the composite signal of MSP length and relative m6A enrichment. A threshold is selected such that mixed-positive labels can be discriminated from negative labels at 95% estimated precision (defined below).

2. **XGBoost training.** The subset of mixed-positive labels above this initial threshold is used as positive training examples, combined with the full set of negative labels, to train an XGBoost gradient-boosted decision tree model with 5-fold cross-validation.

3. **Label refinement (Iterations 2-15).** The trained model's predictions are used to re-identify positive labels at 95% estimated precision. These refined positive labels — which now incorporate the model's learned representation — replace the original mixed-positive labels for the next round of training. This process repeats for up to 15 iterations.

4. **Convergence.** Training terminates when the number of positive identifications at 95% estimated precision in the validation set no longer increases, or after 15 iterations, whichever comes first.

#### XGBoost hyperparameters

The XGBoost classifier is configured with the following hyperparameters, selected via grid search with 5-fold cross-validation using ROC-AUC as the scoring metric:

| Parameter | Search space | Purpose |
|---|---|---|
| n_estimators | [200, 300] | Number of boosting rounds |
| max_depth | [9, 15] | Maximum tree depth |
| min_child_weight | [0.1%, 0.5%] of training set | Minimum instance weight in child node |
| colsample_bytree | [0.5, 1.0] | Fraction of features per tree |
| gamma | [1] | Minimum loss reduction for split |
| scale_pos_weight | auto | Ratio of negative to positive labels |

The scale_pos_weight parameter is automatically computed as the ratio of negative to positive examples after each round of label refinement, ensuring the model remains calibrated as the effective class balance shifts during semi-supervised training. The evaluation metric is AUC (area under the ROC curve), and a random seed of 42 is used throughout for reproducibility.

#### Model outputs

The training procedure produces three files: (1) the XGBoost model in binary format (FIRE.xgb.bin), (2) a JSON-format compatible with the Rust-based fibertools implementation (FIRE.gbdt.json), and (3) a confidence lookup table (FIRE.conf.json) that maps XGBoost prediction scores to estimated precision values. This lookup table enables rapid conversion from raw model scores to interpretable precision estimates during application.

#### Why FIRE can outperform its training labels

A natural question is how a classifier trained on DNase-seq and CTCF ChIP-seq labels can identify regulatory elements that these assays miss. The same question arises in proteomics, where semi-supervised methods such as Percolator and Mokapot routinely identify more correct peptide-spectrum matches than the initial database search scores alone (Käll et al., 2007, PMID: 17952086; Spivak et al., 2009, PMID: 19385687; Fondrie & Noble, 2021, PMID: 33596079). Three properties explain this.

Primarily, DNase and ChIP-seq peaks (the training labels) solve a different problem than single-molecule classification. DNase-seq and CTCF ChIP-seq peaks reflect aggregate signal across millions of cells: a region is called accessible when population-level signal exceeds a threshold. FIRE must instead classify individual MSPs on individual molecules. A strong DNase peak contains a mixture of actuated and non-actuated molecules; the bulk labels are therefore a useful but noisy proxy. This parallels proteomics, where database search scores perform well at ranking competing peptides within a single spectrum but lack calibration for absolute scoring across multiple spectra (Spivak et al., 2009, PMID: 19385687). By shifting the objective from predicting population-level peaks to predicting per-molecule regulatory elements, the aggregate FIRE signal may be more accurate than bulk ATAC or DNase.

Second, clean negatives and iterative refinement tolerate this label noise. MSPs from regions devoid of DNase or CTCF signal serve as reliable negatives, analogous to decoy peptide-spectrum matches generated from reversed or shuffled protein sequences (Käll et al., 2007, PMID: 17952086). Starting from these clean negatives and a small set of initial confident positives, the Mokapot framework (Fondrie & Noble, 2021, PMID: 33596079) iteratively retrains an XGBoost classifier, re-ranks all MSPs, and expands the confident set until convergence. This framework accommodates noise in the positive labels while relying on the high confidence of the negative set, allowing FIRE to distinguish between linker-regions (MSPs from regions devoid of DNase) and MSPs that are regulatory elements.

Third, Fiber-seq provides an orthogonal signal and directly measures the negative. DNase-seq and ATAC-seq depend on enzymatic cleavage and fragment capture, which introduces sequence-context biases, PCR artifacts, and mappability limitations. Fiber-seq directly measures m6A accessibility on native molecules and leverages long-read sequencing, thereby improving mappability and reducing sequence bias. Additionally, FIRE's reference-free features enable generalization to regulatory elements that short-read assays fail to capture (Fig. S5D). Critically, Fiber-seq sequences molecules regardless of accessibility, whereas ATAC-seq and DNase capture only cleaved fragments. This difference is important: when bulk assays show increased reads, it is unclear whether this reflects increased accessibility or whether sequence context and regulatory element size independently drive higher cleavage rates or capture efficiency. Because Fiber-seq is not subject to these issues, FIRE may more faithfully capture the underlying biology, improving its generalizability beyond the training labels.

##### <u>Inclusion of CTCF ChIP-seq peaks in training labels.</u>

CTCF ChIP-seq peaks were included among the mixed-positive training labels because visual inspection of the Fiber-seq data revealed that many CTCF-bound sites display clear m6A enrichment patterns on nearly every Fiber consistent with accessible chromatin, yet are not captured or have very low signal in DNase-seq or ATAC-seq data, likely because CTCF-bound sites tend to be narrow and are not always efficiently cleaved by DNase I or transposed by Tn5 (see the Figure below). Excluding these sites from training would have systematically biased the model against a biologically important class of regulatory elements.

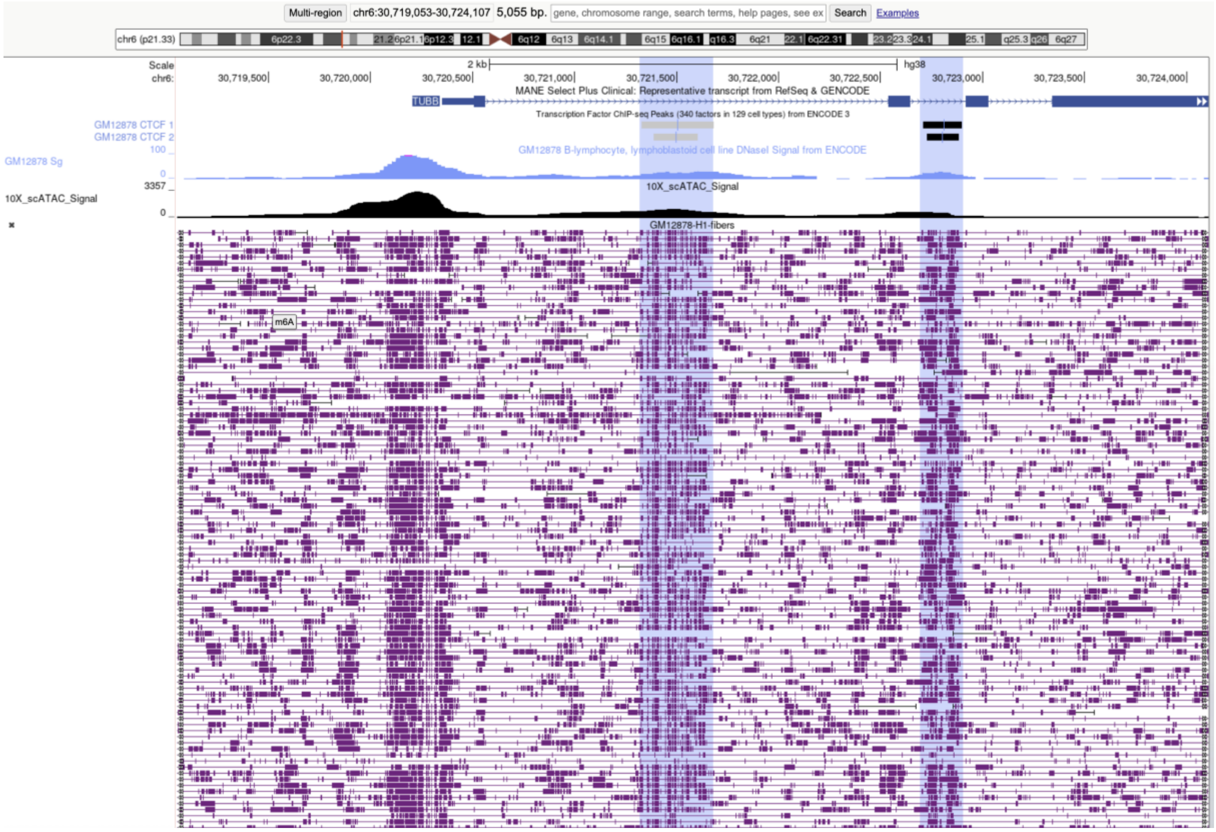


**Figure 1: CTCF-bound sites visible in Fiber-seq but with a weak signal in short-read accessibility assays.** UCSC Genome Browser view of a region on chr6 (chr6:30,719,053–30,724,107, hg38). From top to bottom: gene annotations (MANE Select Plus Clinical transcript), GM12878 CTCF ChIP-seq peaks (two replicates, ENCODE), GM12878 DNase-seq signal (ENCODE), 10X scATAC-seq signal, and individual GM12878 Fiber-seq reads with m6A modifications shown as purple marks. At CTCF-bound sites across this region, individual Fiber-seq reads show clear vertical stripes of m6A enrichment on nearly every fiber, indicative of constitutively accessible chromatin. Yet the DNase-seq and scATAC-seq signal at these same positions is weak relative to neighboring regulatory elements.

While this inclusion makes direct comparisons with DNase and ATAC somewhat asymmetric, the ability to incorporate such sites represents a specific advantage of Fiber-seq that cannot be achieved with the short-read assays alone. Two factors further mitigate the concern that FIRE is leaking CTCF information, thereby improving its correlation. First, the FIRE model operates entirely on m6A modification features computed from each Fiber-seq read and has no access to genomic sequence, motif content, or genomic position. It therefore cannot learn to recognize "CTCF sites" as a category — it can only learn the m6A signature of accessible chromatin regardless of which transcription factor is bound. This contrasts with sequence-to-function models, which could potentially predict CTCF binding by leaking sequence information. We demonstrate this generalizability in Figure S5D, where FIRE identifies REST/NRSF binding sites not detected by ATAC-seq, despite REST not being included in the training labels, indicating that the model learns generalizable features of accessible chromatin rather than memorizing label-specific patterns. Second, we tested the CTCF correlations in a new cell line and cell type (K562), precluding the possibility that a specific accessibility pattern of GM12878 CTCF sites was learned.

#### Empirical validation of the FIRE method.

The ability of FIRE to detect regulatory elements beyond its training labels is confirmed by several independent lines of evidence: (1) 77% of FIRE peaks overlap peaks from scATAC-seq, an orthogonal assay not used in training; (2) FIRE-specific peaks show significant enrichment for disease-associated GWAS variants comparable to peaks shared with DNase/ATAC (Fig. 2c); and (3) FIRE scores at CTCF sites are better correlated with CTCF ChIP-seq signal than ATAC-seq signal at those sites (R^2 = 0.41 vs. 0.17), despite the fact that both assays only measure chromatin accessibility.

#### Estimated precision of individual FIRE elements

Because we do not have access to clean-positive labels, we cannot compute a true precision for each FIRE element. Instead, we define an "estimated precision" (EP) metric using the balanced held-out test set of mixed-positive and negative labels (20% of the data).

For a given XGBoost score threshold, the estimated precision is defined as:

$$EP = 1 - \frac{FP + 1}{TMP}$$

where *TMP* is the number of identifications from mixed-positive labels with at least that XGBoost score, *FP* is the number of identifications from negative labels with at least that XGBoost score, and the pseudocount of 1 in the numerator prevents overly liberal precision estimates when the number of identifications is small.

Both the semi-supervised training and the final classification use the same 95% estimated precision threshold (corresponding to a 5% FDR on the training labels). During training, this threshold is used to iteratively refine positive labels, ensuring that only high-confidence positive examples are used to retrain the model. For the final classification of MSPs as FIRE elements, MSPs with an estimated precision at or above this threshold are classified as FIRE elements. In practice, estimated precision values are stored as unsigned 8-bit integers (u8) in the output BAM file, so the effective threshold is 243/255 ≈ 94.9%. Users can adjust this threshold depending on their application.

#### Aggregate FIRE score calculation

After classifying individual MSPs as FIRE elements or linker regions, we aggregate the per-molecule classifications across all Fiber-seq reads covering each genomic position to produce a continuous score reflecting the evidence for regulatory element presence.

The aggregate FIRE score $S_g$ at genomic position $g$ is defined as:

$$S_g = \frac{-50}{R_g} \sum_{i=1}^{C_g} \log_{10}(1 - \min(EP_i, 0.99))$$

where $C_g$ is the number of FIRE elements overlapping position $g$, $R_g$ is the total number of Fiber-seq reads covering position $g$, and $EP_i$ is the estimated precision of the $i$-th FIRE element at position $g$.

This formulation has several desirable properties. First, the division by $R_g$ normalizes for sequencing coverage, ensuring that the score reflects the fraction of fibers with regulatory element evidence rather than the absolute number of supporting reads. Second, the logarithmic transformation of $(1 - EP_i)$ places greater weight on high-confidence FIRE elements: an element with 99% estimated precision contributes $-\log_{10}(0.01) = 2$ to the sum, while an element with 90% estimated precision contributes $-\log_{10}(0.1) = 1$. Third, capping $EP_i$ at 0.99

and multiplying by -50 bounds the scores to the range [0, 100]. Positions covered by fewer than 4 FIRE elements ($C_g < 4$) are not scored and are assigned a value of -1.

In addition to the aggregate FIRE score, we compute the "percent actuation" at each peak, defined as $C_g/R_g$ — the fraction of all Fiber-seq reads at a position that carry a FIRE element. This metric provides an intuitive, biologically interpretable measure of chromatin accessibility: a percent actuation of 50% means that half of the sequenced chromatin fibers at that locus have an accessible regulatory element. Unlike the arbitrary signal units of DNase-seq or ATAC-seq, percent actuation has a direct physical interpretation in terms of the fraction of molecules with accessible chromatin.

#### FDR estimation via read shuffling

To calibrate the statistical significance of aggregate FIRE scores, we estimate a false discovery rate using a read-shuffling null model.

**Generating the null distribution.** We shuffle the genomic location of each Fiber-seq read by randomly selecting a new start position within the same chromosome, preserving the read's internal structure (MSP positions, FIRE classifications, and all features) while randomizing its genomic context (bedtools shuffle -chrom -excl). Reads originating from regions with unreliable coverage — defined as regions where sequencing coverage deviates from the median by more than 5 standard deviations, where the standard deviation is derived from a Poisson model (i.e., the square root of the mean coverage) — are excluded from shuffling, and shuffled reads are not placed into unreliable coverage regions.

**Computing FDR.** After shuffling, we recompute aggregate FIRE scores across the genome using the shuffled read positions. The FDR associated with a given FIRE score threshold $S_g$ is then estimated as:

$$FDR(S_g) = \frac{\text{number of bases with shuffled FIRE score} \geq S_g}{\text{number of bases with observed FIRE score} \geq S_g}$$

This approach generates a monotonic mapping from FIRE score thresholds to FDR values, enabling us to assign an FDR to every scored position in the genome.

#### Peak calling

FIRE peaks are identified through a multi-step procedure that combines FDR-based thresholding with element-aware merging.

**Step 1: Local maxima identification.** We identify all positions that represent local maxima of the aggregate FIRE score within a 200 bp rolling window. A position qualifies as a local maximum if its FIRE score equals the maximum score within the window. When multiple adjacent positions share the same maximum score, the midpoint of the plateau is selected.

**Step 2: FDR filtering.** Local maxima are filtered to retain only those with an FDR below the user-specified threshold (default: 5%) and a minimum percent actuation of 10% ($C_g/R_g \geq 0.10$).

**Step 3: Peak merging.** Adjacent peaks are merged through a multi-stage iterative algorithm:

- *Stage 1:* Peaks with 90% or greater reciprocal genomic overlap are merged, retaining the higher-scoring local maximum.
- *Stage 2:* Peaks sharing 50% or more of their underlying FIRE elements are merged.

- *Stage 3:* A final round of 90% reciprocal overlap merging is applied.

This multi-stage approach ensures that closely spaced regulatory elements are appropriately merged when they share substantial molecular evidence, while distinct elements in close genomic proximity are maintained as separate peaks.

**Step 4: Boundary definition.** Peak boundaries are defined by the median start and end positions of the underlying single-molecule FIRE elements, rather than by arbitrary score thresholds. This provides near-nucleotide-resolution boundaries that directly reflect the physical extent of chromatin accessibility observed across individual molecules.

**Wide peaks.** We additionally report "wide peaks" by taking the union of all FIRE peaks and all genomic regions with FDR below the threshold, then merging regions within 147 bp (one nucleosome length) of each other. These wide peaks capture broader domains of regulatory activity that may encompass multiple closely spaced elements.
